# To hybridise or not to hybridise? Systematic review and meta-analysis of woodpeckers

**DOI:** 10.1101/2025.01.20.633118

**Authors:** Antonii Bakai, Jérôme Fuchs, Gerard Gorman, Dominika Sajdak, Łukasz Kajtoch

## Abstract

Hybridisation is a common phenomenon among birds in general. Woodpeckers (Picidae) are no exception, as approximately 20% of species are known to hybridise and for many others interspecific mating is suspected. However, the mechanisms and consequences (phenotypic and genetic) of hybridisation are known for only a fraction of woodpecker species. Here, we conduct a systematic review on the literature that deals with hybridisation in woodpeckers and use a meta-analytical approach to examine the available geographical and genetic data. According to available published data, the majority of woodpeckers that hybridise inhabit the Neotropics, followed by the Nearctic and the Palearctic. Hybridisation appears to be less common in the Afrotropic and Oriental regions. As expected, genetic distances are substantially lower between hybridising species pairs than between non hybridising congenerics. This implies that hybridisation is typical for “young” (sister) pairs of species, that typically have similarities in their respective reproductive biology, ecology and ethology, and thus had less time for genetic incompatibilities to build up. These similarities also explain the difficulties experienced in studies on hybrid woodpeckers, but these could be overcome by the use of modern techniques (remote sensing and/or citizen science combining with AI identification and genomic identification). As hybridisation plays a role in woodpecker evolution and ecology it should be considered when conservation policies for rare species are drafted.

## INTRODUCTION

Hybridisation has played a significant role in the history of species formation (Barton 2001). Mixed species pairs initially appear to contradict the classic “biological concept” of species, which assumes the existence of barriers that prevent a mixing of gene pools among organisms. Yet, like many human-invented concepts, numerous exceptions exist because the evolution of reproductive isolation is a continuous process (Zachos 2016) irrespective of species life history traits or ecology (Roux et al. 2016). There are many examples of microorganisms, and also plants and fungi, which are known or suspected to be descendants of interspecific breeding, or which simply have genes inherited from various ancestors (Arnold 1992). Among animals, hybrids have been often presumed to be rare, inviable and infertile (Pala & Coelho 2005), but owing to the use of more sophisticated methods of detection and analyses of integrative data, hybrids have been found to be not as rare in speciation or population evolution (Schwenk et al. 2008).

Hybridisation among closely related phylogenetic species is a fairly common phenomenon in birds (Grant & Grant 1992, Ottenburghs 2023). This could be in part explained by the fact that most extant species are relatively recent, having been formed during the Pliocene and Pleistocene climatic oscillations during which the evolution of pre-zygotic isolating mechanisms may have not had enough time to form. Populations, lineages and species formed during that time period had several opportunities to be in secondary contact and interbreed with populations from other refugia during more favourable periods, which could have allowed new allelic combinations. Consequently, many hybrid zones among species for which ranges overlap have been described (Edwards et al. 2005; Pereira et al. 2011). The probability of hybridisation is further reinforced because in many closely related species, similar ecological and morphological traits enabled hybrids to continue to occur (Roux et al. 2016).

In many cases, the key point in hybridisation is mate abundance; if one species is rare (for example, on the edge of its range), it may be forced to look for mates from other species. There are many examples of such hybrids, and they are usually associated with certain hybridisation zones or “hot spots” (Ottenburghs 2018).

The Picidae family (wrynecks, piculets, woodpeckers) is relatively homogeneous morphologically (Prum et al. 2015, Shakya et al. 2017). Most woodpeckers are sedentary, rather poor dispersers and are highly territorial, and this can result in strong barriers to interspecific breeding because opportunities to mate with another related species are thus reduced. Among the approximately 240 woodpecker species (Gorman 2014), 88 are known to be able to interspecifically breed (Ottenburghs & Nicolaï 2024) but reliable records exist for only 46. There are 68 known hybrid combinations, but only 37 have been reliably confirmed and documented. Ottenburghs & Nicolaï (2024) focused on hybridisation versus mimicry in woodpeckers and concluded that nine million years is the threshold between these two phenomena. Hybridisation is typical for “young” pairs of species and mimicry (convergent evolution) is seen in “older” species pairs. Ottenburghs & Nicolaï (2024) is an important contribution to our general understanding of hybridisation in woodpeckers, especially in relation to plumage mimicry, but does not cover all biological, ecological and behavioural details. Neither does that review consider phylogenetic and biogeographic relationships among hybridising species, particularly if hybrids are more frequent between sister or non-sister species, and if the extend of overlapped ranges (parapatric vs sympatric) is a crucial factor in hybrid formation or not.

Here, we systematically earched the literature to examine and describe the available information on this subject. We also used genetic data to verify differences in divergence between hybridising and non-hybridising congeneric species and to map hybridisation events along phylogenetic relatedness. In particular, our aims were to: i) summarise the state of knowledge on hybridisation and introgression among the Picidae; ii) outline hybridisation among selected complexes, taking into account their morphological, ecological, ethological and geographic features; iii) compare genetic distances among hybridising and non-hybridising species in order to resolve if sibling species are really prone to hybridise and whether hybridisation is more common in some specific clades; iv) map hybridisation events along phylogenetic lines in order to understand if interspecific breeding originated in a random fashion or if it was associated with some events (geological, climatic or environmental). We also intended to estimate what any possible consequences of hybridisation and introgression on the specification of woodpeckers could be, as well as on their ecology and conservation. Moreover, we emphasised gaps in knowledge, listing less studied genera, populations from particular regions or habitats, and highlighted the prospects for future studies on hybridisation in woodpeckers using modern tools.

## METHODS

### Literature search

To review the current state of knowledge on hybridisation in woodpeckers, a literature search, using the Web of Science and Scopus open electronic databases, was conducted in December 2023 according to Preferred Reporting Items for Systematic reviews and Meta-Analyses (PRISMA; https://www.prisma-statement.org/). The following keys were used: (Picidae OR woodpeckers) AND (hybrid OR hybridisation OR “mixed pair”). The downloaded metadata was arranged in a single sheet holding the information on the authors, title, journal, volume, first and last page numbers, year of publication, and abstract (when available). Each text was briefly evaluated on the basis of its title and abstract to eliminate publications that were not connected to theme (e. g. hybridisation in non-woodpecker species). Topics which were not directly connected to our study were excluded from the analysis and the reason for the rejection was noted. All duplicates within databases and between databases were later removed. Publications for which we were unable to obtain the full text (e.g., old or internal publications), were also excluded from the analysis.

### Descriptive systematic review

After the publications had been filtered, the remaining were examined in detail in order to include more information in the sheet. The texts were screened for: 1) hybridising genera and species; 2) continent, country and habitat type where hybridisation was confirmed; 3) methods and views mentioned in study (e.g., ecological, behavioural); 4) reliability of the hybridisation event reported (mixed pairs, presumed or confirmed). The obtained data was then used to classify the examined scientific publications and to provide a descriptive summary on each category.

For all known hybridisation cases, a brief summary of the genera and species was provided using added literature.

### Meta-analysis

#### Hybridisation geography

For each pair of hybridising species, based on Ottenburghs & Nicolaï (2024) their respective distributions were assigned using data from Birds of the World (Billerman et al. 2022). The species were assigned to continent and zoogeographic realm, following Cox et al. (2001) (we do not consider newer classification e.g., by Holt et al. (2013) as woodpeckers as associated with woodlands and some new realms are inhabited by a few or no species). Moreover, we assigned woodpecker species to major biomes on the Earth considering four wooded types: boreal, temperate, subtropical (Mediterranean-like) and tropical. Treeless arctic, mountain and dry deserts as well as grasslands and shrublands were not considered.

#### Range overlap

For each species pair, their hybridising ranges, reliable and uncertain, based on Ottenburghs & Nicolaï (2024), were verified using Birds of the World (Billerman et al. 2022) and BirdLife International and Handbook of the Birds of the World (2018). Three categories were assigned: sympatric (if at least part of the ranges overlapped), parapatric (if ranges were adjoining but did not substantially overlap), and allopatric (no overlap). The frequencies of sympatric and parapatric pairs of both hybridising and non-hybridising woodpeckers was compared using a Chi2 test.

#### Karyotypes and genomes

Data on the genomic structure of woodpecker genera with known hybrids (both classic karyotypes and genomes) were retrieved from the literature by searching for articles using keys (genus name AND karyotype) and (genus name AND genome). We did not use PRISMA guidelines here owing to the relatively small number of studies found and because this was not the primary goal of this review. Additionally, we checked GenBank resources to retrieve unpublished genomes of woodpeckers. We used identified information for a brief characterisation of woodpecker karyotypes and genomes in order to link them with hybridisation events in the discussion.

#### Phylogenetic relatedness

Genetic and phylogenetic data on woodpeckers from Shakya et al. (2017) were used. For all species with data, three genetic markers were downloaded from GenBank: ATP synthase 6 (hereafter ATP6); autosomal transcription growth factor β 2 intron 5 (hereafter TGFb2) and Z-linked muscle skeletal receptor tyrosine kinase intron 4 (hereafter MUSK). Accession numbers were taken from Table S1 in Shakya et al. (2017). For each marker, a genetic distance was calculated for species pairs in genera that include at least one reliable or uncertain pair of hybridising species according to Ottenburghs & Nicolaï (2024). Three datasets were obtained per marker for: hybridising pairs (reliable), hybridising pairs (uncertain), and non-hybridising (only intrageneric). Median, average, span and standard deviation values were calculated for each of these groups (separately for each marker). Differences between the three defined groups were compared using an analysis of variance and the data were visualised using R 4.0.4 (R Core Team 2024).

The phylogenetic tree from Shakya et al. (2017) was used to track hybridising pairs across the current estimate of the woodpecker phylogeny. First, each pair (including those uncertain according to Ottenburghs & Nicolaï 2024), were assigned as ‘sister’ or not (as ‘sister’ species were considered as being those that are most closely related). Second, for selected genera (that included hybridising species according to Ottenburghs & Nicolaï (2024), subsets of the tree from Shakya et al. (2017) were used to visualise the distribution of hybridising pairs on the topology. In addition, genetic differences for both hybridising species and non-hybridising, were obtained from Shakya et al. (2017) and used in the discussion on evolutionary and biogeographic hybridisation in woodpeckers.

## RESULTS

### Literature search summary

Out of the 290 original publications in the database, 200 were rejected (Figure 1). Of those, 106 texts were rejected because of unrelated content. That is, they did not include information relevant to this review, such as studies on non-woodpeckers. For the 32 publications (mostly papers published during the 1950-2000 time period in local journals), we were unable to obtain abstracts in order to check the content. A further 62 publications were duplicates, mainly owing to repeated searches in two different databases by the same key (of course, some publications were found in both the Web of Science and Scopus).

**Figure 1.**
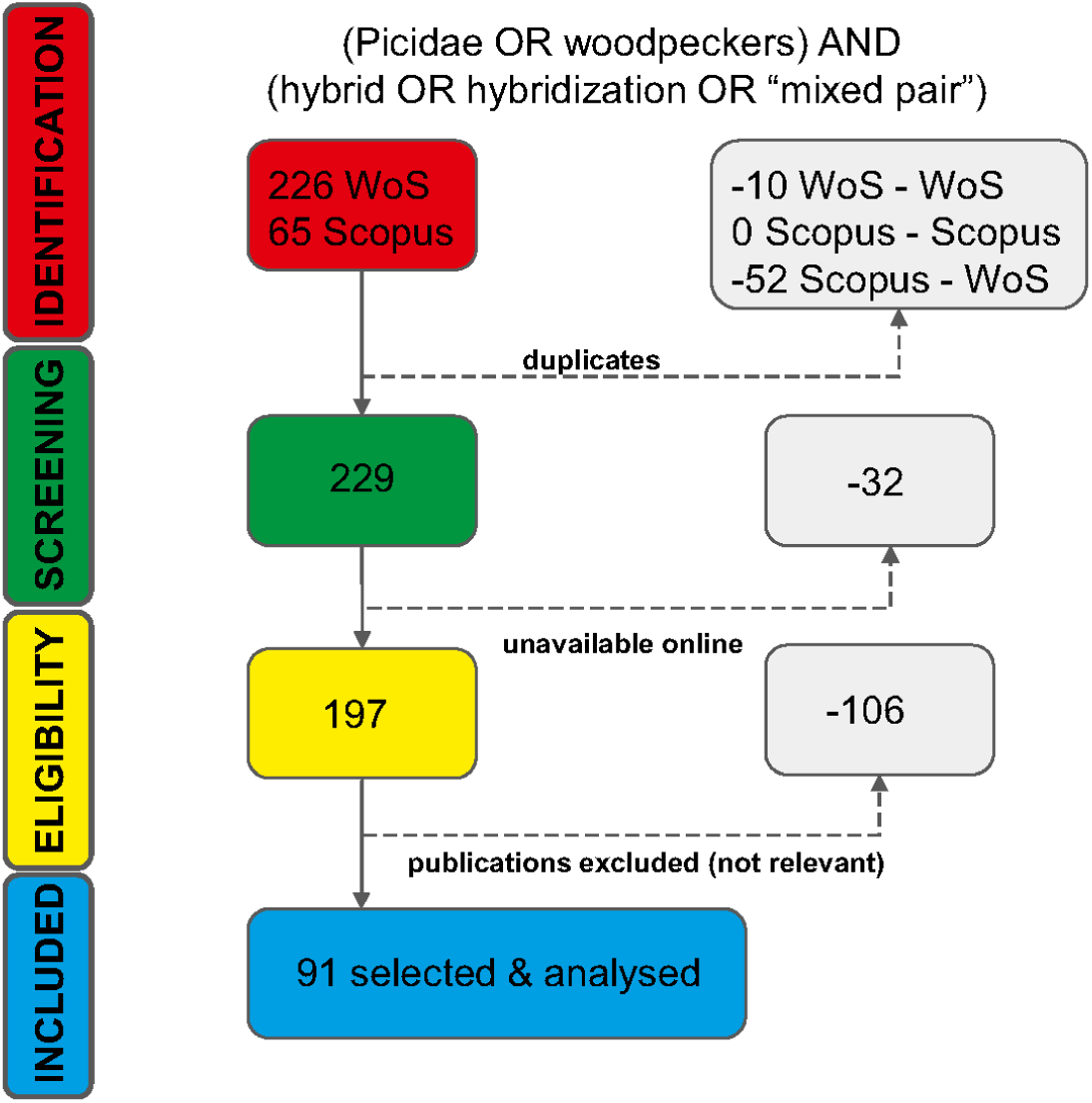
Literature search workflow according to Preferred Reporting Items for Systematic reviews and Meta-Analyses (PRISMA).

Of the 90 publications selected (Table S1), 62 described hybrids or mixed pairs in the field and 39 were based on data from museum specimens; 21 included both sources. Hybrids or mixed pairs were mostly found in forests, wooded grasslands or other natural habitats (61), out of which many were in riverine ecosystems (14), especially for the genera *Picus* and *Colaptes*. The second major category was found in anthropogenic wooded areas (22), especially in the cases of *Dendrocopos* (*D. syriacus* x *D. major*) and *Picus* (*P. canus* x *P. viridis*).

Mixed pairs were reported in 21 publications, and hybrids in 83 publications (Table S1), and from these hybrids were confirmed in 67 cases and in 17 merely presumed.

Most of the literature deals with woodpeckers hybridising in the Palearctic (13 Poland, 1 Germany, 1 France, 1 Denmark, 1 Russia) and the Nearctic (55 USA, 26 Canada) bioregions, less so in the Neotropics (3 Mexico, 4 Brazil & Bolivia, 2 Paraguay). A few publications describe hybridisation in the sub-Saharan Africa (1 Tanzania, 1 Congo) and Oriental regions (1 Laos & Thailand, 2 Sri Lanka) (Table S1).

The works we found began with a note by Baldwin on hybridisation between Northern Flicker *Colaptes auratus lutens* and Red-shafted Flicker *Colaptes cafer collaris* in 1910, but more often texts mentioning woodpecker hybridisation started to appear from 1969 on, with most published in the 1980s (based on morphology) and in last two decades (when molecular DNA analysis became more common) (Figure 2). The highest number of studies have been conducted on *Colaptes* (34 publications), followed by *Sphyrapicus* (19), *Dendrocopos* (10) and *Picus* (8). Publications concerning other hybridising genera do not exceed five texts per genus.

**Figure 2.**
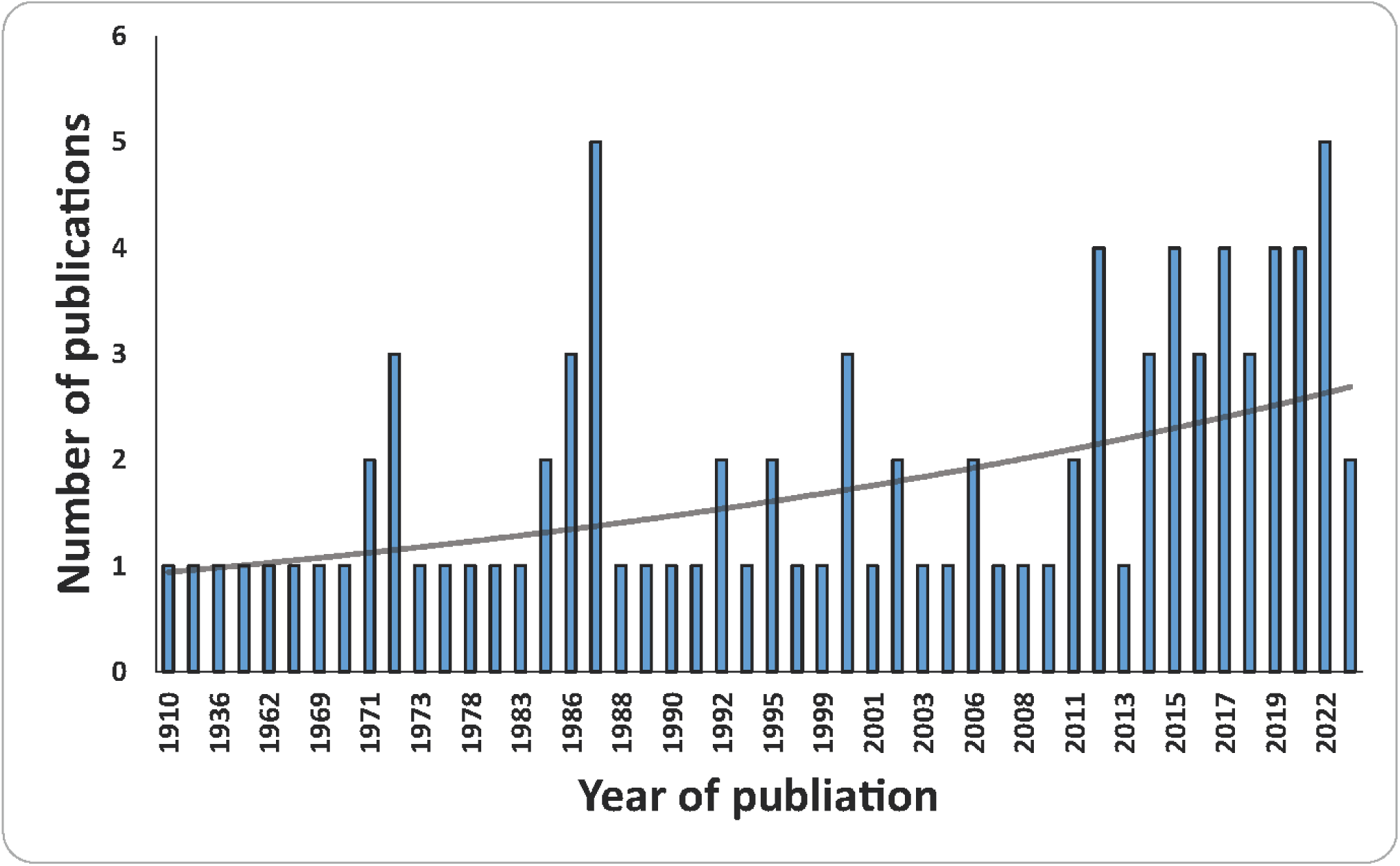
Progress in studies on hybridisation among woodpeckers worldwide.

Most of the studies examined dealt with the ecology of the hybridising species (22) and genetics (27) (Table S1). Information on breeding biology was present in 20 studies, on the four most often studied genera: *Colaptes, Dendrocopos, Sphyrapicus* and *Picus*. Behaviour was described in 16 publications and involved the same four genera.

### Geography of woodpecker hybridisation

The majority of species pairs known to hybridise (excluding uncertain cases listed by Ottenburghs & Nicolaï 2024) occur in North America – 16 pairs (Table 1), followed by South America – 12 pairs, Asia – 7 pairs, Africa – 4 pairs, and Europe – 4 pairs (some pairs are present both in Asia and Europe). The ratio of hybridising species differs significantly when compared to the total of all woodpecker species on any given continent (χ^2^=9.63, p=0.04) (Figure 3).

**Table 1.** Geographic and phylogenetic characteristics of hybridising pairs of woodpeckers.

| species 1 | species 2 | continent | zoogeographic realm | biome | range overlap | sister taxa |
| --- | --- | --- | --- | --- | --- | --- |
| *Campephilus leucopogon* | *Campephilus melanoleucos* | S America | Neotropical | tropical | sympatric | no |
| *Campethera cailliautii* | *Campethera maculosa* | Africa | Afrotropical | subtropical, tropical | sympatric | yes |
| *Celeus elegans* | *Celeus lugubris* | S America | Neotropical | tropical | sympatric | yes |
| *Colaptes chrysoides* | *Colaptes auratus* | N America | Nearctic, Neotropical | boreal, temperate, subtropical | parapatric | yes |
| *Dendrocopos assimilis* | *Dendrocopos syriacus* | Asia | Paleartic | subtropical | parapatric | yes |
| *Dendrocopos leucopterus* | *Dendrocopos major* | Asia | Paleartic | temperate, subtropical | parapatric | yes |
| *Dendrocopos major* | *Dendrocopos syriacus* | Asia, Europe | Paleartic | temperate, subtropical | sympatric | no |
| *Dendrocopos major* | *Dendrocopos leucotos* | Asia, Europe | Paleartic | boreal, temperate | sympatric | no |
| *Dendropicos goertae* | *Dendropicos griseocephalus* | Africa | Afrotropical | tropical | sympatric | no |
| *Dendropicos goertae* | *Dendropicos spodocephalus* | Africa | Afrotropical | subtropical, tropical | sympatric | no |
| *Dendropicos griseocephalus* | *Dendropicos spodocephalus* | Africa | Afrotropical | subtropical, tropical | parapatric | yes |
| *Dryobates nuttallii* | *Dryobates pubescens* | N America | Nearctic | boreal, temperate, subtropical | sympatric | no |
| *Dryobates scalaris* | *Dryobates nuttallii* | N America | Nearctic, Neotropical | subtropical | sympatric | yes |
| *Dryobates scalaris* | *Dryobates pubescens* | N America | Nearctic, Neotropical | boreal, temperate, subtropical | parapatric | no |
| *Dryobates scalaris* | *Leuconotopicus villosus* | N America | Nearctic, Neotropical | boreal, temperate, subtropical | sympatric | no |
| *Dryocopus lineatus* | *Dryocopus schultzei* | N America, S America | Nearctic, Neotropical | subtropical, tropical | parapatric | yes |
| *Leuconotopicus villosus* | *Dryobates pubescens* | N America | Nearctic, Neotropical | boreal, temperate, subtropical | sympatric | no |
| *Gecinulus grantia* | *Gecinulus viridis* | Asia | Oriental | tropical | parapatric | yes |
| *Melanerpes aurifrons* | *Melanerpes uropygialis* | N America | Nearctic | tropical | sympatric | no |
| *Melanerpes aurifrons* | *Melanerpes hoffmannii* | N America | Nearctic, Neotropical | tropical | sympatric | no |
| *Melanerpes carolinus* | *Melanerpes aurifrons* | N America | Nearctic | tropical | sympatric | yes |
| *Melanerpes hoffmannii* | *Melanerpes santacruzi* | N America | Nearctic, Neotropical | tropical | sympatric | no |
| *Melanerpes hoffmannii* | *Melanerpes rubricapillus* | N America, S America | Nearctic, Neotropical | tropical | sympatric | yes |
| *Picumnus albosquamatus* | *Picumnus cirratus* | S America | Neotropical | tropical | sympatric | no |
| *Picumnus albosquamatus* | *Picumnus dorbygianus* | S America | Neotropical | tropical | parapatric | no |
| *Picumnus albosquamatus* | *Picumnus temminckii* | S America | Neotropical | tropical | sympatric | no |
| *Picumnus castelnau* | *Picumnus subtilis* | S America | Neotropical | tropical | parapatric | yes |
| *Picumnus cirratus* | *Picumnus dorbygianus* | S America | Neotropical | tropical | parapatric | yes |
| *Picumnus cirratus* | *Picumnus varzeae* | S America | Neotropical | tropical | sympatric | no |
| *Picumnus temminckii* | *Picumnus cirratus* | S America | Neotropical | tropical | sympatric | no |
| *Picus canus* | *Picus viridis* | Asia, Europe | Palaearctic | boreal, temperate | sympatric | no |
| *Picus sharpei* | *Picus viridis* | Asia, Europe | Palaearctic | temperate, subtropical | sympatric | yes |
| *Sphyrapicus nuchalis* | *Sphyrapicus varius* | N America | Nearctic | boreal, temperate | parapatric | no |
| *Sphyrapicus ruber* | *Sphyrapicus nuchalis* | N America | Nearctic | temperate | parapatric | yes |
| *Sphyrapicus ruber* | *Sphyrapicus varius* | N America | Nearctic | boreal, temperate | parapatric | no |
| *Sphyrapicus thyroideus* | *Sphyrapicus nuchalis* | N America | Nearctic | temperate | sympatric | no |
| *Veniliornis frontalis* | *Veniliornis passerinus* | S America | Neotropical | subtropical, tropical | sympatric | no |

**Figure 3.**
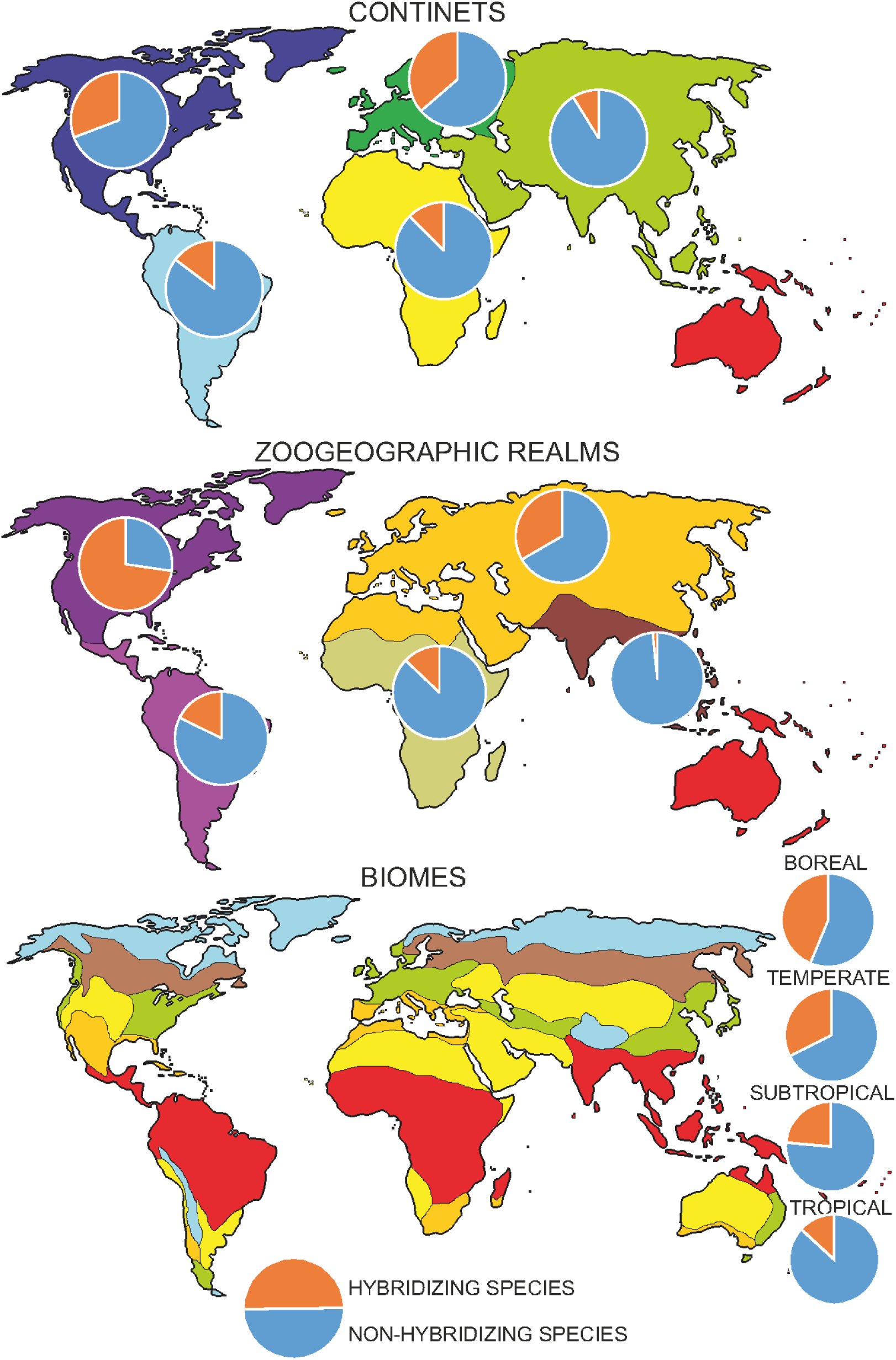
Share of hybridising and non-hybridising woodpecker species in continents, zoogeographic realms and major biomes.

When zoogeographic realms are considered, the majority of hybridising pairs are from the Neotropical (19) and Nearctic regions (16), followed by the Palearctic (6), Afrotropical (4) and Oriental (1) (Table 1). Some hybridising pairs are distributed in Central America, but their ranges span from the Nearctic to the Neotropical realms. Again, the ratio of hybridising species differs significantly when compared to the total number of all woodpecker species in any given region (χ^2^=31.22, p<0.01) (Figure 3).

Considering the major wooded biomes, the largest number of hybridising woodpeckers were found in the tropics (18) and subtropics (17), followed by the temperate (11) and boreal regions (7). As before, the ratio of hybridising species differs significantly when compared to the total number of all woodpecker species in any given region (χ^2^=8.79, p=0.03) (Figure 3).

### Range overlap

Twenty-four woodpecker species pairs are sympatric, at least in part of their ranges, while 13 are parapatric, meeting only in narrow zones (Table 1). Among the remaining pairs of congeneric species (non-hybridising, including these termed uncertain), there were 356 allopatric, 59 parapatric and 75 sympatric cases. A comparison of the frequency of sympatry vs parapatry and that of hybridising and non-hybridising pairs found that they did not differ (χ²=1.555, p=0.213).

In some regions, especially Europe and North America, distribution range shifts occurred quite recently (in the most recent case, just two hundred years ago) owing to changes in land use by humans (mostly forest plantations in the former prairie belt in North America and deforestation in Europe), so some of the currently sympatric/parapatric species may have had allopatric ranges in the recent past. Recent anthropogenic change most likely enabled hybridisation between, for example, *Dendrocopos syriacus* (expanding from Asia Minor into Europe) and *D. major*, and *Melanerpes hoffmannii* and *M. aurifrons* in North America (Short & Morony 1970, Gorman 1999).

### Karyotypes and genomes

Except for some general articles dealing with bird karyotypes that included data on woodpeckers, very few papers described details about chromosomes in woodpeckers that are known to hybridise: *Colaptes auratus* (Shields 1982), *Picus canus* (Xiao-zhuang & Qing-wei 1989), *Picus viridis* (Hammar 1970), *Leuconotopicos villosus* (Shields 1982)*, Dryobates pubescens* (Shields 1982),, *Dendrocopos leucotos* (Lee et al. 1991), *Dendrocopos major* (Shields 1982, Jenni & Müller 1983, Lee et al. 1991) and *Sphyrapicus varius* (Shields 1982). In addition, five studies included information on the general structure of chromosomes or genomes (Shields et al. 1982, de Oliveira et al. 2017, Bertocchi et al. 2018, Romanov 2022, Alves Barcellos et al. 2024). These data showed that most of the Picidae have between 2n=80 and 2n=92 chromosomes, with the noticeable exceptions of *Picumnus cirratus*: 2n=70; *Picumnus nebulosus*, 2n=110; *Melanerpes candidus*, 2n=60 and *Dryobates minor,* 2n=108. An exception may involve the piculets *Picumnus,* but further data are clearly needed as the only two species for which such information is available involve very distantly related species so it unsure whether major karyotypical changes are more common in Picumnus or if the two studies species are exceptions (Shakya et al. 2017).

Genomes are known for nine woodpeckers: *Picus viridis* (Forest et al. 2024; GCA_033816785), *Colaptes auratus* (Hruska & Manthey 2020; GCA_015227895), *Dryobates pubescens* (GCF_014839835), *Melanerpes aurifrons* (Wiley & Miller 2020; GCA_011125475), *Melanerpes formicivorus*; GCF_026170545), *Dendrocopos noguchii* (GCA_004320165), *Sasia africana* (GCA_043658195), *Campethera caroli* (GCA_041296175) and *Pardipicus nivosus* (GCA_041108175) but only the first three species have been assembled at the chromosome level.

### Phylogenetic relatedness

Among hybridising woodpeckers, 15 involved sister species, whereas the remaining 22 were more distantly related phylogenetically. Nearly all cases implied intrageneric hybridisation (Table 1, Figure S1). The only reliable exception involving inter generic hybridisation was for *Leuconotopicus villosus* and *Dryobates pubescens*.

Examination of the genetic distances calculated for three molecular markers, ATP6 (mitochondrial DNA), TGFb2 (autosomal intron) and MUSK (Z-linked intron), revealed that hybridising species (confirmed species pairs) have 2-3-fold lower values of distances than not-hybridising species (considering only congeneric pairs) (Table S2, S3, Figure 4). This resulted in a skewed distribution of pairwise distance histograms of hybridising pairs in contrast to non-hybridising (Figure 4). Differences among three defined groups of woodpeckers were significant for ATP6 (ANOVA = 44.596, p < 0.001), TGFb2 (ANOVA = 42.393, p < 0.001), and MUSK (ANOVA = 47.724, p < 0.001).

**Figure 4.**
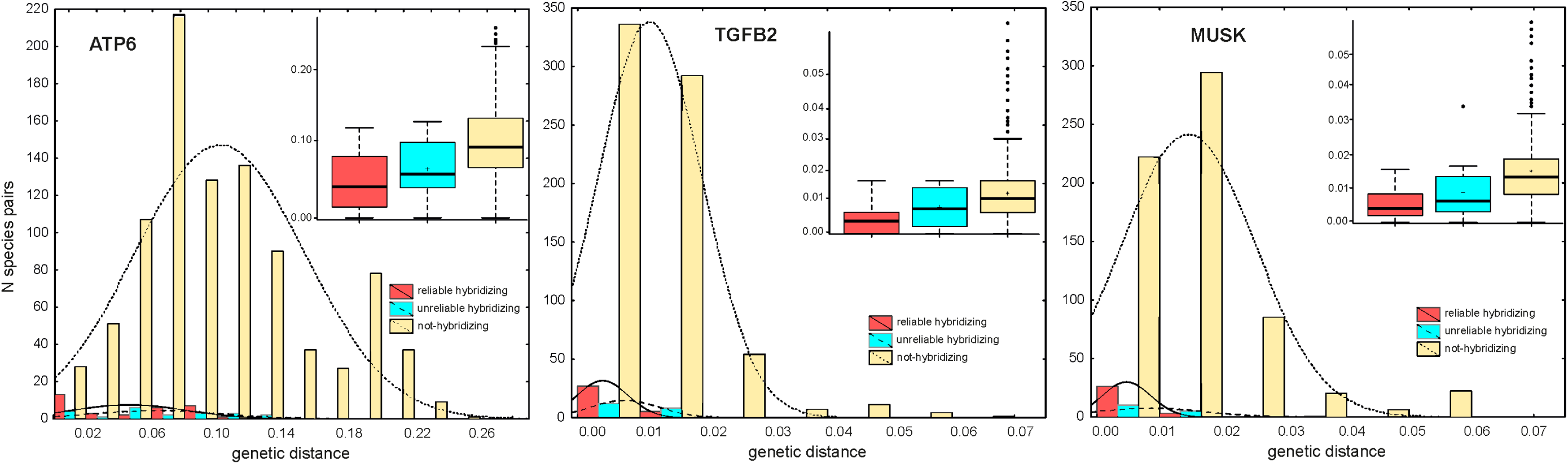
Histograms showing the distribution of pairwise distances among hybridising woodpecker species, pairs of species suspected to hybridise, and all remaining pairs of woodpecker species. Box-plots visualise differences in genetic distances measured among three groups of woodpecker species based on three markers: ATP synthase 6 (hereafter ATP6); autosomal transcription growth factor β 2 intron 5 (hereafter TGFB2) and Z-linked muscle skeletal receptor tyrosine kinase intron 4 (hereafter MUSK) (based on Shakya et al. 2017).

### Characteristics of hybridising woodpeckers

Hybrids are known to have occurred between members of 16 woodpecker genera. According to Ottenburghs & Nicolаї (2024), hybridisation has been reliably and most widely reported in the Neotropical region. For example, involving the piculets (*Picumnus*), Ochre-collared Piculet *P. temminckii,* White-wedged Piculet *P. albosquamatus*; White-barred Piculet *P. cirratus,* Ocellated Piculet *P. dorbignyanus* and Varzea Piculet *P. varzeae*) (Fig. 5), the *Melanerpes* Golden-fronted Woodpecker *M. aurifrons,* Red-bellied Woodpecker *M. carolinus*, Hoffmann’s Woodpecker *M. hoffmannii*, Gila Woodpecker *M. uropygialis*, Velasquez’s Woodpecker *M. santacruzi* and Red-crowned Woodpecker *M. rubricapillus*) and the *Sphyrapicus* Red-naped Sapsucker *S. nuchalis,* Red-breasted Sapsucker *S. ruber*, Williamson’s Sapsucker *S. thyroideus* and Yellow-bellied Sapsucker *S. varius* (Fig. 5, 6). In the Afrotropical region, hybridisation has been reliably documented for the *Campethera* species Green-backed *C. cailliautii* and Little Green Woodpecker *C. maculosa*) (Fig. 5) and the *Dendropicos* Grey Woodpecker *D. goertae,* African Grey-headed Woodpecker *D. spodocephalus* and Olive Woodpecker *D. griseocephalus.* Amongst the Nearctic and Neotropical *Dryobates* and *Leuconotopicus,* Downy Woodpecker *D. pubescens,* Ladder-backed Woodpecker *D. scalaris*, Nuttall’s Woodpecker *D. nuttallii* and Hairy Woodpecker *L. villosus* and the *Veniliornis* Dot-fronted Woodpecker *V. frontalis* and Little Woodpecker *V. passerinus* have been recorded. In the Palearctic and Oriental regions, the *Dendrocopos* Sind Woodpecker *D. assimilis,* Syrian Woodpecker *D. syriacus,* Great Spotted Woodpecker *D. major,* White-winged Woodpecker *D. leucopterus* and White-backed Woodpecker *D. leucotos*) (Fig. 5, 6) have been documented as hybridising. In the Neotropical and Nearctic *Colaptes* genus, Red-shafted Flicker *Colaptes auratus cafer* and Gilded Flicker *C. chrysoides*) often hybridise and in the Neotropics the *Celeus,* Chestnut Woodpecker *C. elegans* and Pale-crested Woodpecker *C. lugubris*, the *Dryocopus* Black-bodied Woodpecker *D. schulzi* and Lineated Woodpecker *D. lineatus* and the *Campephilus* Cream-backed Woodpecker *C. leucopogon* and Crimson-crested Woodpecker *C. melanoleucus* have all be reliably reported. In the Palearctic region, are known to hybridise (the Eurasian Green Woodpecker *P. viridis* is known to hybridise with the Grey-headed Woodpecker *P. canus* in Russia and Poland, and with the Iberian Woodpeckers *P. sharpei* in Southwest France. In the Oriental bioregion, known hybridisation exists for the Sri Lankan Black-rumped Flameback *Dinopium benghalence jaffnense* and Lesser Sri Lankan Flameback *D. psarodes* as well as the Pale-headed Woodpecker *Gecinulus. grantia* and Bamboo Woodpecker *G. viridis* in Thailand, have also been documented.

**Figure 5.**
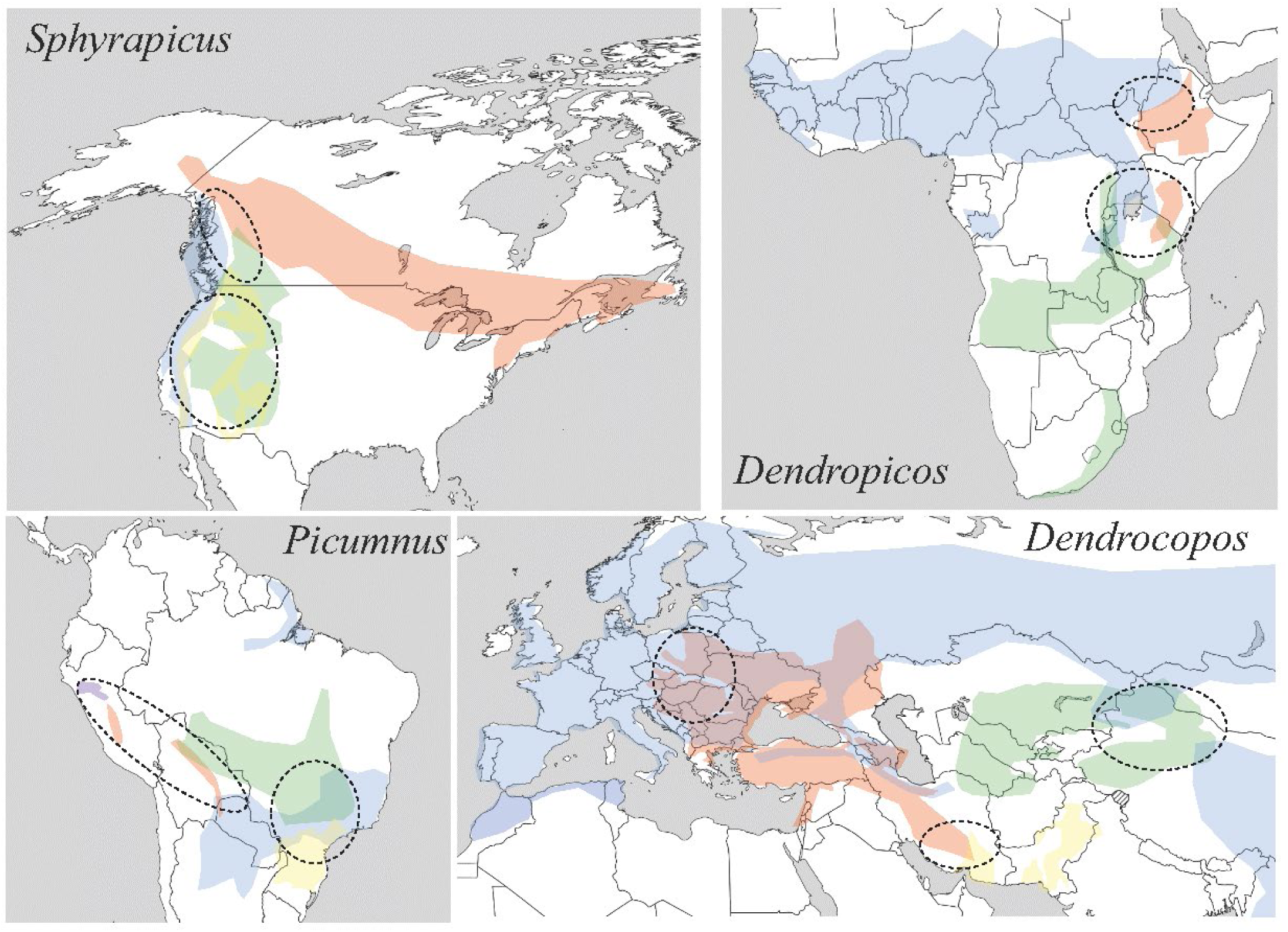
Ranges of some woodpecker species complexes (based on https://birdsoftheworld.org/ and https://datazone.birdlife.org/) that are known to hybridise. North America – genus *Sphyrapicus*: *S. varius* (red), *S. ruber* (blue), *S. nuchalis* (green) and *S. thyroideus* (yellow). South America – genus *Picumnus*: *P. cirratus* (blue), *P. dorbignyanus* (red), *P. albosquamatus* (green), *P. steindachneri* (yellow), and *P. temminckii* (violet). Africa - genus *Dendropicos*: *D. spodocephalus*, (red), *D. goertae* (blue), *D. griseocephalus* (green). Eurasia – genus *Dendrocopos*: *D. syriacus* (red), *D. major* (blue), *D. leucopterus* (green), and *D. assimilis* (yellow). Circles with broken lines correspond to areas of confirmed or presumable hybridisation (including hybrid zones).

**Figure 6.**
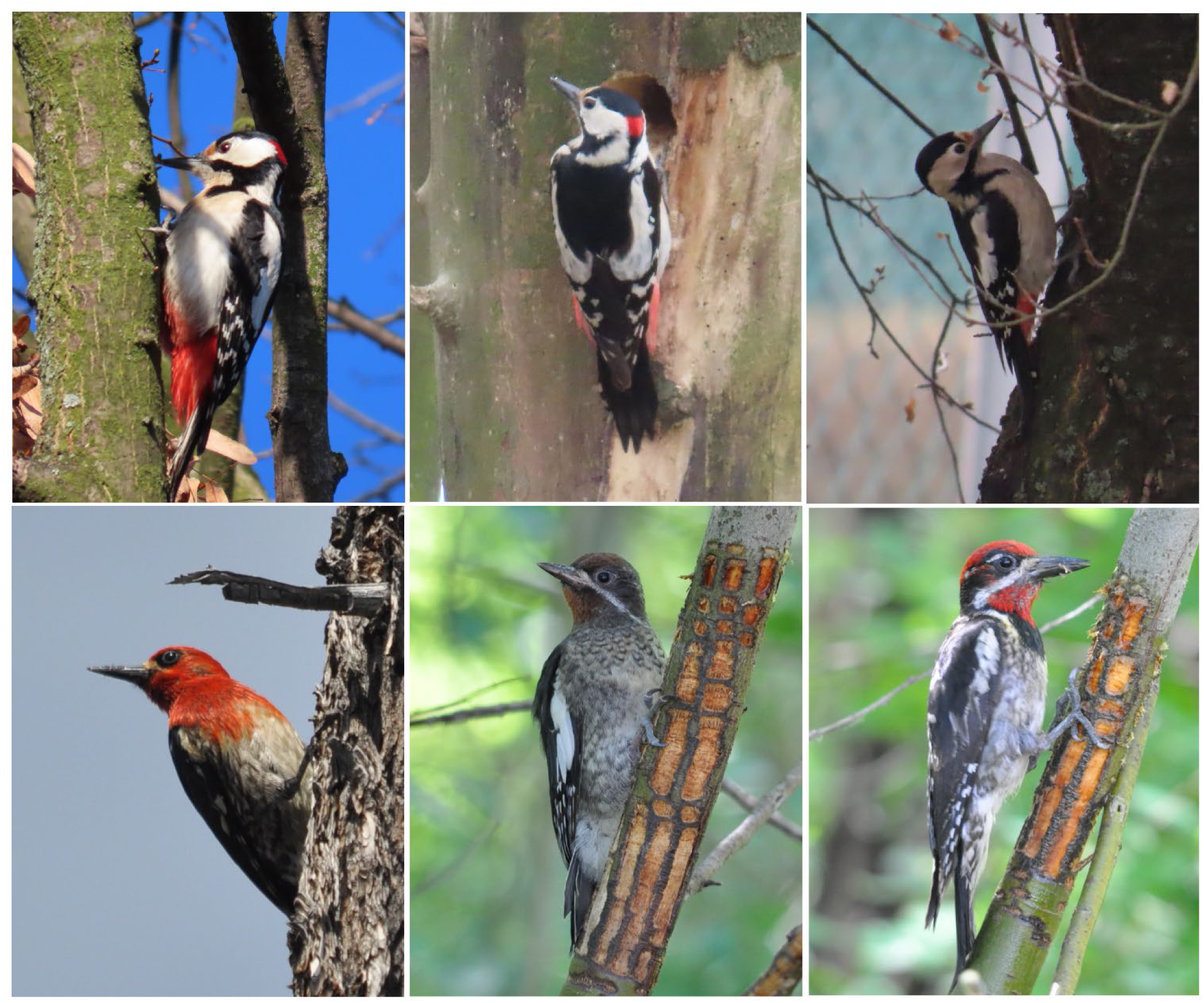
Examples of hybridising pairs of woodpeckers with their hybrids. Upper: *Dendrocopos major*, *D. major* x *syriacus* hybrid, *D. syriacus* (photo Ł. Kajtoch). Lower: *Spyraphicus nuchalis*, *S. ruber* x *nuchalis* hybrid (juvenile), *S. ruber* (photo G. Gorman).

A summary of the species for which, according to Ottenburghs & Nicolaï (2024), firm evidence of hybridisation exists is presented in file S1. Characteristics of pairs of subspecies that were reported as hybridising are presented in supplementary file S2. Uncertain cases are presented in supplementary file S3. Information on collections holding the individuals reported in the literature are summarized in supplementary table S1.

## DISCUSSION

### The evolution of hybridising woodpeckers

Hybridisation can strongly affect the species formation process (Barton 2001), although its effects are often overlooked. At least 46 out of the approximately 240 woodpecker species (∼19%) have been confirmed as hybridising. This value is slightly higher than the recent estimate rate of 16% for all bird species (Ottenburghs et al. 2015), and only lower than the rates for Anseriformes, Charadriformes, Galliformes and Pelecaniformes. On woodpecker evolution, hybridisation has played a continuous role since early times and, in particular, most likely involved the ancestors of the current *Campephilus* and *Melanerpes* genera (Fuchs et al. 2013). Indeed, the extant *Campephilus* species genome possibly still holds a small fraction of the genome left from this ancient hybridisation. If we consider divergence times from Shakya et al. (2017), the mean age of the split between hybridising woodpecker species is about 1.2 Mya (maximum: 6.5 Mya, minimum 200 Kya). This means that currently hybridising woodpeckers mostly (73%) originated during the Pleistocene. During those ages, significant climatic and environmental changes occurred (Whittaker & Horn 2006) due to the increase in frequency and amplitude of the Milankovitch cycles. For example, climate became drier in the Pliocene which caused a contraction and fragmentation of wooded areas, what accelerated during glacial periods in the Pleistocene (Smith et al. 2014, Maffre et al. 2023). Finally, the newly emerging species spread as forest cover increased once again during the Holocene, in the tropical, temperate and boreal zones. Hybridising pairs are more closely related phylogenetically than non-hybridising pairs, which is consistent with the substantial number of sister species found among hybridising woodpeckers. This pattern is consistent across all the examined genetic markers (mitochondrial, autosomal and sex chromosome-linked), although the span of genetic distances is large (mtDNA being more diverse than nuclear introns). Recently split lineages are more similar genetically; thus, their hybrids are less affected by strong incompatibilities and therefore often produce fertile offspring. This process usually does not necessarily ad to the fusion of established lineages or species because hybrids are often counter-selected. Cases of lineages merging have been described (Hogner et al. 2012, Kearns et al. 2018).

Such a process may apply to the Neotropical *Celeus undatus* and *Celeus grammicus* (Sampaio et al. 2018). Among non-hybridising species, pairs with distances close to zero in mtDNA are known. This could be explained by incomplete lineage sorting and low polymorphism of some markers among closely related species (species complexes) (Maddison & Knowles 2006), but undetected hybrids or gene introgression is also likely in some cases (see *Knowledge gaps and perspectives*).

### Biogeography of woodpecker hybridisation

Woodpeckers are widely distributed globally, with the notable exceptions of Madagascar and Australasia (Gorman 1997, Winkler et al. 2014). One of the reasons for the unequal occurrence of hybridising species is that more have been detected in areas where intensive zoological studies and birdwatching (geographic biases in studies, Hughes et al. 2021) have taken place. However, this may be true for the Palearctic and Nearctic America, but is less likely to be so in the Neotropics, where there is a high number of hybridising species, and in sub-Saharan Africa and the Oriental region. Some regions of the world were affected more drastically by climatic and environmental changes in the Quaternary than others, particularly during the Pleistocene glaciations (Whittaker & Horn 2006). Ice sheets were more extensive in Northern Hemisphere due to more emerged lands, and they forced species to retreat to southern refugia (e.g., Mediterranean Basin, Mexican Gulf Coast, etc.). During that period of allopatric isolation, populations diverged and, when the climate changed once more, these populations came into secondary contact during their natural range expansions, putatively resulting in inter-lineage hybridisation. Similar phenomenon also occurred in South America, particularly in mountain regions and in the southernmost part of this continent which was affected by glaciation. The impact of the glacial cycles on population differentiation was possibly heterogeneous across the globe. Examination of the time of origin of species across the four primary bioregions indicate that species are younger in the Neotropics (average: 1.8 Mya) and older in the Oriental region (average 3.4 Mya), with intermediate values for the Nearctic (2.8 Mya; 2.3 Mya excluding two Caribbean outliers with no close relatives *Melanerpes striatus* and *Nesoctites micromegas*), sub-Saharan Africa (2.7 Mya) and the Palearctic (2.1 Mya). Hence, it also appears that the number of hybridising species within a bioregion is also linked to the mean age of species in that realm. The majority of known hybridising woodpeckers are from the Neotropics. This region was once considered to be the original geographical source of woodpeckers (Short 1982), but this view was later rejected on the basis of molecular data with woodpeckers now thought to have colonised South America multiple times at the end of the Miocene (Fuchs et al. 2007). The formation of the Panama Isthmus, around 3 Mya (O’Dea et al. 2016), enabled the spread of some woodpecker lineages from North to South America (*Colaptes, Dryobates, Leuconotopicus, Melanerpes*). The mean divergence time between Nearctic hybridising species was around 1.8 Mya (Shakya et al. 2017), where it is about 1 Mya for Neotropical species (Table S5). Thus, the formation of a land bridge between the Americas probably did not provide more opportunities for hybridisation. Far fewer hybrid examples are known from the Afrotropical and Oriental regions. These two regions were not significantly affected by Pleistocene climatic fluctuations, although there was a retreat of forest cover owing to the drying out of land (Rosqvist 1990), that could have affected speciation and the formation of a secondary contact zone. The mean divergence time of hybridising woodpeckers in the Oriental region is 0.6 Mya, which is much more recent than the average divergence of woodpeckers in this region (3.4 Mya). In the Afrotropical region, hybridising species diverged around 2.0 Mya, and this estimate is close to the average divergence time of all African woodpeckers.

The highly unequal occurrence of hybridising species in the tropics (which is more common in the Neotropics than in the Afrotropic or Oriental regions) is puzzling. Most likely this situation developed in the time when speciation occurred and when opportunities for further divergence occurred. Species are older r in the Afrotropic and Oriental realms (end of the Pliocene) when compared with the Neotropics (mid Pleistocene) (Shakya et al. 2017). The presence of many “young” species in South and Central America more readily enables the formation of hybrids, in contrast to “older” species in Africa and SE Asia, but this is not a “rule”. There are known examples of woodpecker species diverging relatively recently and hybridising, such as *Dinopium* flamebacks on a tropical island in SE Asia (Sri Lanka) (Ranasinghe et al. 2024). This example provides evidence of cryptic diversification and within-island population divergence, highlighting the complexity of hybridisation and speciation processes. Moreover, there are more para- or sympatric species in the Neotropics than in the Old World tropics (Ottenburghs & Nicolaï 2024) and this may have facilitated interspecific breeding in the former region.

Differences between hybridising species in temperate and boreal regions and Nearctic and Palearctic regions, are also visible in North America than in Eurasia. Explaining this difference may involve the fact that there is an overlap in Neotropical and Nearctic woodpeckers at the borders of these areas; some Neotropical genera are also found in Mexico and the southern USA (Gorman 2014), whereas in Eurasia there are geographic barriers between Palearctic and Afrotropic species (the Mediterranean Sea and deserts) and Oriental species (mountain ranges and deserts).

Many hybridising woodpeckers are known from mountainous areas, a fact that may have played a role in altitudinal isolation of populations and the further connection of newly emerged woodpecker species, for example, in the Andes in South America (*Campephilus, Celeus, Picumnus, Veniliornis*), the Rocky Mountains in North America (*Sphyrapicus, Melanerpes*), mountains in east Africa (*Campethera*) and mountains in central Asia (*Dendrocopos*) (Fig. 5).

Yet, there are known stable hybridisation zones, including particularly well-documented ones in North America involving *Sphyrapicus* and *Melanerpes* (Seneviratne et al. 2012, 2016, Barrowclough et al. 2018, Billerman et al., 2019, Natola et al. 2021, 2022, Lanes-Quevedo et al. 2022). Likely hybrid zones also exist where parapatric ranges of other woodpeckers meet, such as is the case with *Dendrocopos* species in Central Asia (https://birdsoftheworld.org/) or *Picus* in the Western Palearctic. In *Dendrocopos*, there is an example of a species pair that started to hybridise in very recent times due to environmental changes caused by human activity - deforestation of SE Europe and the expansion of rural and urban green areas (Michalczuk 2014). This most certainly enabled the expansion of Syrian Woodpecker from its native range in Middle East into Europe, where it meets and hybridises with Great Spotted Woodpecker which is mainly a forest-dwelling species (Figarski & Kajtoch 2018).

### Genomic background of woodpecker hybridisation

Hybridisation of woodpeckers is dependent not only on their phenotypic properties (similar plumage, voices, habitat requirements and breeding biology) but also on their genomic conditions. Karyotypes of woodpeckers are known to be constituted by substantial number of micro chromosomes (Romanov 2022). The majority of woodpecker species have a typical diploid number of chromosomes for birds, with most woodpeckers being around 80-92 (Barcellos et al. 2024). Only a few woodpeckers have a lower figure (2N=64 in *Melanerpes candidus*; de Oliveira et al. 2017 and 2N=70 in *Picumnus cirratus*, Gunski et al. 2004c) or higher (2N=108 *Dryobates minor*, 2N=110 in *Picumnus nebulosus*, Barcellos et al. 2024). Thus, some woodpecker lineages show significantly rearranged karyotypes compared to the putative ancestral karyotype (Barcellos et al. 2024). At the same time, what is characteristic for woodpeckers is enlarged sex chromosomes (Z) (Shields et al. 1982), that have repetitive DNA like microsatellites and transposable elements (de Oliveira et al. 2017). It is likely that sex chromosomes were crucial in speciation and play a vital role in hybridisation (Shields 1982). Noticeable hybridising species with known karyotypes have the same number of chromosomes, as in the case of *Leuconotopicos villosus* and *Dryobates pubescens* (2N=92), or very similar *Dendrocopos major* and *D. leucotos* (2N=90 or 92 depending on study), *Picus canus* and *Picus viridis* 2N=92 and 2N=94, respectively). Likely hybridising species presumably have the same or very similar numbers and structure of karyotypes otherwise prezygotic isolation would occur.

Genomes are known for several woodpecker species but studies comparing genomes of hybridising species are nearly absent, therefore determination of (dis)similarities in genome structure between woodpecker species that hybridise and those which do not interspecifically breed are not known. The same goes for hybrids, with the exception of sapsuckers *Sphyrapicus*, which are well-studied (Seneviratne et al. 2016, Grossen et al. 2016, Natola & Burg 2018, Natola et al. 2021, Billerman et al. 2019), as well as the Northern Flicker *Colaptes auratus* (Aguillon et al. 2021) and *Dryobates* (Manthey et al. 2019).

Several studies focused on the sapsuckers hybrid zones throughout the western nearctic (British Columbia, Canada: Seneviratne et al. 2016, Grossen et al. 2016, Natola et al. 2022; Alberta, Canada: Natola and Burg 2018; California and Oregon: Billerman et al. (2019). These studies concluded for more hybridisation and lower genome wide differentiation between *S. ruber* and *S. nuchalis*, that splitted about 0.3 mya, than between *S. ruber* and *S. nuchalis*, that splitted between 0.8 mya (Shakya et al. 2017). The frequency of *S. ruber*/*S. nuchalis* hybrids in the hybrid zone was highly variable (77% in California and Oregon versus 15 % in British Columbia (Seneviratne et al. 2016, Billerman et al 2019). Such heterogeneity was attributed by Billermena et al. (2019) to the difference in habitat transition between the two areas (transition being clinal in British Columbia versus patchy in California/Oregon). In both cases, available data suggested some form of selection against hybrids, with F1 and F2 hybrids being rare (Seneviratne et al. 2016, Billerman et al. 2019) suggesting pre-zygotic isolation via possibly assortative mating. Genome scan possibly identified one locus on Chromosome 11 that was associated to plumage characters (Grossen et al. 2016). A study of the tri-species hybrid zone in British Columbia, where each species density is variable and each combination of crossing is seen, found strong evidence for assortative mating and that hybrids do not suffer from lower survival (Natola et al. 2022). The authors proposed that this hybrid region is three-way tension zone maintained by reduced migration in and out of the hybrid zone and selection against hybrid (Natola et al. 2022). Natola & Irwin (2023) supported a important role of Z chromosomes in the speciation of *Spyraphicus* woodpeckers as sex chromosomes show a 2X-3X higher relative differentiation (Fst) between species than autosomes and harbour signatures of recurrent divergent selection, high linkage disequilibrium, and likely results in epistasis among Z haplotypes. Even greater genomic differences between species pairs which separated long ago not necessarily imply reproductive isolation, so there are lengthy periods of time when hybridisation is possible if diverging populations are in geographic contact.

The situation in the Northern Flicker (Colaptes auratus ssp) is somewhat similar, with very little genome wide differentiation (Fst=0.08) and approximately six times differentiation of the Z-chromosomes than autosomes (Aguillon et al. 2021). Both forms, referreed as red- and yellow-shafted flicker could be easily distinguished on the PCA, despite the absence of fixed single-nucleotide polymoprhism with hybrids occupying intermediate positions. Genome wide associations allowed the identification of putative twelve genetic regions associated to differences in colors between the two forms (three of them being on the Z chromosome) (Aguillon et al. 2021).

Finally, Single nucleotide polymorphism panels and whole-genomic data revealed that three hybridizing *Dryobates* woodpecker species are genomically distinct. Apart from hybridization between *D. pubescens* and the remaining species, the low levels of gene flow occur between *D. nuttallii* and D*. scalaris* across the contact zone with no evidence for widespread genomic introgression between these species (Manthey et al. 2019).

### Hybridisation as an undervalued factor in woodpecker studies and conservation

Apart from ancient introgression events, interspecific breeding has had a notable influence on woodpeckers if hybrids are fertile and became common in emerging populations. Hybrids harbour alleles from two genetic pools; therefore, they are highly heterozygous (Fitzpatrick 2012). This could enhance their fitness owing to a greater genetic polymorphism and possession of alleles that may allow for better adaptations to environmental conditions and/or changing behaviour and/or physiology. On the other hand, if alleles that converge in a hybrid individual are not compatible, it could decrease hybrid vitality and/or fertility (Moehring 2011). Unfortunately, studies that deal with the consequences of hybridisation on woodpecker fitness are lacking due to their low densities, certainly when compared to passerines. On the one hand, hybrids and backcrosses, for example within the *Dendrocopos*, *Colaptes*, *Picus* and *Sphyrapicus,* are known to be fertile (Aguillon et al. 2018; Billerman et al., 2019; Pons et al. 2019; Kajtoch & Kusal 2022). In the case of some genera, well-known populations consist of ancestral “pure species” birds, hybrids and backcrosses, which suggests that an existence as “hybrid swarms.” (Manthey et al. 2016). This phenomenon might be stable in the well-established contact zones, which are maintained according to tension zone (*Sphyrapicus, Picus*) or hybrid superiority (*Colaptes*) models, which prevent the spread of hybrids across both ancestral species ranges and contain clinal barriers preventing genome introgression into allopatric populations (Manthey et al. 2016; Billerman et al., 2019). Nevertheless, molecular studies on hybridising woodpeckers are still often rather superficial and introgression is poorly known. Hybridisation between recent overlapping species might appear to be different but, for example, introgression from Syrian into Great Spotted Woodpecker likely facilitated the recent expansion of the later into urban habitats which could pose a threat for Syrian Woodpecker (Kajtoch & Kusal 2022). Such genomic exchange could be an opportunity or threat for the conservation of woodpeckers (see reasons listed above). Unfortunately, hybrids are neglected in nature protection laws in most countries. Yet, hybrids have started to be appreciated also in conservation policies in the USA and Canada (Piett et al. 2015).

### Knowledge gaps and perspectives

Most of the documented cases of woodpecker hybridisation have been occasional. This is probably because hybrids rarely occur, or perhaps that finding and identifying interspecific individuals is difficult in the field, particularly in remote areas such as in the tropics. Most information on hybridising woodpeckers is anecdotal, often published in local journals or, in recent times, in databases, discussion groups and social media. Despite a few actively studied species, such as *Spyraphicus* (Johnson & Johnson 1985, Trombino, 1998, Seneviratne et al. 2012, 2016, Billerman et al., 2019, Natola et al. 2021, 2022), *Picus* (Pons et al. 2011, 2013, 2019, Fuchs et al. in prep), and *Dendrocopos* (Gorman 1997, Michalczuk 2014, Michalczuk et al. 2014, Figarski & Kajtoch 2018, Gurgul at al. 2019, Kajtoch & Kusal 2022), research on the other hybridising woodpeckers is scarce. In the best cases, publications have basic information on the plumage of hybrids, but the information on biology, ecology and behaviour is lacking. The other problem is that authors are not always given information about the environment and habitat where hybrids or mixed broods were seen. Missing or deficient information is mostly a result of difficulties in studies on hybrids, which are often hard to detect in field and classical methods of bird inventories and research usually does not allow for proper identification and collection of reliable data (Bakai et al. unpublished). Some pairs of woodpecker species are suspected to hybridise but are not confirmed as doing so (Ottenburghs & Nicolaï 2024). Interestingly, when looking at the unconfirmed pairs of hybridising woodpecker species, the phylogenetic distances between them are on average intermediate between hybridising and not-hybridising species (Table S1). It should be verified whether or not some of these “unreliable” species pairs indeed are capable of mating, but the lack of observations or difficulties in determining hybrids are the reasons that some potentially hybridising pairs are still unconfirmed. Among 27 such pairs of woodpecker species, 12 have sympatric and 4 parapatric ranges, so geographically they are able to meet. Among these pairs of sister species, hybridisation is most likely in the following: *Picumnus minutissimus* x *Picumnus spilogaster*, *Sasia abnormis* x *Sasia ochracea*, *Campethera abingoni* x *Campethera mombassica*, *Dendropicos gabonensis* x *Dendropicos lugubris*, *Veniliornis lignarius* x *Veniliornis mixtus*, *Dendrocopos atratus* x *Dendrocopos macei*, *Celeus grammicus* x *Celeus undatus*, *Picus viridanus* x *Picus vittatus*, *Dinopium javanese* x *Dinopium shorii*. These pairs should be particularly examined to verify if hybrids between then really occur and for that museum collections should be used. Sampaio et al. (2018) revealed a lack of genetic distinctiveness between *Celeus grammicus* and *Celeus undatus*, which means their statis as species should perhaps be reconsidered. This may also be a case of despeciation similar to that seen in ravens (Kearns et al. 2018). Phylogeographic studies have been conducted for all biogeographic regions where the Picidae are found, hence information and further hypotheses on where putative contact zones among avian lineages should exist. Studies aiming to evaluate the possibility of hybridisation among less studied woodpecker species, or bioregions, can take advantage of this information to screen for the presence of specimens in natural history collections from these areas. Online databases (Vertnet, Gbif) would allow the identification of specimens prone to further morphological or genetic examination. It should be noted, however, that the identification of species that are known to be able to interbreed, is uncertain when it is based only on their plumage characteristics as these were probably similar amongst ancestral species. To improve identification and limit errors, the incorporation of modern techniques in research on hybridising woodpeckers is needed. The use of photography (including remote sensing) will also help find and define the plumage features unique for parental species and their hybrids. Artificial intelligence and advanced statistical methods (e.g., machine learning) (Petso et al. 2022) could be used, too, although records should only be accepted after verification by specialists. The following method should be combined with the use of citizen science - the collection of data (records with photographs) from various localities and times (Perry et al. 2021). Better understanding of morphological characters of hybrid woodpeckers should also be applicable for museum specimens among which still could be uncovered hybrid individuals of those species which are currently uncertain. Nevertheless, morphology is not always a reliable way of proving hybridisation, as variances in the phenotype and haplotype of hybridising birds are known (Natola et al. 2022) and atypical plumage can be caused by diverse kinds of aberration but reports on presumable hybrids is still valuable information. This problem could be overcome with the use of rapidly developing molecular techniques (Cottenet et al. 2020). Genetic data open much better ways of studies on hybridisation starting from certain determination of hybrids and backcrosses, though understanding of mechanisms of hybrid development referring to woodpecker biology, ecology and ethology, to understanding genomic basis of hybrid anatomy, physiology (Payseur & Rieseberg 2016). Unfortunately, current knowledge and data is insufficient to tell how differences in ecology influence hybridisation in woodpeckers. Among the documented species pairs (both dependable and unreliable), all birds are similar in general plumage patterns and morphology, although some morphometric features (bill shape, body size, tarsus length, etc.) do differ between hybridising species, as well as distinct plumage features exist. Those studies that compare hybridising species and the behaviour of hybrid young birds, and how that influences mixed pair formation, are also limited to the most commonly studied species. As far as we know, *Dendrocopos syriacus x D. major* hybrids behave is the same manner as their pure parents, with individuals actively responding to the territorial calls of both species and defending their territories (Figarski 2017). The same has been seen for *Sphyrapicus ruber x S. nuchalis* hybrids living in sympatry with “pure” birds (Billerman & Carling 2017), although some differences in drumming signals might indicate a form of interbreeding barrier (Trombino 1998). On the other hand, significant differences between *Picus canus* and *P. viridis* courtship behaviour present a major obstacle to mixed pair formation (Fridemann 2011). Despite these examples, the behaviour component role in mixed pair formation is still unclear and ethological studies on the other hybridising woodpecker species are needed to clarify this question. Modern molecular techniques could also be important in studies of museum samples, especially of rare species or woodpeckers from regions that are difficult to access (Burrell et al. 2015). Finally, a better understanding of woodpecker hybrids could be helpful in the planning of proper conservation actions for other rare and vulnerable species, which would be especially important in a world affected by environmental and climatic changes that are causing a decline in biodiversity (Walters et al. 2020). Overall, hybrids are natural elements of biodiversity that play a significant role in evolution and ecology (Saetre 2013).

## Supplementary materials to the article

File S1. Characteristics of woodpecker species with known records of hybridisation.

### Genus Picumnus

This genus currently includes 19 species, most found in South America and, one in Central America. Single species in Southeast Asia (*Picumnus innominatus* is considered as *Vivia innominate*).. All woodpeckers are small (up to 10 cm in length) with short bills and tails. The plumage colouration and patterns differs among the species, but all birds have rather blank colours, distinct colours of back and abdomen and most have the black crown with tiny white spots. The males have red, orange or yellow forehead. The birds in this genus exploit the various types of habitats of diverse humidity and elevation, even within a single species. Some species easily occupy human-transformed habitats such as arable land and pastures. According to Ottenburghs & Nicolаї (2024), there are seven species pairs that have hybridised, two of which are “doubtful”.

Ochre-collared Piculet *P. temminckii*, White-wedged Piculet *P. albosquamatus*, White-barred Piculet *P. cirratus* and Ocellated Piculet *P. dorbignyanus* are all known to hybridise with each other (Neto 1995; Short 1982; Winkler et al. 1995; Billerman et al. 2022), though no Ocellated x Ochre-collared hybrid has been documented. Although White-wedged and White-barred seem to prefer different habitats (open dry savannas and forest galleries) to Ocellated (humid tropical montane forests) and Ochre-collared (humid forests), all of them show preferences for shrubs in open areas and thus could meet at forest edges, and able to occupy light lowland moist forests. White-winged x Ochre-collared hybrids from Santa Mariana, Brazil, have been described as mainly resembling White-barred Piculet, but with their undertail more like that of Ochre-collared, with the throat and chest having an intermediate pattern.

White-barred Piculet is also mentioned by Short (1982) as hybridising with Varzea Piculet (*P. varzeae*) along the Amazon River in the East-North of Brazil, as both species occur in lowland scrub and forest ecotones with dense undergrowth.

### Genus Melanerpes

These small to medium New World woodpeckers occupy a variety of habitats. Most occur in Central America and the Caribbean, with a few in South and North America. Sexual dimorphism is expressed in various distinctive head plumage features, such as nape patches, forehead, caps and crown colour.

According to Ottenburghs & Nicolаї (2024), four species have been confirmed as hybridising, and one subspecies pair said to be doubtful, although it is supported by Lanes-Quevedo et al. (2022).

Golden-fronted Woodpecker *M. aurifrons* inhabits Central and partly North America and is known to hybridise with three species in the same genus: Red-bellied Woodpecker *M. carolinus*, Hoffmann’s Woodpecker *M. hoffmannii* and Gila Woodpecker *M. uropygialis* in the southern USA, southern Honduras and Central Mexico, respectively. The Golden-fronted Woodpecker is adaptable, able to occupy a wide variety of habitats, from sea level to 2500 m, from mesic to xeric environments and swamps, open areas, dense forests and also urban and suburban habitats.

Golden-fronted and Red-bellied Woodpeckers come into contact n S Oklahoma and E Texas, where both species share their preferred habitats of mosaic riparian, oak woodlands and savannah. Both species are omnivorous, consuming a significant amount of plant food. Smith (1987) provides morphological and genetic (protein electrophoresis) evidence for the hybridisation and introgression of the two species. According to this author, the proportion of hybrid individuals can be 15.8% and it is also stated that “pure” females and hybrids of the two species are more easily distinguished, whereas male hybrids are harder to identify as they have a higher variation in plumage patterns and colour. Female hybrids were characterized by large white patches on the tail, an absence of tail barring, red napes and orange bellies. Male hybrids were described as having the same features as female, but with additional ones such as orange breasts and no extra lateral colour on the nape. The author also pointed out that both species had undergone recent range expansions, which led to an increase in their overlap zone. It is, however, unclear whether environmental factors block their further expansion or competition between these two species. Smith (1987) also suggested that females of both species may prefer Red-bellied males owing to their more extensive red plumage.

Short (1982) describes Hoffmann’s Woodpecker as parapatric with Golden-fronted, and that hybridisation between them occurs in a narrow zone along the Pespire River in Honduras. Hybridisation between these species is also mentioned in Winkler et al. (1995). Both species occupy open and semi-open xeric environments but also plantations and urban habitats. The Hoffmann’s Woodpecker is smaller than the Golden-fronted Woodpecker.

Selander & Giller (1963) writes about contact zone between Golden-fronted Woodpecker and Gila Woodpecker in northern Mexico. The authors documented few mixed pairs and even described a several hybrid specimens in the areas where both species occurred, but one of them was less frequent. They describe the hybrid individuals to be rare and having many intermediate plumage features, visibly different in pattern from aberrant individuals. According to Short (1982), the range of overlap between these species is limited to some parts of Aguascalientes and Jalisco, Mexico, where hybridisation involves about 5% of the population. This hybridisation is also mentioned by Winkler et al. (1995). Although the Golden-fronted Woodpecker is less adapted for arid habitats, the two species met in the highland (5000-6000 feet) river banks at northern and central Mexico. Selander & Giller (1963) pints that after extensive human alternation of land in the Mexico, some natural suitable habitats for these woodpeckers were lost at the local scale, but at the same time many suitable habitats for them appeared in the large areas (mainly due wood planting in the agricultural areas) which resulted in range expansion of both species at the central and northern Mexico.

Velasquez’s Woodpecker *M. santacruzi* was previously considered a subspecies of Golden-fronted Woodpecker. Lanes-Quevedo, et al. (2022) provide evidence a significant genetic admixture between these two species in E Mexico where their ranges overlap. Winkler (1995) wrote that Velasquez’s and Hoffmann’s Woodpeckers interbreed in a small area of overlap in NE Honduras but treated Velasquez’s Woodpecker as the subspecies of the Golden-fronted Woodpecker.

Hoffmann’s Woodpeckers avoids dense, humid woodlands, and lives in xeric and mesic areas in open and semi-open country, as well as in urban and suburban habitats. Cases of Hoffmann’s Woodpecker hybridisation with the smaller, Red-crowned Woodpecker *M. rubricapillus* in Costa Rica are also known (Stiles & Skutch 1989). Red-crowned Woodpecker inhabits open and semi-open areas (forest edges, clearings, secondary growth, coastal shrubland), plantations and gardens, so co-occurrence in the same habitat in this pair of species is possible.

### Genus Sphyrapicus

*Sphyrapicus* genus of four medium-sized woodpeckers is endemic to North and Central America. All four species are similar in colour, mostly consisting of a black-and-white body with a yellowish breast and red on the head. Sexual dimorphism is minimal in three species but extreme in Williamson’s Sapsucker. These closely related species co-occur in many areas and hybridisation is common (Grossen et al. 2016; Natola 2017).

Ottenburghs & Nicolаї (2024), state that there are four species pairs that can hybridise. Red-naped *S. nuchalis* and Red-breasted *S. ruber* Sapsuckers are sister species that hybridise in several areas from S British Columbia to California. Red-naped is the smaller species and prefers the interior of dry coniferous forests, while Red-bellied inhabits more humid and closed forests dominated by hemlock (*Tsuga*). The first species is characterized by the higher dichromatic sexual dimorphism (male have a completely red throat, while females have a white chin) than the second (almost monomorphic). Their hybrids are reported to have variable intermediate phenotypes with features from both parental species, and in some cases almost entirely resembling one of them (Johnson & Johnson 1985; Natola et al. 2022). Johnson & Johnson (1985) describe 13 hybrid types mainly based on their head plumage patterns. The drumming of the two species is very similar where they are sympatric but different where they are allopatric (Trombino, 1998). However, there are no scientific works that show differences in vocalisations. The hybrid zone between these two species is a narrow area (65 km wide) with a steep environmental gradient from warm and wet to cool and dry in S British Columbia. No gene flow has been observed outside the hybridisation zone (Natola et al., 2023). The backcross individuals arereported to be rare in the local populations, comparable to birds with of parental phenotypes or F1 hybrids. Their share in the local populations is lower than the predicted one under conditions that two species interbreed freely, for reasons that are unclear (Johnson & Johnson, 1985). Some authors report that hybrids have reduced return rates (Trombino, 1998), but others did not find evidence for this (Natola et al. 2022). Hatching success also proved to be similar to that of “pure” pairs (Trombino, 1998). Experiments with genetic screening of individuals show that 77% of birds had a genetic mixture, but the majority were backcrosses (Billerman et al., 2019). The proportion of males among hybrids seemed to be bigger than females. Population genetical analysis and the rarity of F1 hybrids pointed to assortative mating in the hybrid zone (Johnson & Johnson, 1985; Natola, 2017). Analysis of nesting pairs also showed that birds tended to pair with mates of the same phenotype, and that hybrids tended to backcross with one of their parental species, while hybrid x hybrid pairs were extremely rare (Johnson & Johnson 1985). Most mixed pairs had males with redder heads. Experiments with playback as a stimulus for aggression in sapsuckers showed that Red-naped are more aggressive than Red-breasted (Billerman & Carling, 2017). Genetic clines, minimal hybrid phenotypes and a narrow hybridisation zone, are consistent with the ‘tension zone’ model, which means the zone persists because selection against hybrids is balanced by dispersal of the parental species (Seneviratne et al. 2016). The hybrid zone has shifted recently due to climate change, Red-breasted expanding eastwards into the range of Red-naped, and replacing it in some locations (Billerman et al., 2016; Billerman et al., 2019).

Hybridisation between Red-naped and Williamson’s S. thyroideus sapsuckers was recorded for the first time by Oberholser (1930), while Short & Morony (1970) later described a hybrid between these two species in more detail. According to his description the hybrids resembled Williamson’s more than Red-naped. Both specimens were found in wintering areas in Mexico. In the breeding area, these species usually overlap in montane forests in California, but they differ significantly in their habitat preferences and behaviour. Red-naped prefers deciduous forest stands with aspen (*Populus tremuloides*), while Williamson’s is more attached to open coniferous forests with Ponderosa pine (*Pinus ponderosa*).

The Yellow-bellied Sapsucker *S. varius* inhabits boreal forests east of the Rocky Mountains. The hybridisation zone is in NW Alberta, Canada, and around 275 km wide. These two species are similar in plumage, differing in red-naped sapsucker having a red nape, and more yellow on the belly and white in the wings than Yellow-bellied Sapsucker. Also, Yellow-bellied Sapsucker is more sexually dimorphic, females having a distinct white throat and chin. Parental and hybrid phenotypes can be distinguished by a few plumage features: extent of the red malar stripe and cheek patch in males, amount of red on the throat of females and amount and extent of white markings on the upperparts. Researchers have defined 8 hybrid indexes, but the frequent mismatch between phenotype and genotype classes can be seen (Natola et al. 2021). Genetic studies, however, show a clear pattern of introgression in both species. These two sapsuckers are more alike in their ecologies than the other two species in this genus. Both prefer secondary growth deciduous and mixed forests, particularly with aspen and birch. *S. nuchalis* breeds at higher elevations and winters in S California and N Mexico, while Yellow-bellied Sapsucker migrates to Central America and the Caribbean. In Natola et al. 2021, hybrids were mostly found in riparian corridors with large aspens in the ecotone between montane and boreal forests, where suitable habitats are scarce which results in low densities. There is no proof of assortative mating.

Red-breasted and Yellow-bellied Sapsuckers also hybridise in an 84 km wide zone in northern British Columbia. Yellow-bellied Sapsucker has prominent white and black patches, while Red-breasted Sapsucker has mostly red head and breast. The Red-breasted Sapsucker is slightly bigger than Yellow-bellied Sapsucker. Yellow-bellied Sapsucker males showed a higher level of heterospecific aggression towards Red-breasted Sapsucker decoys, than vice-versa in the hybrid zone (Seneviratne et al. 2012). Hybrids show various intermediate plumage features, for example, *ruber*-type has black-and-white spotting on the face and *varius*-type has red patches on the face and breast. There are speculations about females of both species to show preference for reddish plumage in males, causing assortative mating shift towards more reddish Red-breasted males, resulting in asymmetric hybridisation pattern (Seneviratne et al. 2012). The hybridisation zone, in the Rocky Mountains in NW British Columbia, is 122 km wide. The furthest away records of genetic introgression from the centre of hybridisation are 100 km to the west (Seneviratne et al. 2012). About 32% of birds in the Rocky Mountains contact zone were hybrids, 83% of them backcrosses (Natola et al. 2023). The parental species have different migration routes and wintering areas, thus F1 hybrids were thought to have lower survival rates due to a maladaptive migration strategy. However, analysis of sapsucker age classes did not show hybrid disadvantages in survival (Natola et al. 2022).

There is genetic evidence for hybrids involving three sapsucker species, *ruber × nuchalis × varius,* in a zone in the Prince George area, Canada, with the highest proportion of genomes relating to Red-breasted Sapsucker (Natola et al. 2022). Most of the birds in this zone were backcrosses in further generations (second or more) (73.3%), with F1 hybrids constituting 3.3%. Phenotypically, most were Red-breasted Sapsucker (70.4%), with others having hybrid (20.2%), Red-naped Sapsucker (4%) or Yellow-bellied Sapsucker (5.4%) phenotypes. The hybrid phenotypes present in this zone were either Red-breasted Sapsucker × Red-naped Sapsucker or Red-breasted Sapsucker × Yellow-bellied Sapsucker, but the traces of genetic introgression of each sapsucker species could be found in all three “pure” phenotypes (e.g. from Red-breasted Sapsucker to Red-naped Sapsucker and vice versa). There were also a few birds for which phenotype did not match their genomic clusters, that is, “pure” looking birds with a hybrid genotype or hybrid-looking birds with a distinct genotype (Seneviratne et al. 2016; Natola et al. 2022).

### Genus Campethera

The genus Campethera includes ten African species which generally have white underparts, green wings, back and tail and red napes. Sexual dimorphism is expressed in males having a red crown and malar stripe. According to Ottenburghs & Nicolаї (2024), only one species pair is known to hybridise.

Prigogine (1987) mention a hybrid between Green-backed *C. cailliautii* and Little Green Woodpecker *C. maculosa* in Ghana. Both species are adaptable in habitat preferences and are very closely related (divergence time about 0.5 Mya, Fuchs et al. 2015).

### Genus Dendropicos

This genus currently includes 12 small to medium woodpecker species distributed across sub-Saharan Africa, but occupying different habitat types. Their plumages vary from yellow to grey, often with green wings, and males possess a red crown. Ottenburghs & Nicolаї (2024) listed three reliable and one doubtful case of hybridisation.

Grey Woodpecker D. goertae, African Grey-headed Woodpecker D. spodocephalus and Olive Woodpecker D. griseocephalus are considered to form a superspecies and confirmed to interbreed. Short (1982) writes that Grey and Olive may overlap in Kilimanjaro, Tanzania, and N Ituri in DR Congo.. M. Loutte found that these species hybridise in Rwanda (Prigogine & Louette (1983), although in the Birds of Rwanda (Gaël 2018) they are described as having different habitats. In the previous work authors also mentioned the hybridization between Grey and African Grey-headed woodpeckers, describing it as rare in limited contact zones. Winkler et. al (1995) state that hybridisation occurs between all three. Although those woodpecker species differ in some habitat preferences, they’re all generally co-occur in humid woodlands with large trees and palms, but details on their hybridisation processes and biology are lacking.

### Genera Dryobates and Leuconotopicus

The *Dryobates* (five species) and *Leuconotopicus* (six species) genera consist of small arboreal species (Fuchs & Pons, 2015), which occur in the Palearctic (*D. minor*), Indo-Malayan (*D. cathpharius*) and Northern (*D. pubescens*, *D. nuttallii*, *D. scalaris, L. borealis, L. albolarvatus, L. arizonae, L. villosus*) and Central (*D. scalaris, L. fumigatus, L. stricklandi*) Americas. According to Ottenburghs & Nicolаї (2024), the North American *Dryobates* species have been confirmed as able to hybridise both with each other and with *Leuconotopicus villosus*.

Dixon (1989) summarised the knowledge on Downy *D. pubescens* and Ladder-backed Woodpeckers *D. scalaris*, noting that both species were closely related and overlapped in range in the Southern Great Plains of the USA, although having distinct foraging behaviour and habitat preferences. In contact zones, Downy occupies orchards and riparian woods, while Ladder-backed prefers open, xeric environments with mesquites, cacti and hackberry trees for foraging. In the Handbook of Avian Hybrids of the World (McCarthy 2006) the report by Sexton (1986) on apparent hybrids of these species in Texas was mentioned.

Downy Woodpecker also interbreeds with Nuttall’s Woodpecker D. nuttallii, the first presumed hybrid reported by Ridgway (1914). Short (1971) wrote of hybrids in the San Diego River Valley and emphasised that in that area the two species co-occurred, but that Nuttall’s was common and Downy rare. Unitt (1986) described another presumable hybrid in California that had intermediate vocalisations and many intermediate plumage features. The possibility of a backcross was suggested, as this hybrid mostly resembled Nuttall’s.

Nuttall’s x Ladder-backed Woodpecker hybrids in a contact zone in southern California (USA) and Northern Baja California (Mexico) were also mentioned by Short (1971). Nuttall’s prefers oak and riparian forests, while Ladder-backed inhabits arid, stunted wooded areas and deserts but they are sympatric in river valleys and the borders between dry and mesic forests. Despite differences in habitat, morphology (size, bill length and width), vocalisations and plumage features, hybrid individuals were recorded several times. Manthey, Boissinot & Moyle (2019) found low levels of introgression present in the phenotypically “pure” individuals that were in sympatric areas.

Mlodinow, et al. (2015) reported on apparent hybrids between Downy and Hairy (*D. villosus,* now placed in the genus *Leuconotopicus*) Woodpeckers in riparian forest in Colorado. The individual was described as having a call that resembled a Hairy, but with some plumage features of Downy. Overall size and bill length were intermediate between the two species.

Miller (1955) reported a presumed Hairy x Ladder-backed female hybrid. Although the geographical range of these species widely overlaps, the two have significantly different habitat preferences, as Hairy occupies montane coniferous forests and Ladder-backed prefers xeric areas. The hybrid was described as having plumage features, overall size and calls that were intermediate between the parental species.

### Genus Veniliornis

This genus currently includes 13 species distributed across different regions of South America, from which two species also entering Central America. All are small to medium in size, most with heavily barred underparts and green to yellow upperparts (red in some species) and sometimes with red on the head. Two species, previously included in the *Picoides* genus differ greatly, being black-and-white, with spotted underparts. Males have a red crown. Ottenburghs & Nicolаї (2024) reported one reliable and one uncertain case of hybridisation.

del Hoyo & Sargatal (2002) mention possible hybridising between Dot-fronted *V. frontalis* and Little *V. passerinus* Woodpeckers, – the species that occupy humid forests in the Andes in Bolivia and NW Argentina.

### Genus Dendrocopos

This genus currently includes 12 species, all found in the Palearctic and Indo-Malayan regions. They are medium-sized and have similar plumage, mostly black-and-white (pied) with red on the head and undertail. They occupy various forests and woodlands from primeval temperate to forest-steppe and anthropogenic woods (parks, orchards, gardens, etc.). Sexual dimorphism involves red on the head of males. According to Ottenburghs & Nicolaï (2024) only four species pairs had been confirmed to hybridise with a further three uncertain.

Hybridisation between Sind Woodpecker *D. assimilis* and Syrian Woodpecker *D. syriacus* has been confirmed in Eastern Iran (Vaurie 1959; Short 1982).

Great Spotted Woodpecker D. major is the most widespread species in this genus, occurring in most of the Palearctic and is known to hybridise with White-winged Woodpecker Dendrocopos leucopterus in northwestern China, where Vaurie (1959) considered the subspecies *D. major “tianshanicus”* as a hybrid form. There is known several cases of hybridisation between great spotted woodpecker and white-backed woodpecker *D. leucotos* in Scandinavia (Short 1982, Christensen 1999, Sarkanen and Koivusaari 2001), but likely more could be overlooked or published only in local journals (Aulén 1979, Dernfalk 1983).

Most hybrids, however, are great spotted woodpecker x Syrian Woodpecker *D. syriacus*, as the latter has expanded its range into Europe from the Middle East to Europe since the end of the 19^th^ century (Gorman, 1997). These two species were once considered to rarely hybridise, but recent studies on the morphology and genetic of both have found that it is, in fact, quite widespread Central Europe, especially in Poland and Czechia (Michalczuk, 2014). In addition, the Russian literature and the Internet, include cases of misidentified *Dendrocopos* birds that appear to be hybrids (Bakai et al, unpublished). Clearly, knowledge on hybridisation between these two close relatives has improved but is still far from complete. Hybrids have variable plumages, consisting of features from both parental species, but it is likely that appearance ultimately depends on the share of parental genomes, as *major x syriacus* hybrids are fertile and produce backcrosses (Figarski & Kajtoch, 2018). Furthermore, cases of mixed pairs involving two hybrid individuals that produced offspring have also been confirmed (Kajtoch & Kusal, 2022). In urban areas in SE Poland, about 5% of pairs were found to be mixed, with most formed by a female Syrian or female hybrid with a Great Spotted male (Figarski & Kajtoch, 2018). In the same study, hybrids constituted about 4% of all individuals observed and when dead birds were included, that value increased to 7%.

Recent genetic studies suggest that hybrids (including backcrosses) form up to 20% of sympatric urban populations (Michalczuk et al., 2014; Gurgul et al., 2019). Research on this subject concerning Syrian and Great Spotted Woodpeckers is ongoing.

### Genus Colaptes

The *Colaptes* genus includes 11 medium-sized species. These birds are heavily barred/spotted, with distinct back and abdomen coloration and bright contrasting colours only on underwing and undertail (sometimes head plumage has also distinct colouration). Sexual dimorphism is slight, generally expressed in malar stripe presence and/or colouration presence. Six species occur in S America, Fernandina’s Flicker *C. fernandinae* is endemic to Cuba, Gilded Flicker *C. chrysoides* is distributed around the Gulf of California and Northern Flicker *C. auratus* has a wide geographical range including N and C Americas and Cuba.

According to Ottenburghs & Nicolаї (2024), there one species pair hybridise. Many studies focus on hybridisation between two subspecies of Northern Flicker (*C. auratus*).

Ottenburghs & Nicolаї (2024) found information about Red-shafted Flicker *Colaptes auratus cafer x* Gilded Flicker hybridisation along a hybrid zone in Arizona (Johnson 1969). The differences in habitat preferences of two species are visible, with Gilded Flicker inhabit arid shrub and deserts at lowlads, and Red-shafted Flicker prefer more humid open land. However, both species could breed in the riverine woodlands, where they hybridise. Short (1982, p. 381) reports about limited hybridisation in a zone along one Arizona river and isolated or semi-isolated “hybrid swarm” populations in several river valleys of Arizona and in Baja California. Manthey et al. (2016) work provides evidence of genome admixture between two species.

### Genus *Celeus*

This genus currently includes 12 medium-sized, often brightly coloured, species. They inhabit varied environments, mostly across South America, with two in Central America. Sexual dimorphism is expressed by males having a red malar stripe. Chestnut-coloured *C. castaneus*, Chestnut *C. elegans*, Pale-crested *C. lugubris* and Blond-crested *C. flavescens* Woodpeckers are very highly similar morphologically (Short 1972). Despite these species being closely related, and co-occurring at the edges of their ranges, Short stated that, except rare cases involving Chestnut and Pale-chested, they do not hybridise.

According to Ottenburghs & Nicolаї (2024), there is one species pair confirmed to hybridise, and three species pair that may do so.

Two female hybrid individuals of Chestnut and Pale-chested were reported based on intermediate plumage characteristics and size (Short 1972; Kratter et al. 1992). The specimens were collected in the dry Chaco in Argentina as well as in humid Amazonian forest in Bolivia, thus mixed pairs hypothetically formed in the ecotone between these two habitat types.

### Genus Dryocopus

This genus of consists of six large species distributed in the Neotropics, Indo-tropics, Nearctic and Palearctic. Their plumage is basically black, occasionally with white and/or red on the head, and sometimes with a crest. Sexual dimorphism is slight, males usually being redder on head. Ottenburghs & Nicolаї (2024) mentions one pair of species in which hybridisation was confirmed.

Short (1982) stated that Black-bodied Woodpecker *D. schulzi* and Lineated Woodpecker *D. lineatus* hybrids are known from N Argentina, Paraguay and S Bolivia. The morphology of these hybrids is similar, except for Lineated which is bigger and whiter in its plumage. These two species barely differ in mitochondrial DNA (Shakya et al. 2017). The typical habitats used by the two species are different, with Black-bodied preferring xeric areas, but both meet in transitional areas.

### Genus Campephilus

The genus *Campephilus* includes 11 large species distributed across the Neotropics. They typically have a red head with a crest and black-and-white body plumage. Some are generalists and others are specialists and thus occupy diverse habitat types.

One species pair has been confirmed as hybridising (Ottenburghs & Nicolаї (2024)). C. leucopogon and C. melanoleucus are parapatric in Paraguay with the border between the two being along the Paraguay River. C. leucopogon inhabits dry Chaco woodlands while C. melanoleucus is less habitat-specific, occurring in semi-open areas, grasslands and gallery forests, but in contrast to its relative it prefers more humid, hill woodlands. Contreras, Chialchia & Smith (2014) found a male specimen in the IBIS collection (the University of Pilar, Ñeembucú Departament, Paraguay) that had plumage features of both species (head, chin) as well as intermediate ones (neck, belly).

### Genus *Picus*

This genus includes 15 large species that are distributed in diverse habitats throughout the Palearctic and Indo-tropic regions. They are typically green, males usually differing from females by having more red on the cap and in the malar stripe. According to Ottenburghs & Nicolаї (2024), two species pairs hybridise with one other possible.

Green Woodpecker *P. viridis* and Grey-headed Woodpecker *P. canus* overlap in range across much of the Western Palearctic. Both species most often co-occur at wooded areas, but hybridisation occurs only rarely. In known cases the hybrids were forming pairs with a pure parental bird, but the eggs were not fertilized (Ivanchev 1993). Hybrid individuals have variable plumage patterns, the features being most like one of parental species or intermediate (Gorman 2020). One of the most reliable features is the width of the malar stripe, but also a dark nape patch, although not all hybrids have the latter (Lawicki et al. 2015). Size, bill and silhouette are variable, but usually most resemble those of one of the parents. Unusual calls and drumming have also been reported (Czechowski & Bocheński 2012). It is thought that hybridisation occurs more often at the edge of geographical distribution of one of the species, as most of documented cases originate from such areas (Friedmann, 1993; Bird & Suedbeck, 2004; Lawicki et al., 2015). Friedmann (2011) noted that most hybrids were males. He also describes the dynamics of mixed pairs in the Moscow region, where during the range expansion of Grey-headed Woodpeckers between 1989-1998, mixed pairs, involving both sexes, were formed, but only the combination of Grey-faced females with Green males persisted for longer although only some of them succeeded to lay eggs. Friedmann (1993) explained the low rate of successful pairings by incompatibilities in mating behaviour between two species.

Green (*P*. *viridis*) and Iberian Woodpeckers (*P. sharpei*) were formerly considered conspecific. But recent molecular (Pons et al. 2011, 2019) and morphological differences (Olioso & Pons 2011) instead indicated that the two were separate species. The two hybridise along a 200 km wide zone in Southern France, in places where the range limits of both species merge and the densities of both are at their lowest. Information about this hybridisation case includes only genetic evidence of introgression of average 5-15 % introgressed loci genes (Pons et al. 2019) and variation in face patterns (Olioso & Pons 2011) among birds in the hybrid zone.

### Genus Dinopium

This genus currently includes five large tropical species distributed in S Asia, mostly in humid environments though some species are rather adaptable and synanthropic. All species are characterized by bright-coloured plumage, a black-and-white underbody, yellow or red wings and a prominent crest. Males differ from females by crest colouration. According to Ottenburghs & Nicolаї (2024), there is only one species pair that hybridises, but according to Freed et al. (2015) another pair may also do so.

Hybridisation between Sri Lankan Black-rumped Flameback Dinopium benghalence jaffnense and Lesser Sri Lankan Flameback Dinopium psarodes occurs in Central Sri Lanka and was studied by Freed, et al. (2015), Saminda, et al. (2016) and Ranasinghe et al. (2024). The first species occupies the northern part of the island and is believed to have colonized the island from Western Ghats, India (Freed et al. 2015). This first species has characteristic yellow wings and back. The second species is closely related to the former and used to be considered as a subspecies, however, being a more ancient coloniser, it diverged further, developing red colouration on its back and wings and a smaller body size. It occupies a bigger area in the southern half of the island. Hybrids tend to have different combinations of parental plumage features, including orange backs or wings. Mixed pairs are documented to form between two “pure” parental species, a parental species and a hybrid and between two hybrids, regardless of sex. Hybrids are reported to be most prevalent in the northern part of the hybrid zone, less so in the southern part of the zone (Freed et al. 2015; Saminda, et al. 2016).Recently, Ranasinghe et al. (2024) revised subspecies involved in this hybridization in northern Sri Lanka and revealed three-way hybridization involving the southern island-endemic red-backed *D. psarodes* and two genetically distinct populations of golden-backed *D. benghalense* originated the Indian form *D. b. tehminae*.

---

File S2. Characteristics of woodpecker subspecies with known records of hybridisation.

## Genus Picumnus

One presumed hybrid (AMNH 240125, American Museum of Natural History) between Plain-breasted Piculet P. castelnau and Fine-barred Piculet P. subtilis collected 1927 at Santa Rosa (Peru) on the upper Ucayali River was described by Short (1982). According to description, the hybrid has an intermediate features between two species in some plumage patterns (white spots on nape, crown coloration), but generally resembling Plain-breasted Piculet, thus it could be aberrant as well. There is little information on their biology, but both species seem to prefer humid Andean forests and have a contact zone at southeastern Peru.

## Genus Melanerpes

Barrowclough, et al. (2018) provide evidence for a genetically distinct haplotype of *Melanerpes carolinus perplexus* subspecies in the south of Florida, USA, which forms a hybrid zone with *M. c. carolinus* ssp. Morphologically, these subspecies can be distinguished by the presence of a tan-coloured band between the forehead and the bill in *perplexus*. Unfortunately, this study does not describe how phenotype correlates with haplotype in this woodpecker, nor gives the morphological characteristics of hybrids.

## Genus Campethera

Prigogine (1987) wrote on the two subspecies of Green-backed Woodpecker *C. c. cailliautii* and *C. c. permista* hybridising extensively in the NW of Lake Tanganyika, Tanzania. The *permista* subspecies is confined to lowland equatorial forest, whereas *cailliautii* prefers more-open habitats. The hybrid zone corresponds to an ecotone. However, in the case of subspecies it is unclear if it is a case of hybridisation or integration of populations (Rieseberg et al. 2004).

## Genus Colaptes

The two distinctive geographical races of the Northern Flicker *Colaptes auratus* (Yellow-shafted Flicker *C. a. auratus* x Red-shafted Flicker *C. a. cafer*), were previously considered as different species, but recently gained the status of subspecies due extensive hybridisation and little genetic divergence (Moore, Graham & Price 1991; Aguillon et al. 2018). Red-shafted Flicker occupies rocky land western from Great Plains while Yellow-shafted Flicker distributed eastern from it up to the Atlantic coast. The hybrid zone of two flicker ssp, being 4000km long and 300-500 km wide, spreading from Texas to Alaska (Wiebe & Moore 2008), where both taxa met at riparian woodlands. This hybrid zone has been stable in width for more than 100 years in the U.S. Great Planes (Moore & Buchanan 1985), although Aguillon & Rohwer 2022 indicated a recent shift of cline center associated with climatic changes. In the north-east part of the zone (Alberta, Canada) the shift caused by Red-shafted Flicker eastwards expansion was indicated at the middle of XX c. (McGillivray & Biermann 1987). The zone is believed to be maintained via bounded-hybrid superiority model when hybrid phenotypes performance is not different or higher than that of parental types in the ecotone (Moore & Koenig 1986; Flockhart & Wiebe 2009). The share of hybrids in population could be up to 95% (Anderson 1971). There are no differences between “pure” or hybrid phenotypes in annual survival, returning or recruiting rates in the northern part of a hybrid zone (British Columbia) where the populations are migratory (Wiebe & Bortolotti 2001; Flockhart & Wiebe 2008). Average clutch (8) and brood (6) sizes does not differ between phenotypes (Moore & Koenig 1986). Flockhart & Wiebe (2009) tested the prebreeding aggression level for the males of both phenotypes and the experiment shows no significant differences between them. There is also no evidence for these taxa to interbreed assortative at the major part of the hybrid zone (Bock 1971; Moore 1987), however, the flickers might do so in the northern part of the hybrid zone, where the breeding dates could differ between two geographical races (Wiebe 2000; Flockhart & Wiebe 2007). These flicker races are closely related and produce vital and fertile offspring, which freely interbred with parental phenotypes. The two flicker subspecies are mostly similar in morphology (Red-shafted Flicker is slightly bigger than Yellow-shafted), but with visible differences in plumage. The Red-shafted Flicker have a red malar strip in males, no red on nape and red underwings and undertail, while Yellow-shafted Flicker have black malar strip in males, red nape and yellow underwings and undertail. Short (1965) developed a hybrid index for identification based on plumage colouration. The morphometric analysis showed that few morphological features of hybrids were different from both parental ssp., but most of the features were closer or the same as in Red-shafted Flicker (Wiebe 2000). The genetic analysis shows that the flicker populations inside the hybrid zone contain various individuals with the different level of introgression – from F1 hybrids to highly advanced crossbreds of both ssp.. On average the Red-shafted Flicker have a later eggs laying dates than Yellow-shafted Flicker. The shaft colouration in hybrids vary from red and yellow (similar to parental) to intermediate orange (Wiebe & Bortolotti 2001), and they often have disrupted colouration of that feathers (Hudon, Wiebe & Stradi 2021). The habitat preferences of both subspecies generally the similar: both are present in the various open land, such as forest edges, riverbanks and farm tree-stands.

Short (1972) writes about hybridisation and introgression between some races within *Colaptes* species. Within Green-barred Woodpecker *C. melanochloros* there are two ecologically and morphologically distinct groups – the melanochloros (denser vegetation, more arboreal) and melanolaimus (open country, partly ground-foraging), which are hybridising freely in many areas.

Within Andean Flicker *C. rupicola* the topographically and ecologically limited hybridisation with introgression occurs between three distinct groups – *rupicola, puna,* and *cinereicapillus*.

Short (1972) also wrote about Campo flicker’s “two rather weakly marked races of the terrestrial *C. campestris*, northern campestris with a black throat and southern campestroides with a white throat. These races have come into secondary contact recently, and they interbreed in the only major area of contact, central Paraguay.”

---

File S3. Summary and characteristics of woodpecker species with records of hybridisation classified as “unreliable” according to Ottenburghs & Nicolаї (2024).

## Genus *Jynx*

The genera *Jynx* consists of two small woodpecker species. The Eurasian Wryneck *J. torquilla* is mostly migratory, breeding across the Palearctic and wintering in sub-Saharan Africa and southern Asia. The Red-throated Wryneck *J. ruficollis* is resident in central and southern Africa (Gorman 2022).

A note by Desfayes (1969) on a presumed hybrid individual between Eurasian and Red-throated Wrynecks, collected in Tanzania, detailed that the bird had some intermediate plumage features. However, Ottenburghs & Nicolаї (2024), following a re-examination by Short & Bock (1972) state that the bird was merely an atypical, aberrant, Red-throated Wryneck.

## Genus Picumnus

There are 19 small woodpecker species in this genus, most occurring in Central and South America. All species show preferences for open areas with low but dense vegetation, that is, shrubs and understory. Sexual dimorphism is minimal, males differing from females in having a yellow, orange or red spot on the forehead. Ottenburghs & Nicolаї (2024) reports about two doubtful hybridization cases.

The possibility of hybridisation between Grayish Piculet *P. granadensis* and Olivaceous piculet *Picumnus olivaceus* is only presumed as these closely related species overlap in range in the Cauca Valley in the NW Andes in Colombia. Both species occur at low elevations in open areas with abundant dense short vegetation (shrubs, understory, forest edges), although Grayish Piculet prefers drier environments. Nevertheless, there are no conformed records of hybridisation.

Short (1982) reported on a presumed Arrowhead Piculet *P. minutissimus* and White-bellied Piculet *P. spilogaster* hybrid in the collection of the British Museum. In theory, these two species could co-occur in clearings, mangroves and at forest edges.

## Genus *Sasia*

This genus consists of two small Indo-tropical species with bright yellow-green plumage. Sexual dimorphism is minimal, males differing from females in having a yellow forehead. The species distribution ranges overlap in Malaysia.

Ottenburghs & Nicolаї (2024) state that one individual with intermediate features between White-browed Piculet *S. ochracea* and Rufous Piculet *S. abnormis* was mentioned by Handbook of Avian Hybrids of the World McCarthy (2006), and that Short (1982) mentioned the possibility of both species co-occurring in Myanmar and Thailand. Hybridisation between these two species is likely, as they have similar habitat preferences, diet and nesting habitats and their breeding seasons overlap. Yet, there was not documented case of hybridisation so far.

## Genus Campethera

The genera *Campethera* and *Pardipicus* include ten African woodpecker species that mostly have white bodies, green wings, back and tail and a red nape. Sexual dimorphism is expressed in males having a red crown and malar stripe. According to Ottenburghs & Nicolаї (2024) there is three species pairs with uncertain hybridisation.

Short (1982) and Winkler (1995) reported on about apparent hybrids between Mombasa Woodpecker *C. mombassica* and Golden-tailed Woodpecker *C. abigoni* in eastern-central Tanzania. These two species can co-occur in coastal forests and have similar feeding behavior, but more studies are needed to confirm whether they hybridise or not.

Two other doubtful cases of hybridisation in this genus are described in McCarthy (2006), between Reichenow’s Woodpecker *C. scriptoricauda* and Bennett’s Woodpecker *C. bennettii* and between Nubian Woodpecker *C. nubica* and Fine-spotted *C. punctuligera* Woodpecker. The following species occupies different parts of the African continent but have overlapping zones between pairs closer to the continent’s center. The population of Reichenow’s Woodpecker in the area between Tanzania and Mozambique described by the author as “having intermediate plumage features between present species and Bennett’s Woodpecker, thus the birds might be a hybrid population”. The same author says about Fine-spotted Woodpecker’s *balia* subspecies in S Sudan and NE DR Congo, which geographical range overlaps with Nubian Woodpecker. Details on biology are scarce but in general these pairs of species have similar diet, occurs in common open land habitats and their populations in the range of overlap have similar breeding period.

## Genus Chloropicus

The three woodpeckers in this genus inhabit different parts of sub-Saharan Africa. They are generally green in colour with white striped heads. They occur in different types of forests and open woodland.

According to Ottenburghs & Nicolаї (2024), Short (1982) states that Sterling’s Woodpecker *Dendropicus stierlingi,* which occurs around Lake Malawi, is intermediate morphologically, vocally and behaviourally between Bearded Woodpecker *C. namaquus* and Cardinal Woodpecker *Dendropicus fuscescens*. The three species meet in the same geographical area around Lake Malawi and potentially could all occur in the same riparian habitats.

## Genus Dendropicos

This genus consists of twelve small to medium-sized species distributed across much of sub-Saharan Africa, although they occupy different habitats. They are coloured from yellow to gray with green wings and red crowns in males. Ottenburghs & Nicolаї (2024) listed one doubtful case of hybridization in this genus.

Short (1982) and Winkler (1995) state that the northern subspecies of Gabon Woodpecker *D. gabonensis reichenowi* has an intermediate morphology between the nominate of Gabon and Melancholy *D. lugubris* Woodpecker, but that hybridisation was not proven. Gabon subspecies *reichenowi* and Melancholy Woodpecker do co-occur, and the habitats they occupy are similar, but in-depth details on their biology are lacking.

## Genus Veniliornis

This genus consists of 13 species distributed in different parts of South America, and two in Central America. They are small to medium-sized, most with strongly barred lower bodies and green to yellow upper bodies, though some are reddish, and sometimes have red on the head. Two species that previously belonged to the *Picoides* genus differ greatly in plumage, being black and white, with spotted lower bodies. Males have red crowns. Ottenburghs & Nicolаї (2024) write that there is a one unreliable case of hybridization in this genus.

Goodwin (1968) mentions unusual Striped Woodpecker *V. lignarius* specimens in the British Museum which originated from Cordova, Argentina, that have unusually long red nape patches that recall Chequered Woodpecker’s *V. mixtus*. These two species occupy similar moist wooded habitats but tend to differ in altitudinal preference.

## Genus Dendrocopos

There are 12 species currently recognized. They are medium-sized, and all have similar plumage (black and white with red on the undertail. They occupy various forests and woodlands from primeval temperate to forest-steppe and anthropogenic woods (parks, orchards etc.). The usual sexually dimorphic feature is that males have red on the head.

According to Ottenburghs & Nicolaï (2024) for three pairs of species in this genus cases of doubtful hybridization were reported.

Goodwin (1968) and Short (1982) mention intermediate individuals between two SE Asian species that were until recently considered conspecific, Spot-breasted Woodpecker *D. analis* and Fulvous-breasted Woodpecker *D. macei*. Goodwin (1968) also mentioned intermediate individuals between Stripe-breasted Woodpecker *D. atratus* and Fulvous-breasted in India. However, no specimens or photographs of these potential hybrids exist.

## Genus Piculus

A genus of seven medium-sized woodpeckers that occur in humid woodlands in Central and Southern America. The typical plumage pattern is a white or yellow body with dense black barring, green back, wings and tail and yellow and/or red on the head. Males differ from females in having more red on the head and in most species also a red malar strip. Ottenburghs & Nicolaï (2024) mention one case of hybridisation that was not proven.

McCarthy (2006) describes the Stripe-cheeked Woodpecker *P. callopterus* as geographically and morphologically intermediate between the Rufous-winged Woodpecker *P. simplex* and Lita Woodpecker *P. litae*. Although Short (1982) mentions that these species overlap in distribution, according to the distribution maps of BirdLife International (2024) these two woodpeckers are hardy overlap in the C Panama, while the Lita woodpeckers’ geographical range does not overlap with neither of these species. The assumption of Stripe-cheeked Woodpecker to be a hybrid between other two species is very unlikely, and the morphological intermediacy is rather result the geographical gradient.

## Genus *Celeus*

This genus consists of twelve medium-sized, often brightly coloured species. They inhabit various environments, mostly in South America, but two species found in Central America. Sexual dimorphism is expressed by a red malar stripe in males. Chestnut-coloured *C. castaneus*, Chestnut *C. elegans*, Pale-crested *C. lugubris* and Blond-crested *C. flavescens* Woodpeckers are considered to be allospecies due to their high morphological similarities (Short 1972). Despite them being closely related, and co-occurring at the edges of their distributions, Short states that except for rare cases concerning Chestnut and Pale-crested Woodpeckers, they do not hybridise.

According to Ottenburghs & Nicolаї (2024), there are three species pairs which hybridization is doubtful.

Ottenburghs & Nicolаї (2024) found the inter-genera Chestnut-coloured Woodpecker and Golden-olive Woodpecker *Colaptes rubiginosus* interbreeding case mentioned in McCarthy (2006) to have involved a mistakenly identified individual. Although both of these species occur in Central America, and they could potentially occupy the same habitat types, hybridisation between them seems highly unlikely.

Another two examples of doubtful pairings mentioned by authors concern Ringed Woodpecker *C. torquatus* x Blond-crested Woodpecker and Ringed Woodpecker x Cream-coloured Woodpecker *C. flavus.* These hybridisation cases are listed on a hybrid-birds database (http://www.bird-hybrids.com) with incorrect references.

## Genus *Picus*

This genus consists of 15 fairly large woodpeckers that are distributed in diverse habitats throughout the Palearctic and Indotropic region. They are mainly green in colour, with males differing from females by having more red on the head.

According to Ottenburghs & Nicolаї (2024), there is one species pair with uncertain hybridisation.

Short (1982) mentions a case of possible interbreeding between Lesser Yellownape *P. chloropus* and Streak-throated Woodpecker *P. xanthopygaeus* in Sri Lanka, referring to Phillips (1953). Both species have flexible ecologies and occupy various types of forests and so can co-occur, however their morphology differs greatly. Ottenburghs & Nicolаї (2024) mentions that the original paper was unavailable.

## Genus Dinopium

This genus consists of five large tropical woodpecker species that are distributed across S Asia, mostly in humid environments, though some species are synanthropic. They have bright plumage, white and black on the body, yellow or red in the wings and on the back and have prominent crests. Males differ from females in crest coloration. According to Ottenburghs & Nicolаї (2024), there is only one species pair unconfirmed to hybridise.

Short (1982) writes that Himalayan Flameback *D. shorii* and Common Flameback *D. javanence* overlap in C and SW Burma, and a presumed hybrid from the two is in the British Museum. Despite the similar geographical range, the habitat preferences of the two species are quite different, which means fewer opportunities for co-occurrence. The morphology of Himalayan and Common Flamebacks is quite similar.

---

**Figure S1.**
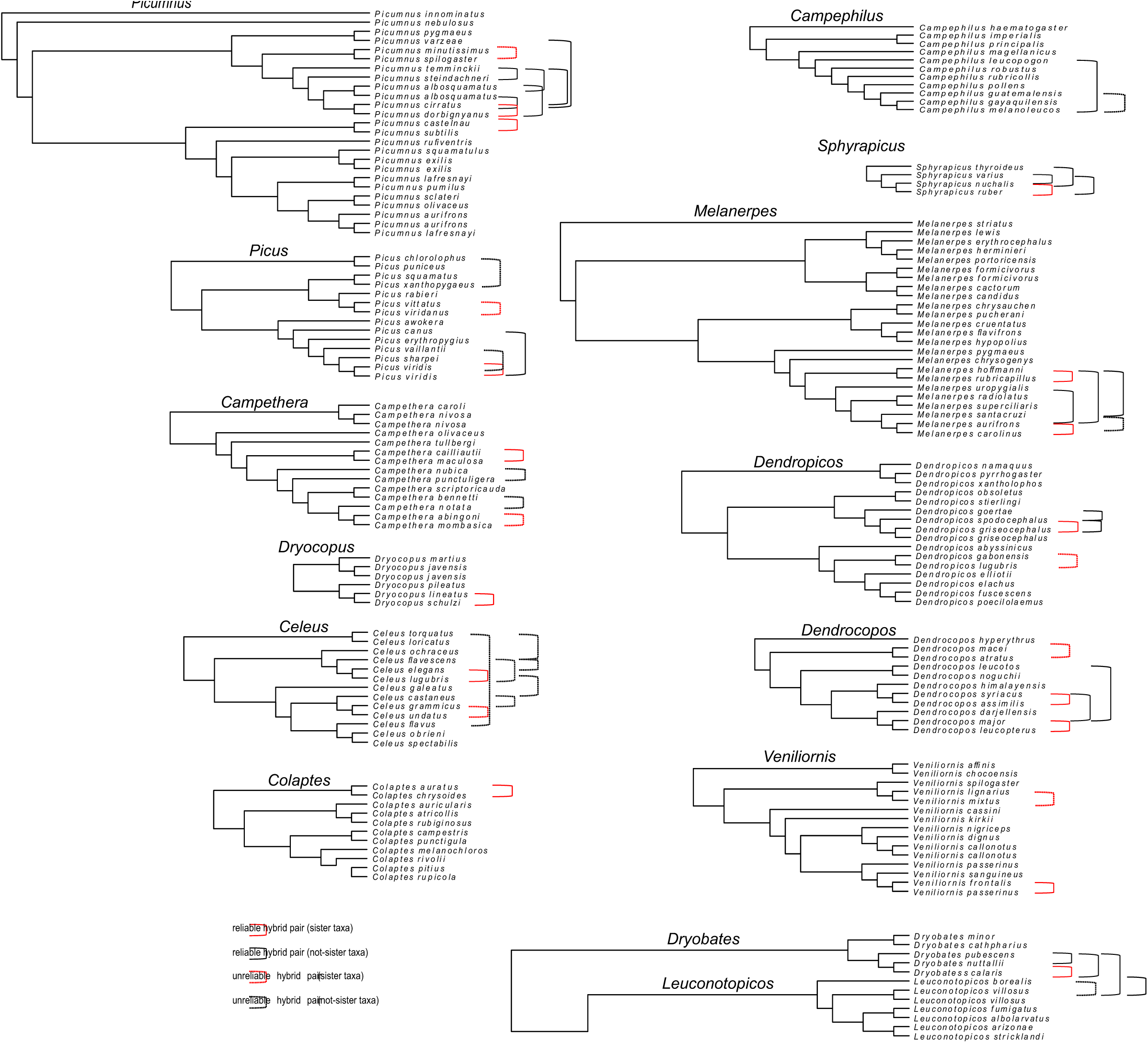
Phylogenetic trees of woodpecker genera (based on Shakya et al. 2017), that include hybridising species (according to Ottenburghs & Nicolaï 2024) with marked species pairs with confirmed hybridisation and those for which interspecific mating and breeding is uncertain.

**Table S1.**
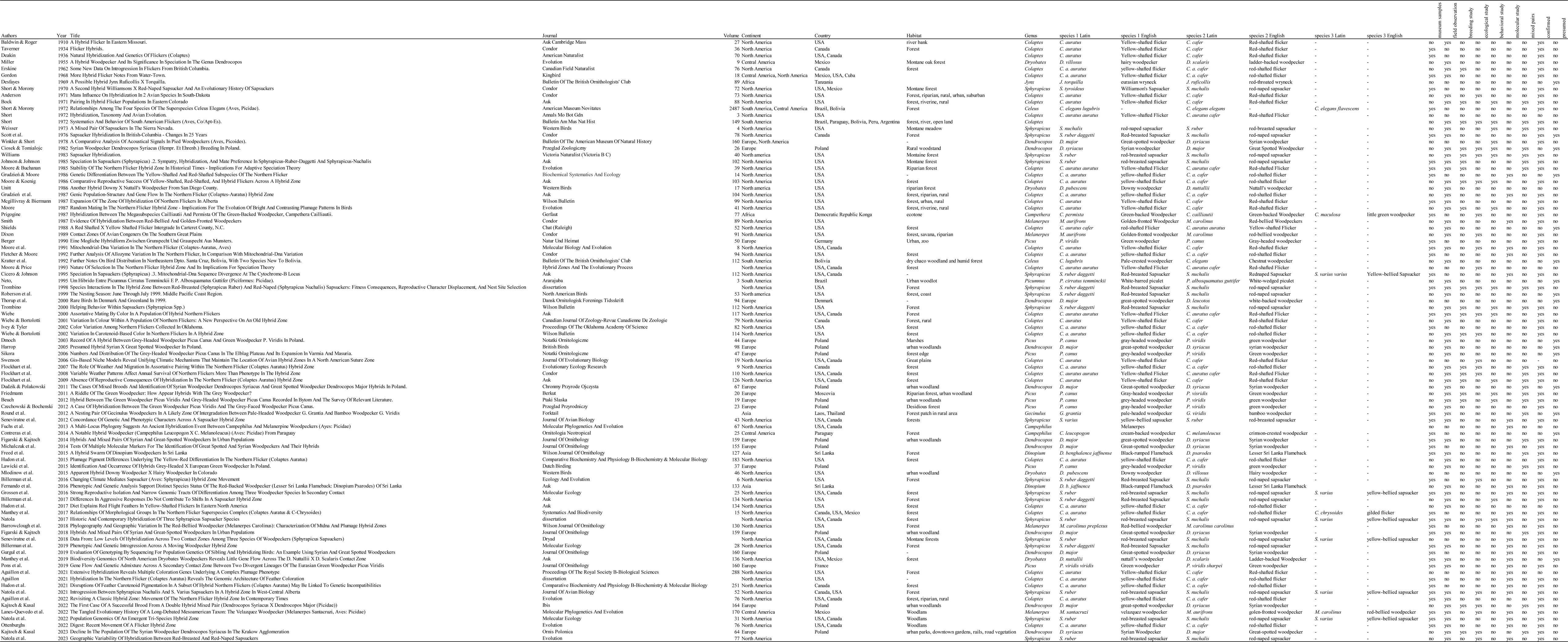
Summary of data collected from selected articles dealing with hybridization of woodpeckers.

**Table S2.**
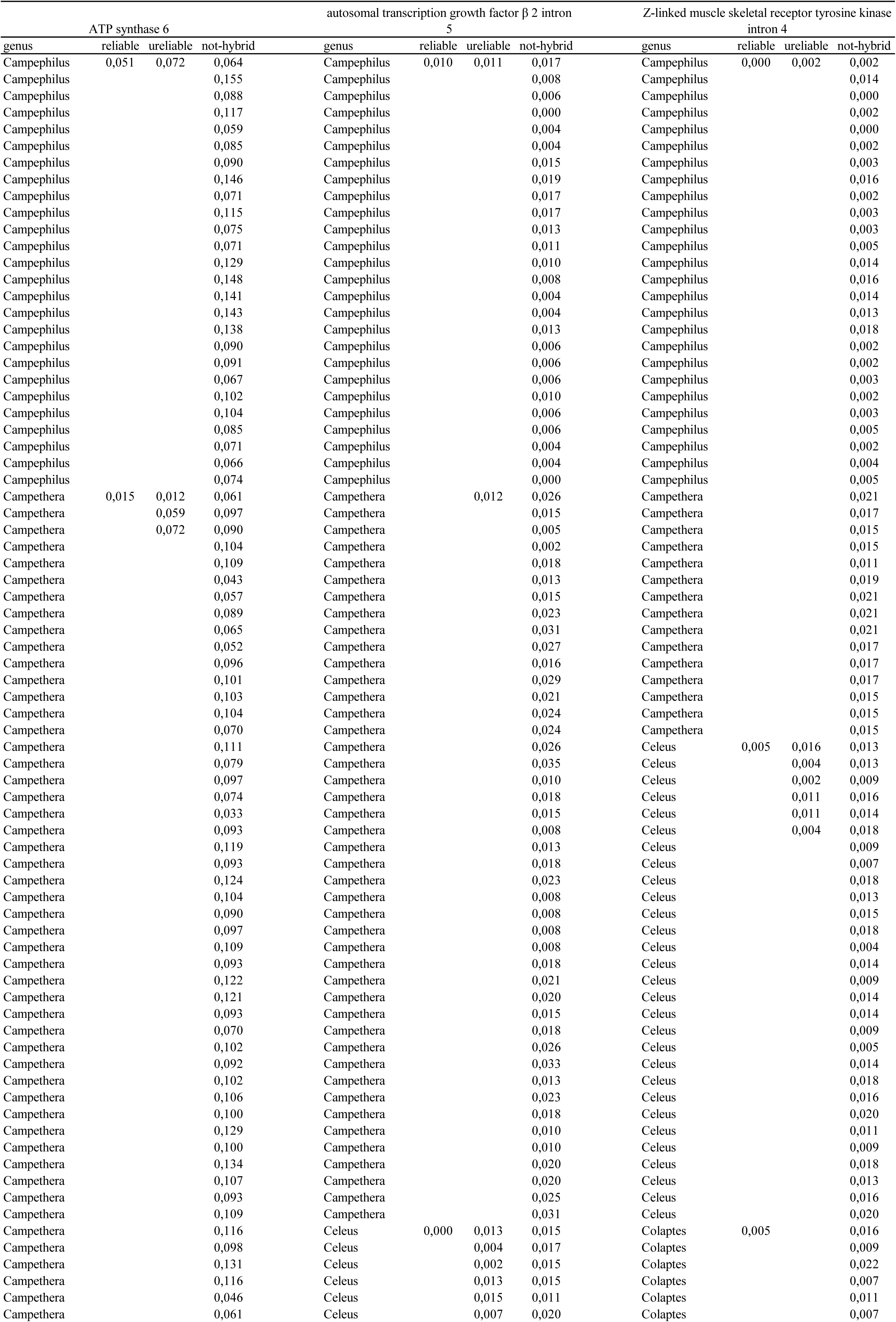
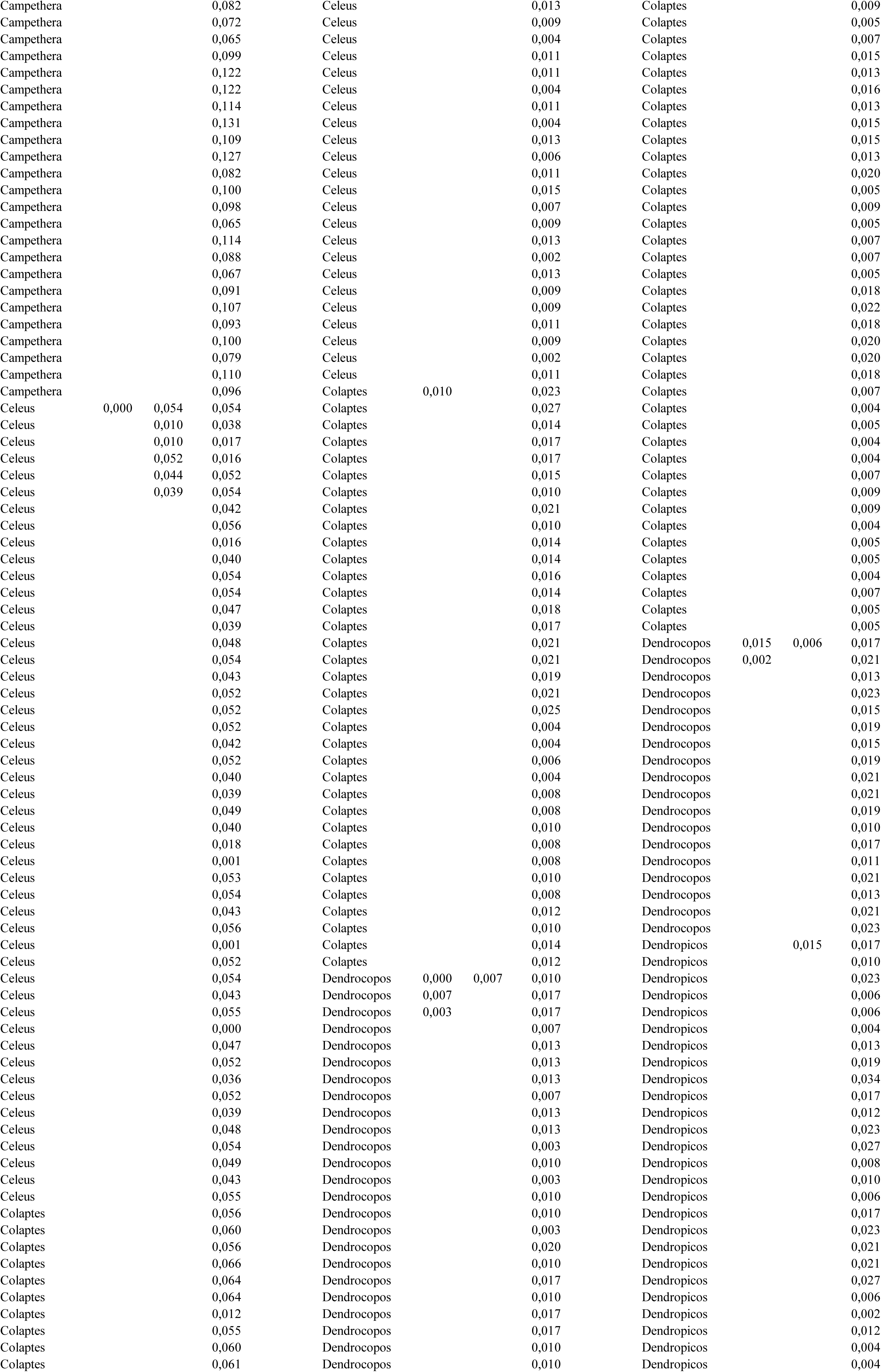
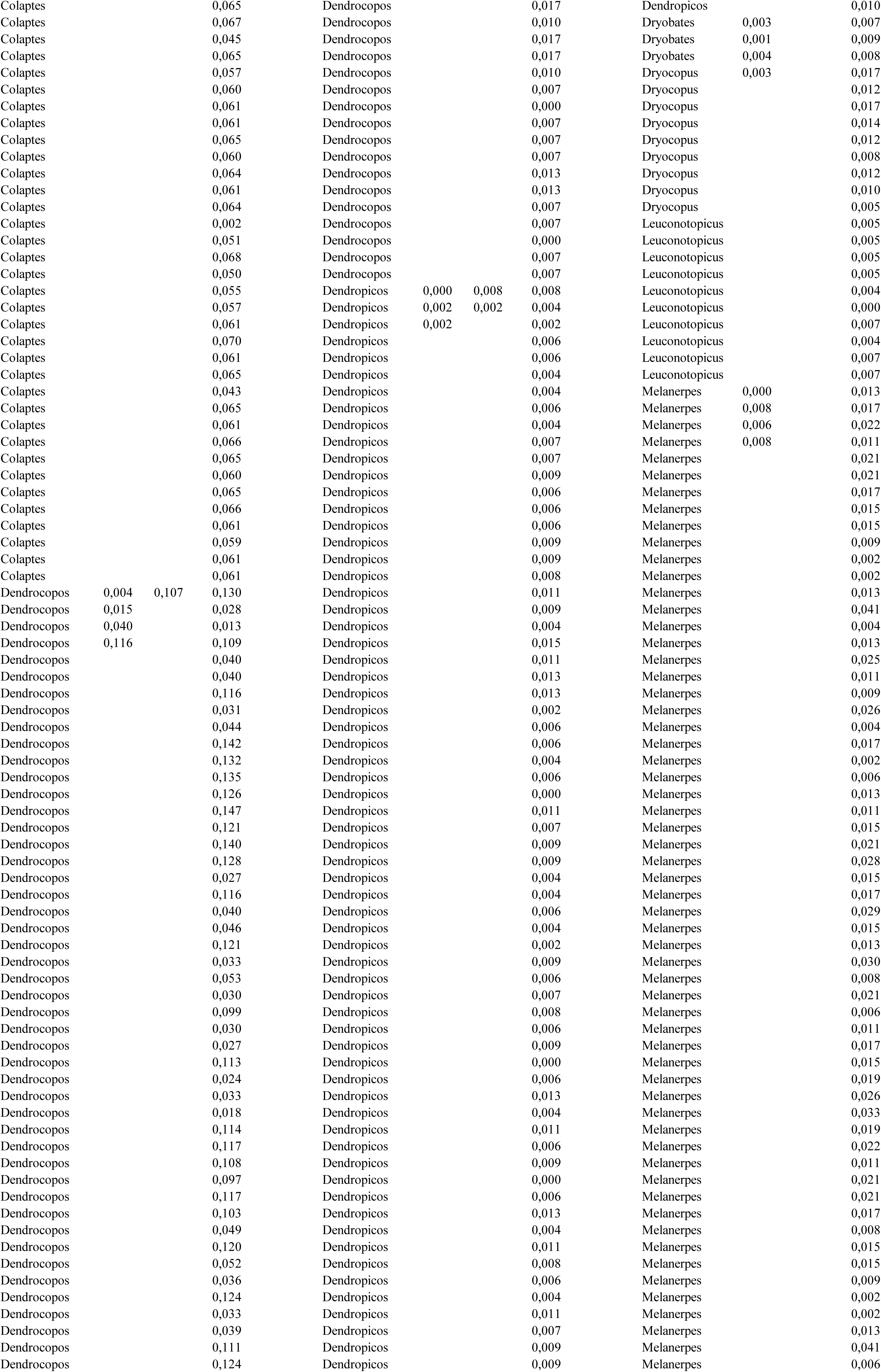
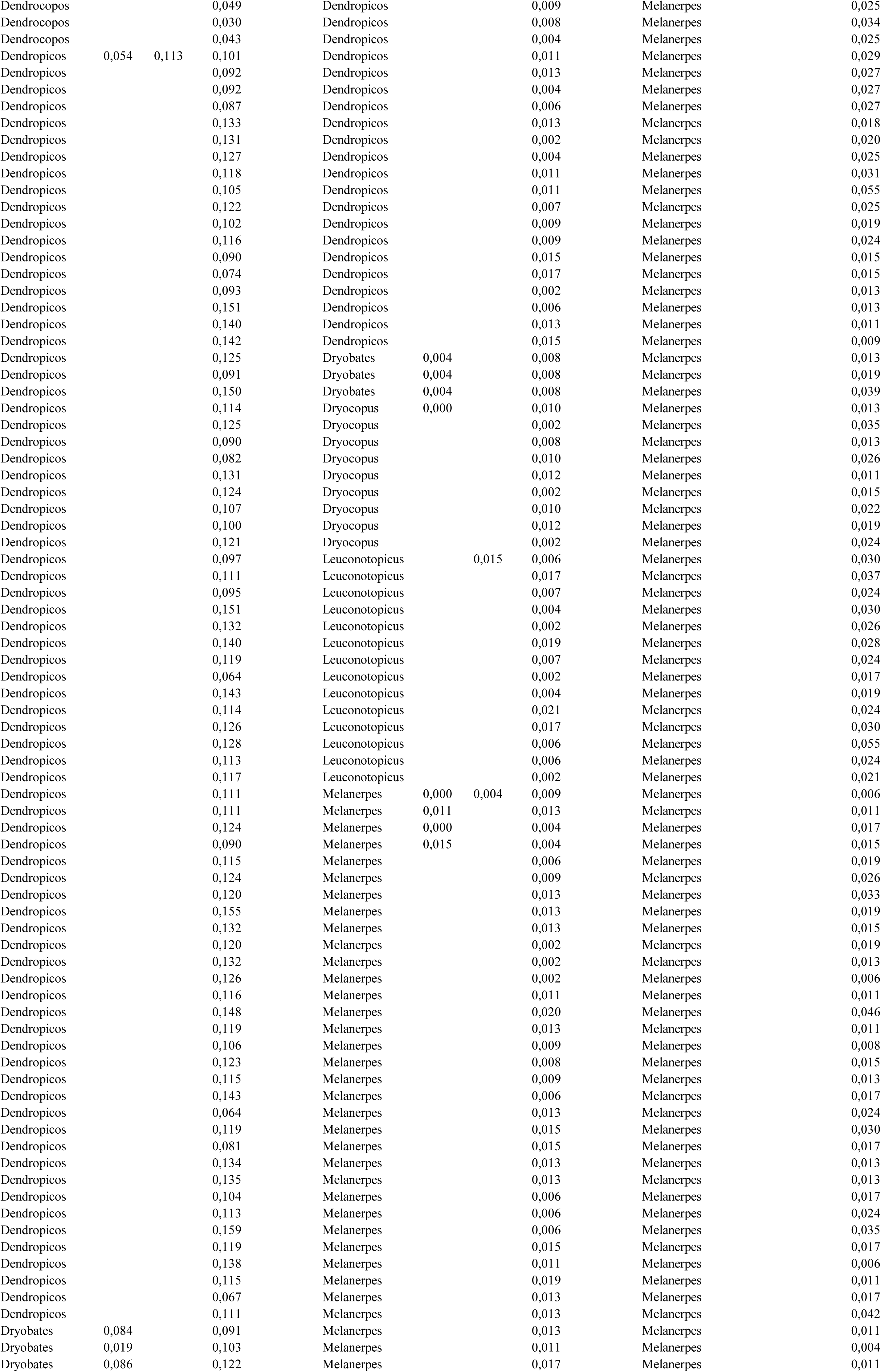
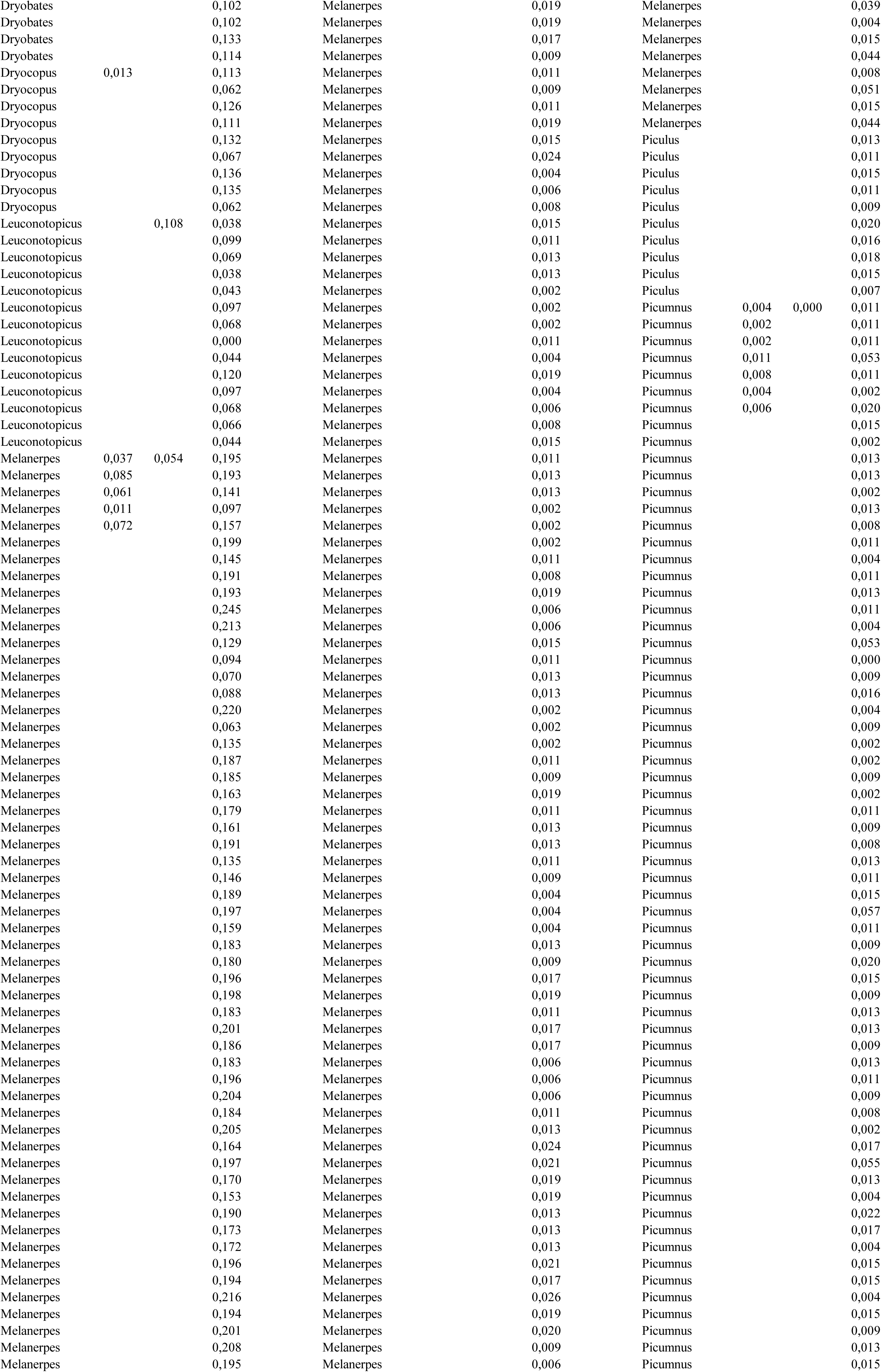
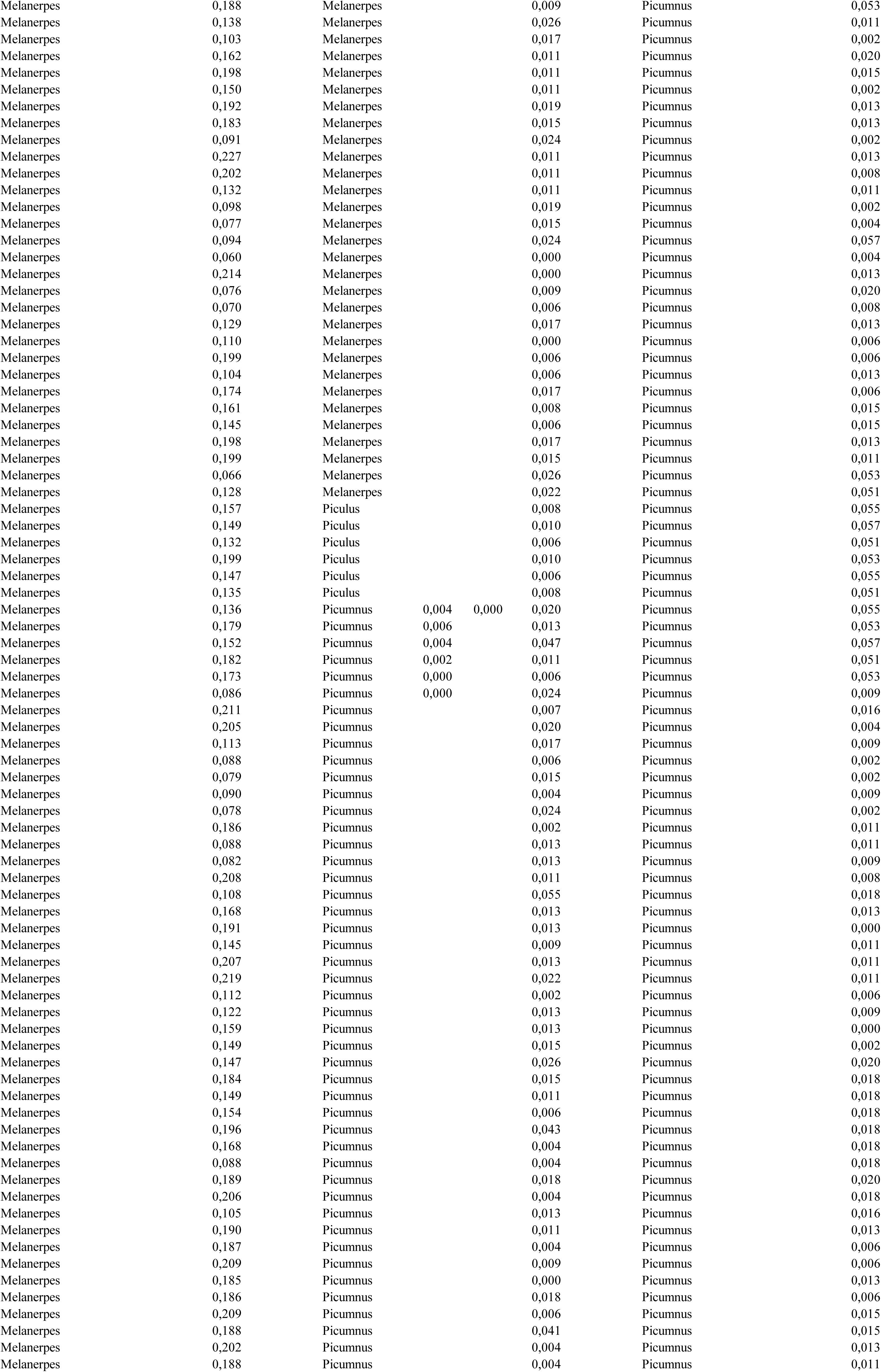
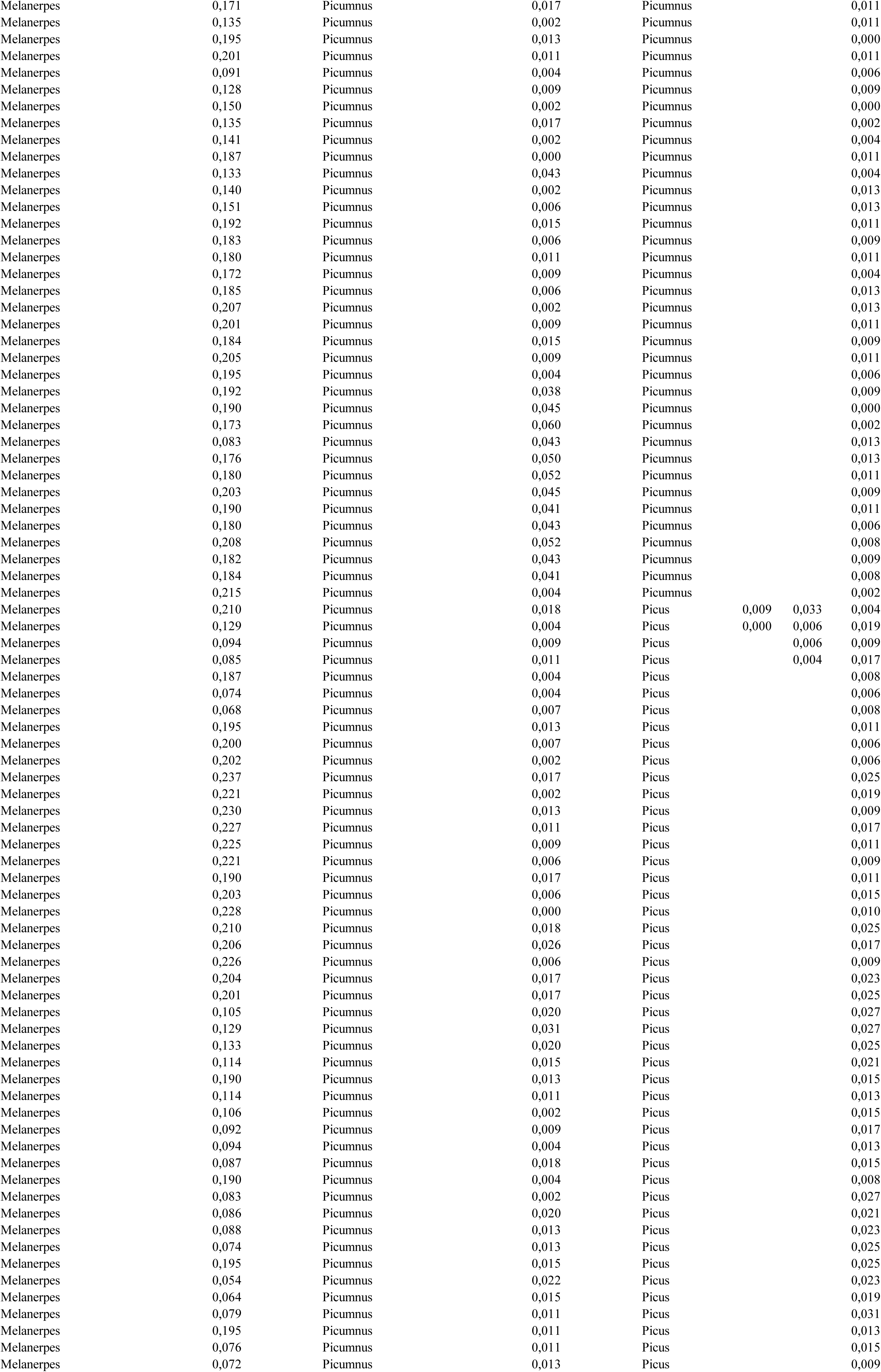
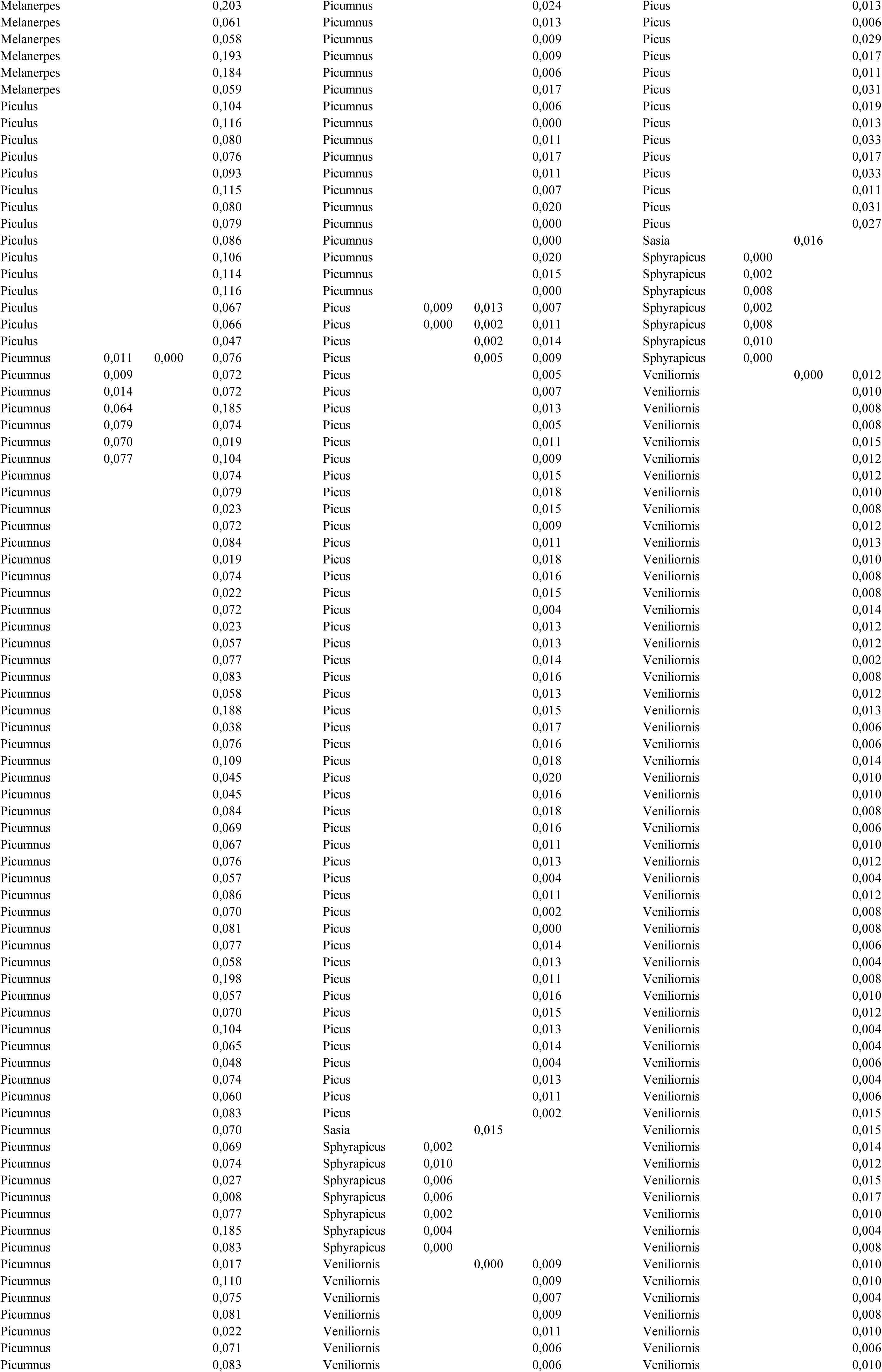
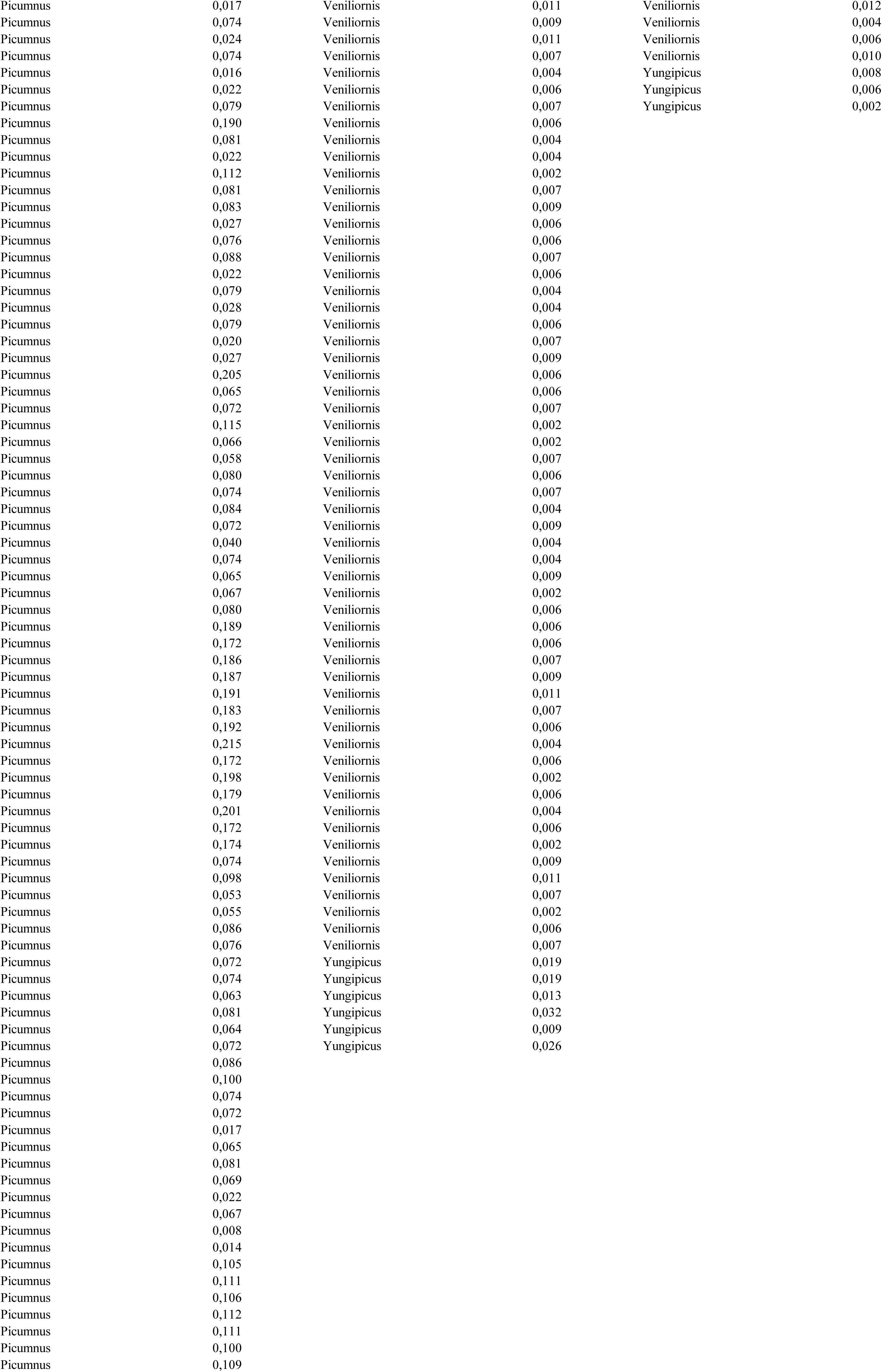
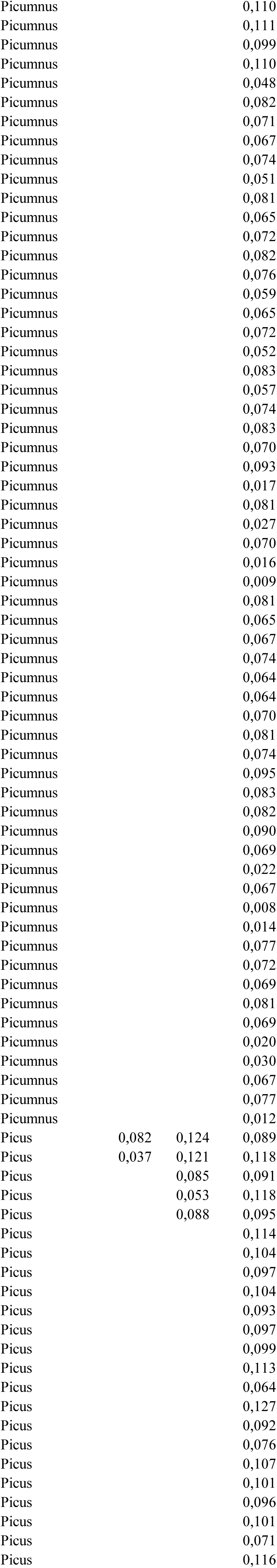
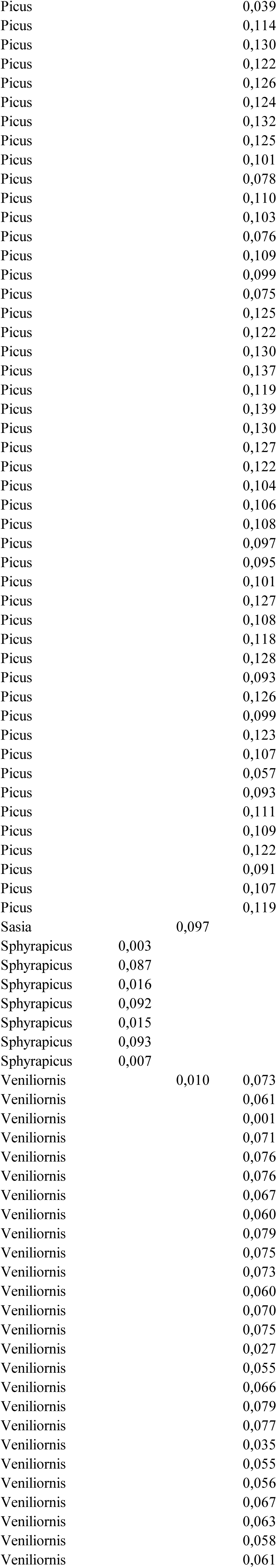
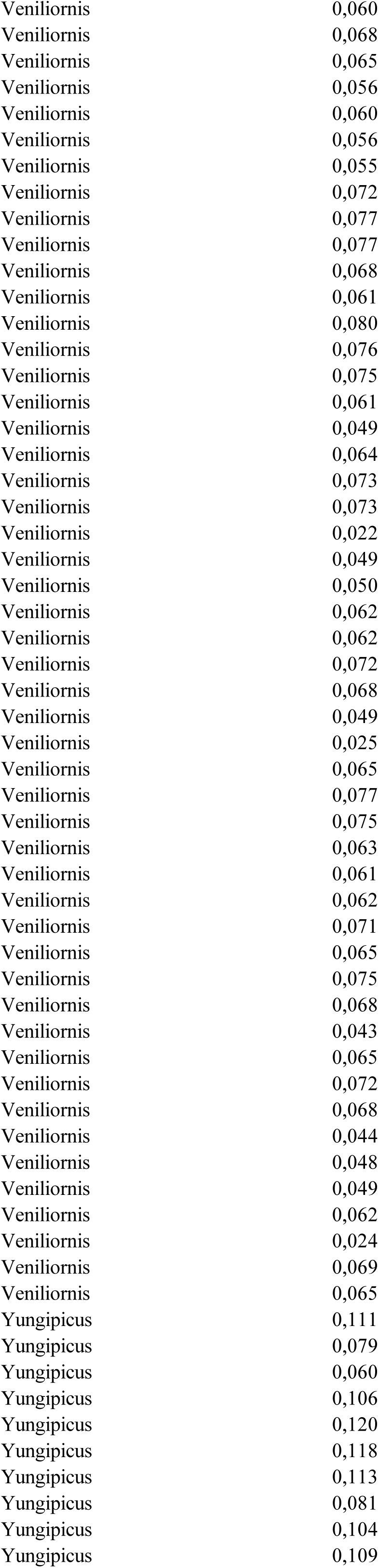
Pairwise genetic distances calculated for three genetic markers between pairs of woodpeckers (only intrageneric) that are known to hybridize, unreliable cases and not-hybridizing.

**Table S3.** Basic statistics describing paiwise genetic distances calculated for three molecular markers ATP synthase 6 (ATP); autosomal transcription growth factor β 2 intron 5 (TGF) and Z-linked muscle skeletal receptor tyrosine kinase intron 4 (MUSK) and three groups of woodpecker pairs (hybridizing, unreliable and not-hybridizing).

| Statistics | species pairs | Average | Median | Min | Max | Standard Deviation |
| --- | --- | --- | --- | --- | --- | --- |
| ATP | reliable hybrids | 0.046 | 0.040 | 0.000 | 0.116 | 0.034 |
|  | unreliable hybrids | 0.063 | 0.057 | 0.000 | 0.124 | 0.038 |
|  | not-hybrids | 0.101 | 0.091 | 0.000 | 0.245 | 0.051 |
| TGF | reliable hybrids | 0.004 | 0.004 | 0.000 | 0.015 | 0.004 |
|  | unreliable hybrids | 0.008 | 0.007 | 0.000 | 0.015 | 0.005 |
|  | not-hybrids | 0.011 | 0.010 | 0.000 | 0.060 | 0.008 |
| MUSC | reliable hybrids | 0.005 | 0.004 | 0.000 | 0.015 | 0.004 |
|  | unreliable hybrids | 0.008 | 0.006 | 0.000 | 0.033 | 0.008 |
|  | not-hybrids | 0.015 | 0.013 | 0.000 | 0.057 | 0.011 |

**Table S4.**
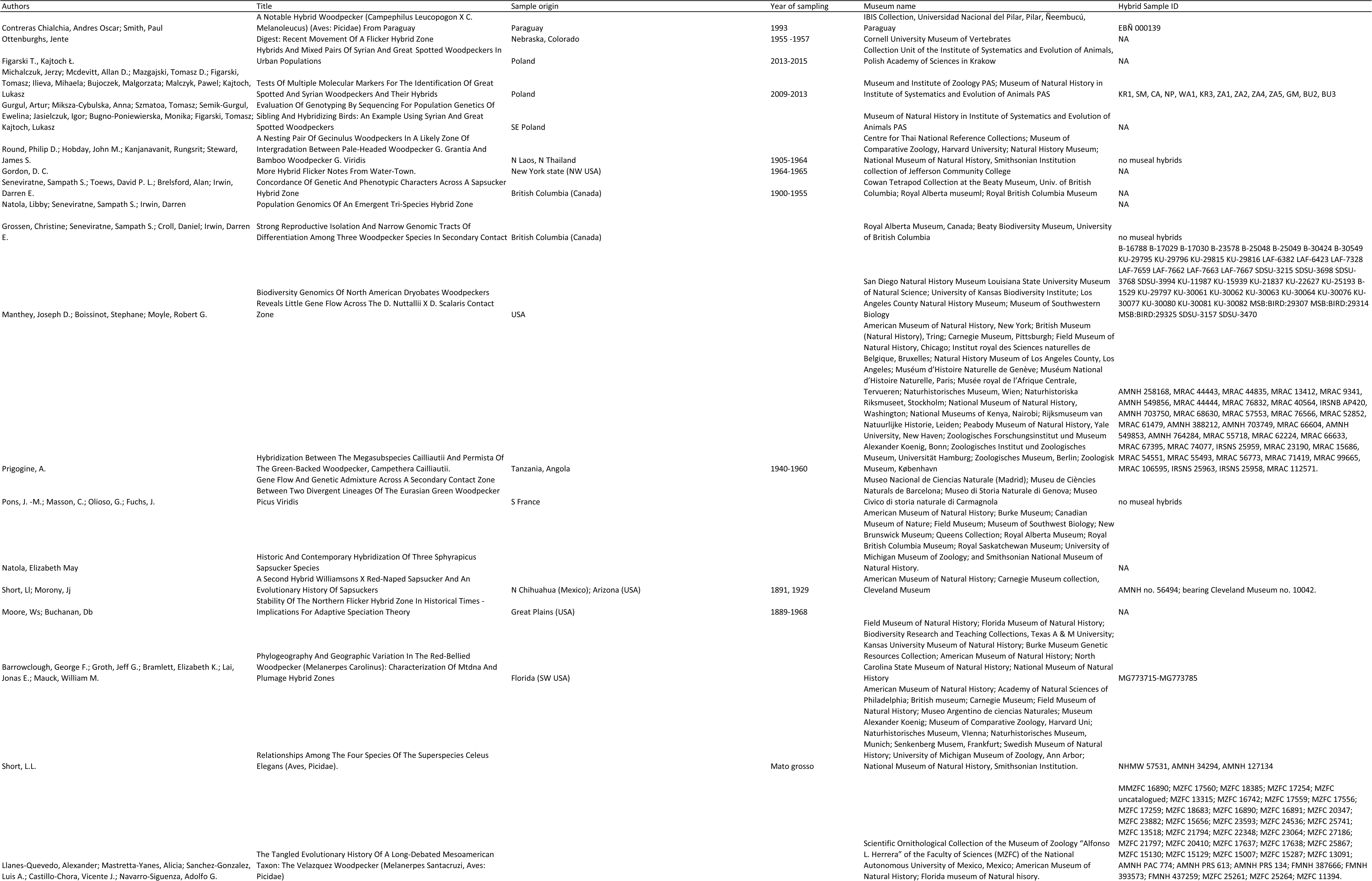
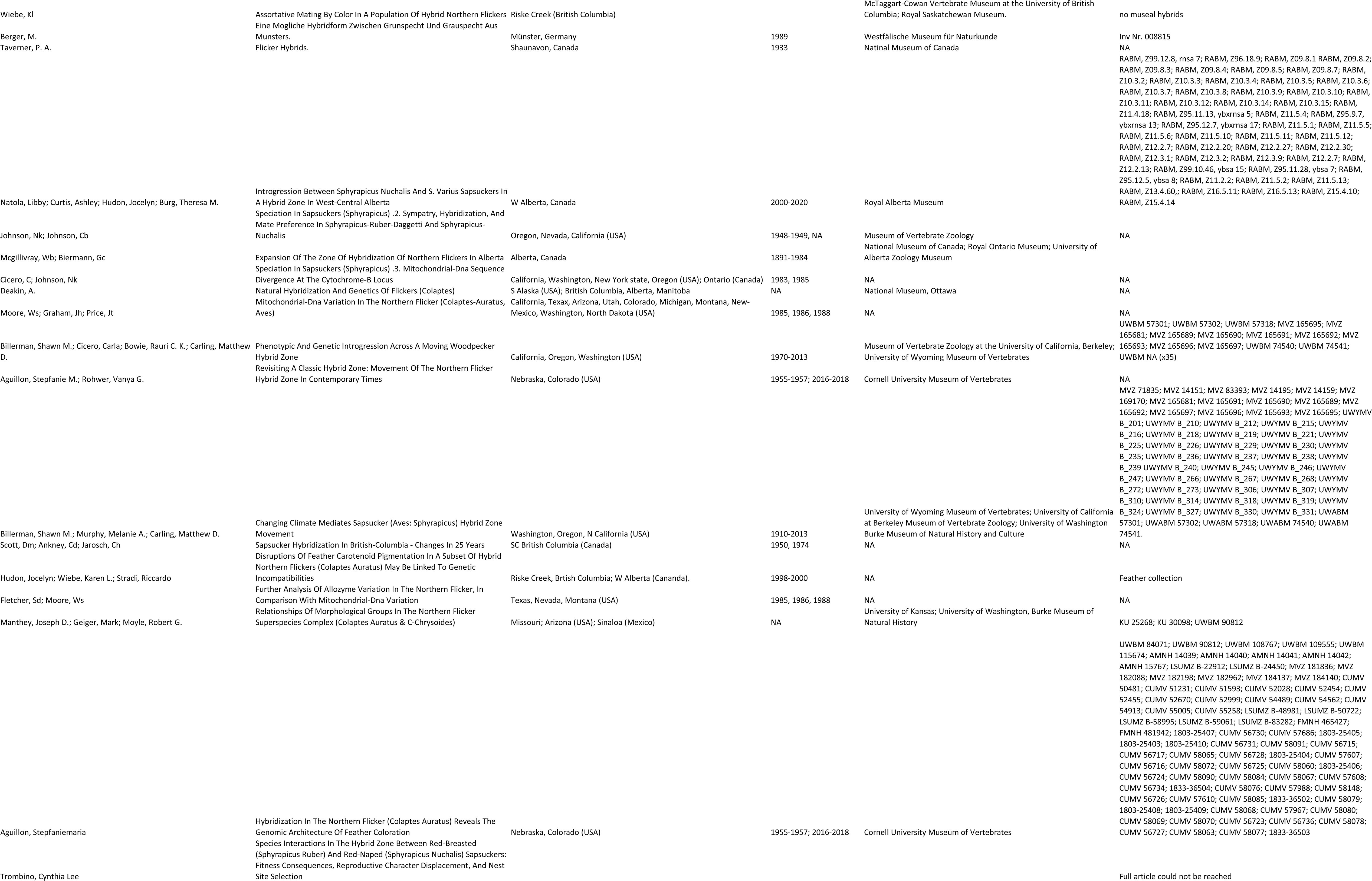
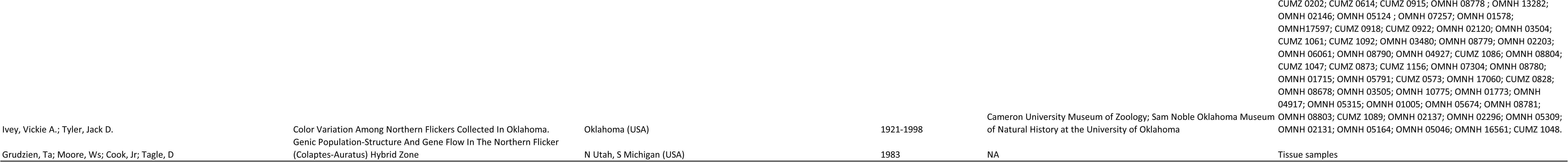
Summary of collection sites and woodpecker’s voucher numbers found in the literature in search for publications dealing with hybridization.

**Table S5.** Divergence time (according to data from Sakhya et al. 2017) and numbers of studies on hybrid woodpeckers

| Species1 | Species2 | Divergence time | Hybrid evidence (number of studies) |  |  |
| --- | --- | --- | --- | --- | --- |
|  |  |  | Field/photography | Museal sample | Genetic data |
| *Picumnus albosquamatus* | *Picumnus cirratus* | 0,6 | 2 |  |  |
| *Picumnus albosquamatus* | *Picumnus dorbygianus* | 0,6 | 1 |  |  |
| *Picumnus albosquamatus* | *Picumnus temminckii* | 1 | 1 |  |  |
| *Picumnus castelnau* | *Picumnus subtilis* | 1,4 | 1 |  |  |
| *Picumnus cirratus* | *Picumnus dorbygianus* | 0,4 | 1 |  |  |
| *Picumnus cirratus* | *Picumnus temminckii* | 1 | 1 |  |  |
| *Picumnus cirratus* | *Picumnus varzeae* | 1 | 1 |  |  |
| *Melanerpes aurifrons* | *Melanerpes carolinus* | 1 | 2 |  | 1 |
| *Melanerpes aurifrons* | *Melanerpes hoffmannii* | 3,2 | 2 |  |  |
| *Melanerpes aurifrons* | *Melanerpes santacruzi* |  |  | 1 | 1 |
| *Melanerpes aurifrons* | *Melanerpes uropygialis* | 2,5 | 3 |  |  |
| *Melanerpes hoffmannii* | *Melanerpes santacruzi* | 3,2 | 1 |  |  |
| *Melanerpes hoffmannii* | *Melanerpes rubricapillus* | 0,4 | 1 |  |  |
| *Shyrapicus nuchalis* | *Spyrapicus ruber* | 0,3 | 11 | 8 | 8 |
| *Shyrapicus nuchalis* | *Spyrapicus thyroides* | 5,1 | 1 | 1 |  |
| *Shyrapicus nuchalis* | *Spyrapicus varius* | 0,8 | 5 | 5 | 6 |
| *Spyrapicus ruber* | *Spyrapicus varius* | 0,8 | 4 | 5 | 6 |
| *Campethera cailliautii* | *Campethera maculosa* | 0,5 | 1 |  |  |
| *Dendropicos goertae* | *Dendropicos griseocephalus* | 3,1 | 2 |  |  |
| *Dendropicos goertae* | *Dendropicos spodocephalus* | 3,1 | 2 |  |  |
| *Dendropicos griseocephalus* | *Dendropicos spodocephalus* | 1,4 | 1 |  |  |
| *Dryobates nuttallii* | *Dryobates pubescens* | 3 | 3 |  |  |
| *Dryobates nuttallii* | *Dryobates scalaris* | 0,6 | 1 | 1 | 1 |
| *Dryobates pubescens* | *Dryobates scalaris* | 3 | 1 |  |  |
| *Dryobates pubescens* | *Leuconotopicus villosus* | 6,5 | 1 |  |  |
| *Dryobates scalaris* | *Leuconotopicus villosus* | 6,5 | 1 |  |  |
| *Veniliornis frontalis* | *Veniliornis passerinus* | 1,5 | 1 |  |  |
| *Dendrocopos assimilis* | *Dendrocopos syriacus* | 0,3 | 2 |  |  |
| *Dendrocopos leucopterus* | *Dendrocopos major* | 0,6 | 1 |  |  |
| *Dendrocopos leucotos* | *Dendrocopos major* | 1,9 | 6 | 1 |  |
| *Dendrocopos major* | *Dendrocopos syriacus* | 1,4 | 10 | 3 | 2 |
| *Colaptes auratus* | *Colaptes chrysoides* | 0,3 | 2 |  | 1 |
| *Celeus elegans* | *Celeus lugubris* | 0,25 | 1 | 2 |  |
| *Dryocopus lineatus* | *Dryocopus schultzei* | 0,2 | 1 |  |  |
| *Campephilus leucopogon* | *Campephilus melanoleucos* | 3,1 |  | 1 |  |
| *Picus canus* | *Picus viridis* | 2,5 | 8 | 2 |  |
| *Picus sharpei* | *Picus viridis* | 0,9 | 1 | 1 | 2 |
| *Dinopium benghalense* | *Dinopium psarodes* | NA | 2 |  |  |
| *Gecinulus grantia* | *Gecinulus viridis* | NA | 1 |  |  |

## LITERATURE

1. Aguillon S.M., Campagna L., Harrison R.G., Lovette I.J. 2018. A flicker of hope: genomic data distinguish northern flicker taxa despite low levels of divergence. Auk 135: 748–766.

2. Aguillon S.M., Walsh J., Lovette I.J. 2021. Extensive hybridization reveals multiple coloration genes underlying a complex plumage phenotype. Proceedings of the Royal Society B: Biological Sciences. 27: 288(1943):20201805.

3. Arnold M. L. 1992. Natural hybridization as an evolutionary process. Annual review of Ecology and Systematics 23: 237–261.

4. Alves Barcellos S., Kretschmer R., Santos de Souza M., Tura V., Pozzobon L. C., Ochotorena de Freitas T. R., Griffin D. K., O’Connor R., Gunski R. J., Del Valle Garnero A. 2024. Understanding microchromosomal organization and evolution in four representative woodpeckers (Picidae, Piciformes) through BAC-FISH analysis. Genome 67: 223–232.

5. Barrowclough G. F., Groth J. G., Bramlett E. K., Lai J. E. & Mauck W. M. 2018. Phylogeography and geographic variation in the Red-bellied Woodpecker (Melanerpes carolinus): characterization of mtDNA and plumage hybrid zones. The Wilson Journal of Ornithology 130: 671–683.

6. Barton N. H. 2001. The role of hybridization in evolution. Molecular ecology 2001: 551–568.

7. Benz B. W., Robbins M. B. & Peterson A. T. 2006. Evolutionary history of woodpeckers and allies (Aves: Picidae): placing key taxa on the phylogenetic tree. Molecular Phylogenetics and Evolution 40: 389–399.

8. Bertocchi N. A., de Oliveira T. D., del Valle Garnero A., Coan R. L. B., Gunski R. J., Martins C. & Torres F. P. 2018. Distribution of CR1-like transposable element in woodpeckers (Aves Piciformes): Z sex chromosomes can act as a refuge for transposable elements. Chromosome Research 26: 333–343.

9. Billerman S. M. & Carling M. D. 2017. Differences in aggressive responses do not contribute to shifts in a sapsucker hybrid zone. The Auk 134: 202–214.

10. Billerman S. M., Cicero C., Bowie R. C. K. & Carling M. D. 2019. Phenotypic and genetic introgression across a moving woodpecker hybrid zone. Molecular Ecology 28: 1692–1708.

11. Billerman S. M., Keeney B. K., Rodewald P. G., & Schulenberg T. S. 2022. Birds of the World. Cornell Laboratory of Ornithology, Ithaca, NY, USA.

12. BirdLife International and Handbook of the Birds of the World (2018) Bird species distribution maps of the world. Version 7.0. Available at https://datazone.birdlife.org/species/requestdis

13. Burrell A. S., Disotell T. R. & Bergey C. M. 2015. The use of museum specimens with high-throughput DNA sequencers. Journal of Human Evolution 79: 35–44.

14. Chialchia A. O. C. & Smith P. 2014. A notable hybrid woodpecker (Campephilus leucopogon x C. melanoleucus) (aves: picidae) from Paraguay. Ornitologia Neotropical 25: 459-464.

15. Christensen J. H. 1999. Gillelejespætten—Historien om Danmarks ‘’halve’’ hvidryggede spætte. DOF Nyt, 5: 8–9.

16. Cottenet G., Blancpain C., Chuah P. F. & Cavin C. 2020. Evaluation and application of a next generation sequencing approach for meat species identification. Food Control 110.

17. Cox B. 2001. The biogeographic regions reconsidered. Journal of Biogeography 28: 511–523.

18. de Oliveira T. D., Kretschmer R., Bertocchi N. A., Degrandi T. M., de Oliveira E. H. C., Cioffi M. dB., del Valle Garnero A & Gunski R. J. 2017. Genomic Organization of Repetitive DNA in Woodpeckers (Aves, Piciformes): Implications for Karyotype and ZW Sex Chromosome Differentiation. PLoS ONE 12: e0169987.

19. del Hoyo J., Elliot A. & Sargatal J. 2002. Handbook of the Birds of the World. Volume 7. Jacamars to Woodpeckers. eds. Lynx Edicions.

20. Dixon K.L. 1989. Contact Zones of Avian Congeners on the Southern Great Plains. The Condor 91: 15–22.

21. Edwards S. V., Kingan S. B., Calkins J. D., Balakrishnan C. N., Jennings W. B., Swanson W. J. & Sorenson M. D. 2005. Speciation in birds: genes, geography, and sexual selection. Proceedings of the National Academy of Sciences 102: 6550–6557.

22. Figarski T. 2017. Contrasting seasonal reactions of two sibling woodpeckers to playback stimulation in urban areas — implications for inventory and monitoring of the Syrian woodpecker. Behaviour 154: 981–996.

23. Figarski T. & Kajtoch Ł. 2018. Hybrids and mixed pairs of Syrian and great-spotted woodpeckers in urban populations. Journal of Ornithology 159: 311–314.

24. Fitzpatrick B. M. 2012. Estimating ancestry and heterozygosity of hybrids using molecular markers. BMC Evolutionary Biology 12: 1–14.

25. Flockhart D. T. T. & Wiebe K. L. 2009. Absence of Reproductive Consequences of Hybridization in the Northern Flicker (Colaptes auratus) Hybrid Zone. The Auk 126: 351–358.

26. Friedman V. S. 1993. Some observations on formation of a mixed pair of grey-headed woodpecker (Picus canus Gm.) and green woodpecker (P. viridis L.). Sbornik Trudov Zoologicheskogo Muzeya MGU 30: 183–196.

27. Friedman V. S. 2011. A riddle of the green woodpecker: how appear hybrids with the grey woodpecker? Berkut 20: 127–138.

28. Fuchs J., Ohlson J. I., Ericson P. G. P. & Pasquet E. 2006. Molecular phylogeny and biogeographic history of the piculets (Piciformes: Picumninae). Journal of Avian Biology 37: 487–496.

29. Fuchs J., Ohlson J. I., Ericson P. G. P. & Pasquet E. 2007. Synchronous intercontinental splits between assemblages of woodpeckers suggested by molecular data. Zoologica Scripta 36: 11–25.

30. Fuchs J. & Pons J-M. 2015. A new classification of the Pied Woodpeckers assemblage (Dendropicini, Picidae) based on a comprehensive multi-locus phylogeny. Moleclar Phylogenetics and Evolution 88: 28–37.

31. Fuchs J., Pons J-M., Liu L., Ericson P. G. P., Couloux A. & Pasquet E. 2013. A multi-locus phylogeny suggests an ancient hybridization event between Campephilus and melanerpine woodpeckers (Ayes: Picidae). Molecular Phylogenetics and Evolution 67: 578–588.

32. Gorman G. 1997. Hybridisation by Syrian Woodpeckers. British Birds 90: 578.

33. Gorman G. 1999. The Identification of Syrian Woodpecker. Alula 3: 82–88.

34. Gorman G. 2014. Woodpeckers of the World: The Complete Guide. A&C Black, Bloomsbury Publishing.

35. Gorman G. 2020. The Green Woodpecker: a monograph on *Picus viridis*. Picus press ISBN-9798676711870.

36. Grant P. R. & Grant B. R. 1992. Hybridization of bird species. Science 256: 193–197.

37. Grossen C., Seneviratne S. S., Croll D. & Irwin D. E. 2016. Strong reproductive isolation and narrow genomic tracts of differentiation among three woodpecker species in secondary contact. Molecular Ecology 25: 4247–4266.

38. Gurgul A., Miksza-Cybulska A., Szmatoła T., Semik-Gurgul E., Jasielczuk I., Bugno-Poniewierska M., Figarski T. & Kajtoch Ł. 2019. Evaluation of genotyping by sequencing for population genetics of sibling and hybridizing birds: An example using Syrian and Great Spotted Woodpeckers. Journal of Ornithology 160: 287–294.

39. Hammar R. 1970. The karyotypes of thirty-one birds. Hereditas 65: 29–58.

40. Holt B. G., Lessard, J. P., Borregaard M. K., Fritz S. A., Araújo M. B., Dimitrov, D., … & Rahbek C. 2013. An update of Wallace’s zoogeographic regions of the world. Science 339: 74–78.

41. Hogner S., Laskemoen T., Lifjeld J.T., Porkert J., Kleven O., Albayrak T., Kabasakal B., Johnsen A. 2012. Deep sympatric mitochondrial divergence without reproductive isolation in the common redstart *Phoenicurus phoenicurus*. Ecology and Evolution 2: 2974–88.

42. Hruska J. P. & Manthey J. D. 2021. De novo assembly of a chromosome-scale reference genome for the northern flicker Colaptes auratus. G3 Genes|Genomes|Genetics 11: jkaa026.

43. Hughes A. C., Orr M. C., Ma K., Costello M. J., Waller J., Provoost P & Qiao H. 2021. Sampling biases shape our view of the natural world. Ecography 44: 1259–1269.

44. Jenni L. & Müller H. 1983. Chromosomes of two species of woodpeckers (Aves: Piciformes). Experientia 39: 201–202.

45. Johnson N. K. & Johnson C. B. 1985. Speciation in Sapsuckers (sphyrapicus). II: Sympatry, hybridization, and mate preference in Sphyrapicus ruber daggetti and Sphyrapicus nuchalis. The Auk 102: 1–15.

46. Kajtoch Ł. & Kusal B. 2022. The first case of a successful brood from a double hybrid mixed pair (Dendrocopos syriacus x Dendrocopos major (Picidae)). IBIS 164: 1273–1277.

47. Kearns, A.M., Restani, M., Szabo, I. et al. 2018. Genomic evidence of speciation reversal in ravens. Nature Communications 9: 906.

48. King, B. 2007. Some 1960s additions to the list of Thailand’s birds. Natural History Bulletin of the Siam Society 55: 105–119.

49. Lee H. Y., Lee S. K. & Yu S. L. 1991. Karyotype of Korean birds, VI; Karyological analysis on two species of the genus dendrocopos by C-banding method. The Korean Journal of Zoology 33: 217–221.

50. Lanes-Quevedo A., Mastretta-Yanes A., Sanchez-Gonzalez L. A., Castillo-Chora V. J. & Navarro-Siguenza A. G. 2022. The tangled evolutionary history of a long-debated Mesoamerican taxon: The Velazquez Woodpecker (Melanerpes santacruzi, Aves: Picidae). Molecular Phylogenetics and Evolution 170.

51. Maddison W. P. & Knowles L. L. 2006. Inferring phylogeny despite incomplete lineage sorting. Systematic Biology 55: 21–30.

52. Maffre P., Chiang J. C. & Swanson-Hysell N. L. 2023. The effect of the Pliocene temperature pattern on silicate weathering and Pliocene–Pleistocene cooling. Climate of the Past 19: 1461–1479.

53. Manthey J. D., Boissinot S. & Moyle R. G. 2019. Biodiversity genomics of North American Dryobates. woodpeckers reveals little gene flow across the D. nuttallii x D. scalaris contact zone. The Auk 136.

54. Manthey J. D., Geiger M. & Moyle R. G. 2016. Relationships of morphological groups in the Northern Flicker superspecies complex (Colaptes auratus & C. chrysoides). Systematics and Biodiversity 15: 183–191.

55. Michalczuk J. 2014. Expansion of the Syrian Woodpecker Dendrocopos syriacus in Europe and Western Asia. Ornis Polonica 55: 149–161.

56. Michalczuk J., McDevitt A. D., Mazgajski T. D., Figarski T., Ilieva M., Bujoczek M., … & Kajtoch Ł. 2014. Tests of multiple molecular markers for the identification of Great Spotted and Syrian Woodpeckers and their hybrids. Journal of Ornithology 155: 591–600.

57. Moehring A. J. 2011. Heterozygosity and its Unexpected Correlations with Hybrid Sterility. Evolution 65: 2621–2630.

58. Natola L. & Burg T. M. 2018. Population Genetics and Speciation of Yellow-Bellied, Red-Naped, and Red-Breasted Sapsuckers (Sphyrapicus varius, S. nuchalis, and S. ruber). Journal of Heredity 109: 663–674.

59. Natola L., Curtis A., Hudon J. & Burg T. M. 2021. Introgression between Sphyrapicus nuchalis and S. varius sapsuckers in a hybrid zone in west-central Alberta. Journal of Avian Biology 52.

60. Natola L., Seneviratne S. S. & Irwin D. E. 2022. Population genomics of an emergent tri-species hybrid zone. Molecular ecology 3: 5356–5367.

61. Natola L., & Irwin D. 2023. Evidence for ancient selective sweeps followed by differentiation among three species of Sphyrapicus sapsuckers. bioRxiv preprint.

62. Olioso G. & Pons J. M. 2011. Variation of the plumage of Green Woodpecker in Languedoc-Roussillon (southern France). Ornithos 18: 73–83.

63. Oliveira U., Vasconcelos M. F. & Santos A. J. 2017. Biogeography of Amazon birds: rivers limit species composition, but not areas of endemism. Scientific Reports 7: 2992.

64. Ottenburghs J. 2018. Exploring the hybrid speciation continuum in birds. Ecology and Evolution 8: 13027–13034.

65. Ottenburghs J. 2023. How common is hybridization in birds? Journal of Ornithology 164: 913–920.

66. Ottenburghs J., Ydenberg R. C., Van Hooft P., Van Wieren S. E. & Prins H. H. T. 2015. The Avian Hybrids Project: gathering the scientific literature on avian hybridization. IBIS 157: 892–894.

67. Ottenburghs J. & Nicolaï M. P. 2024. Hybridization constrains the evolution of mimicry complexes in woodpeckers. Journal of Avian Biology 2024.

68. Pala I. & Coelho M. M. 2005. Contrasting views over a hybrid complex: between speciation and evolutionary “dead-end”. Gene 347: 283–294.

69. Pârâu L. G. & Wink M. 2021. Common patterns in the molecular phylogeography of western palearctic birds: a comprehensive review. Journal of Ornithology 162: 937–959.

70. Payseur B. A. & Rieseberg L. H. 2016. A genomic perspective on hybridization and speciation. Molecular Ecology 25: 2337–2360.

71. Pereira R. J., Monahan W. B. & Wake D. B. 2011. Predictors for reproductive isolation in a ring species complex following genetic and ecological divergence. BMC Evolutionary Biology 11: 1–15.

72. Perry J. R., Sumner S., Thompson C. & Hart A. G. 2021. ‘Citizen identification’: online learning supports highly accurate species identification for insect-focussed citizen science. Insect Conservation and Diversity 14: 862–867.

73. Petso T., Jamisola Jr R. S. & Mpoeleng D. 2022. Review on methods used for wildlife species and individual identification. European Journal of Wildlife Research 68.

74. Piett S., Hager H. A. & Gerrard C. 2015. Characteristics for evaluating the conservation value of species hybrids. Biodiversity and Conservation 24: 1931–1955.

75. Pons J-M., Olioso C., Curaud C. & Fuchs J, 2011. Phylogeography of the Eurasian green woodpecker (Picus viridis). Journal of Biogeography 38: 311–325.

76. Pons J-M., Masson C., Olioso G. & Fuchs J. 2019. Gene flow and genetic admixture across a secondary contact zone between two divergent lineages of the Eurasian Green Woodpecker Picus viridis. Journal of Ornithology 160: 935–945.

77. Ranasinghe R.W., Seneviratne S.S., Irwin D. 2024. Cryptic hybridisation dynamics in a three-way hybrid zone of *Dinopium* flamebacks on a tropical island. Ecology and Evolution 14: e70716.

78. Romanov M. N. 2022. Microchromosome organization in birds. MDPI, Basel, Switzerland Encyclopedia (online)

79. Rosqvist G. 1990. Quaternary glaciations in Africa. Quaternary Science Reviews 9: 281–297.

80. Roux C., Fraïsse C., Romiguier J., Anciaux Y., Galtier N., Bierne N. 2016. Shedding Light on the Grey Zone of Speciation along a Continuum of Genomic Divergence. PLoS Biology 14(12): e2000234.

81. Sampaio L., Aleixo A., Schneider H., et al. 2018. Molecular and plumage analyses indicate the incomplete separation of two woodpeckers (Aves, Picidae), Zoologica Scripta 47: 418–427.

82. Sætre G. P. 2013. Hybridization is important in evolution, but is speciation?. Journal of Evolutionary Biology 26: 256–258.

83. Schwenk K., Brede N. & Streit B. 2008. Introduction. Extent, processes and evolutionary impact of interspecific hybridization in animals. Philosophical Transactions of the Royal Society B: Biological Sciences 363: 2805–2811.

84. Seneviratne S. S., Davidson P., Martin K. & Irwin D. E. 2016. Low levels of hybridization across two contact zones among three species of woodpeckers (Sphyrapicus sapsuckers). Journal of Avian Biology 47: 743–910.

85. Seneviratne S. S., Toews D. P. L., Brelsford A. & Irwin D. E. 2012. Concordance of genetic and phenotypic characters across a sapsucker hybrid zone. Journal of Avian Biology 43: 119–130.

86. Shakya S. B., Fuchs J., Pons, J. M. & Sheldon F. H. 2017. Tapping the woodpecker tree for evolutionary insight. Molecular Phylogenetics and Evolution 116: 182–191.

87. Shields G. F. 1982. Comparative Avian Cytogenetics: A Review. The Condor 84: 45–58.

88. Shields G. F., Jarrell G. H. & Redrupp E. 1982. Enlarged Sex Chromosomes of Woodpeckers (piciformes). The Auk 99: 767–771.

89. Shields M. A. 1988. A red shafted x yellow shafted flicker intergrade in Carteret County, N.C. The Chat (Raleigh) 52: 78–79.

90. Short L. L. 1982. Woodpeckers of the World. Delaware Museum of Natural History, USA

91. Short L. L. & Morony J. J. 1970. A second hybrid Williamsons x Red-Naped Sapsucker and an evolutionary history of sapsuckers. The Condor 72: 310–315.

92. Smith B. T., McCormack J. E., Cuervo A. M., … & Brumfield R. T. 2014. The drivers of tropical speciation. Nature 515: 406–409.

93. Walters E. L., Robles H., Czeszczewik D., Perktas U. & Pasinelli G. 2020. Conservation and ecology of woodpeckers. Foreword to the 8th International Woodpecker Conference Proceedings. Acta Ornithologica 55: 61–62.

94. Whittaker J.E. & Horne D.J. 2009. Pleistocene. In: Whittaker J.E. & Hart M.B (eds). Ostracods in British Stratigraphy. Geological Society of London, Micropaleontological Society.

95. Wiley G. & Miller M. J. 2020. A highly contiguous genome for the Golden-Fronted Woodpecker (*Melanerpes aurifrons*) via Hybrid Oxford Nanopore and short read assembly. G3 (Bethesda) 10: 1829–1836.

96. Winkler H., Gamauf A., Nittinger F. & Haring E. 2014. Relationships of Old World woodpeckers (Aves: Picidae)—new insights and taxonomic implications. Annalen des Naturhistorischen Museums in Wien. Serie B für Botanik und Zoologie 116: 69–86.

97. Xiao-Zhuang B. & Qing-wei L. 1989. Studies on The Karyotypes of Birds Ⅴ.The 20 species of Climber birds.(Aves). Zoological Research 10: 309–317.

98. Zachos F. E. 2016. Species Concepts in Biology Historical Development, Theoretical Foundations and Practical Relevance. Springer Cham, Springer International Publishing Switzerland.

## Literature

100. Aulén G. 1979. En hybrid mellan större - och vitryggig hackspett i Uppsala. Fåglar i Uppsala 6: 27–32.

101. Billerman S. M. & Carling M. D. 2017. Differences in aggressive responses do not contribute to shifts in a sapsucker hybrid zone. The Auk 134: 202–214.

102. Billerman S. M., Cicero C., Bowie R. C. K. & Carling M. D. 2019. Phenotypic and genetic introgression across a moving woodpecker hybrid zone. Molecular Ecology 28: 1692–1708.

103. Billerman S. M., Keeney B. K., Rodewald P. G., & Schulenberg T. S. 2022. Birds of the World. Cornell Laboratory of Ornithology, Ithaca, NY, USA.

104. Billerman S. M., Murphy M. A. & Carling M. D. 2016. Changing climate mediates Sapsucker (Aves:Sphyrapicus) hybrid zone movement. Ecology and Evolution 6: 7976–7990.

105. Bird D. & Suedbeck P. 2004. First record of hybrid between green- and grey-headed woodpecker Picus viridis x P. canus in Sachsen-Anhalt. Notatki Ornitologiczne 44: 273-275.

106. Chialchia A. O. C. & Smith P. 2014. A notable hybrid woodpecker (Campephilus leucopogon x C. melanoleucus) (aves: picidae) from Paraguay. Ornitologia Neotropical 25: 459-464.

107. Christensen J. H. 1999. Historien om Danmarks “halve” Hvidryggede. Dansk Ornithologisk Forenings Tidsskrift 4: 8–9.

108. Czechowski P. & Bocheński M. 2012. A case of hybridization between the Green Woodpecker Picus viridis and the Grey-faced Woodpecker Picus canus. Przegląd Przyrodniczy 23: 72–74.

109. Del Hoyo J., Elliot A. & Sargatal J. 2002. Handbook of the Birds of the World. Volume 7. Jacamars to Woodpeckers. eds. Lynx Edicions.

110. Dernfalk C. 1983. Hybrid mellan större hackspett och vitryggig hackspett i Närke. Fåglar i Närke 6: 11–14.

111. Dixon K. L. 1989. Contact Zones of Avian Congeners on the Southern Great Plains. The Condor 91: 15–22.

112. Figarski T. & Kajtoch Ł. 2018. Hybrids and mixed pairs of Syrian and great-spotted woodpeckers in urban populations. Journal of Ornithology 159: 311–314.

113. Freed L. A., Warakagoda D., Cann R. L., Sirivardana U. & Hettige U. 2015. A hybrid swarm of Dinopium woodpeckers in Sri Lanka. Wilson Journal of Ornithology 127: 13–20.

114. Friedman V. S. 1993. Some observations on formation of a mixed pair of grey-headed woodpecker (Picus canus Gm.) and green woodpecker (P. viridis L.). Sbornik Trudov Zoologicheskogo Muzeya MGU 30: 183–196.

115. Friedmann V. S. 2011. A riddle of the green woodpecker: how appear hybrids with the grey woodpecker? Berkut 20: 127–138.

116. Fuchs J. & Pons J-M. 2015. A new classification of the Pied Woodpeckers assemblage (Dendropicini, Picidae) based on a comprehensive multi-locus phylogeny. Moleclar Phylogenetics and Evolution 88: 28–37.

117. Gaël R. 2018. Birds of Rwanda: A checklist by Gaël R. Vande weghe.

118. Gorman G. 1997. Hybridisation by Syrian Woodpeckers. British Birds 90: 578.

119. Grossen C., Seneviratne S. S., Croll D. & Irwin D. E. 2016. Strong reproductive isolation and narrow genomic tracts of differentiation among three woodpecker species in secondary contact. Molecular Ecology 25: 4247–4266.

120. Gurgul A., Miksza-Cybulska A., Szmatoła T., Semik-Gurgul E., Jasielczuk I., Bugno-Poniewierska M., Figarski T. & Kajtoch Ł. 2019. Evaluation of genotyping by sequencing for population genetics of sibling and hybridizing birds: An example using Syrian and Great Spotted Woodpeckers. Journal of Ornithology 160: 287–294.

121. Ivanchev V. P. 1993. A case of hybridization of woodpeckers of genus Picus. Sbornik Trudow Zoologicheskogo Muzeya MGU 30: 197–200.

122. Johnson N. K. 1969. [Review of] Three papers on variation in flickers (Colaptes) by Lester L. Short, Jr. Wilson Bulletin 81: 225–230.

123. Johnson N. K. & Johnson C. B. 1985. Speciation in Sapsuckers (sphyrapicus). II: Sympatry, hybridization, and mate preference in Sphyrapicus ruber daggetti and Sphyrapicus nuchalis. The Auk 102: 1–15.

124. Kajtoch Ł. & Kusal B. 2022. The first case of a successful brood from a double hybrid mixed pair (Dendrocopos syriacus x Dendrocopos major (Picidae)). IBIS 164: 1273–1277.

125. King, B. 2007. Some 1960s additions to the list of Thailand’s birds. Natural History Bulletin of the Siam Society 55: 105–119.

126. Kratter A. W., Carreno M. D., Chesser R. T., O’Neill J. P. & Sillett T. S. 1992. Further notes on bird distribution in northeastern Dpto. Santa Cruz, Bolivia, with two species new to Bolivia. Bulletin of the British Ornithologists’ Club 112: 143–150.

127. Ławicki Ł., Cofta T., Beuch Sz., Dmoch A., Sikora A., Aftyka S., Czechowski P., Bocheński M., Sieczak K. & Mazgaj Sz. 2015. Identification and occurrence of hybrids Grey-headed x European Green Woodpecker in Poland. Dutch Birding 37: 215–228.

128. Llanes-Quevedo A., Mastretta-Yanes A., Sanchez-Gonzalez L. A., Castillo-Chora V. J. & Navarro-Siguenza A. G. 2022. The tangled evolutionary history of a long-debated Mesoamerican taxon: The Velazquez Woodpecker (Melanerpes santacruzi, Aves: Picidae). Molecular Phylogenetics and Evolution 170.

129. Manthey J. D., Boissinot S. & Moyle R. G. 2019. Biodiversity genomics of North American Dryobates. woodpeckers reveals little gene flow across the D. nuttallii x D. scalaris contact zone. The Auk 136.

130. Manthey J. D., Geiger M. & Moyle R. G. 2016. Relationships of morphological groups in the Northern Flicker superspecies complex (Colaptes auratus & C. chrysoides). Systematics and Biodiversity 15: 183–191.

131. McCarthy E. M. 2006. Handbook of Avian Hybrids of the World. Oxford University Press.

132. Michalczuk J. 2014. Expansion of the Syrian Woodpecker Dendrocopos syriacus in Europe and Western Asia. Ornis Polonica 55: 149–161.

133. Michalczuk J., McDevitt A. D., Mazgajski T. D., Figarski T., Ilieva M., Bujoczek M., … & Kajtoch Ł. 2014. Tests of multiple molecular markers for the identification of Great Spotted and Syrian Woodpeckers and their hybrids. Journal of Ornithology 155: 591–600.

134. Miller A. H. 1955. A hybrid woodpecker and its significance in speciation in the genus Dendrocopos. Evolution 9.

135. Mlodinow S. G., Leukering T., Brooks T. & Moore N. 2015. Apparent hybrid Downy Woodpecker x Hairy Woodpecker in Colorado. Western Birds 46: 2–7.

136. Natola E. M. 2017. Historic and Contemporary Hybridization of Three Sphyrapicus Sapsucker Species. Collections Arts and Science, Faculty of University of Lethbridge Theses.

137. Natola L., Curtis A., Hudon J. & Burg T. M. 2021. Introgression between Sphyrapicus nuchalis and S. varius sapsuckers in a hybrid zone in west-central Alberta. Journal of Avian Biology 52.

138. Natola L., Seneviratne S. S. & Irwin D. E. 2022. Population genomics of an emergent tri species hybrid zone. Molecular ecology 3: 5356–5367.

139. Natola L., Billerman S. M., Carling M. D., Seneviratne S. S. & Irwin D. 2023. Geographic variability of hybridization between Red-Breasted and Red-Naped Sapsuckers. Evolution 77: 580–592.

140. Neto F. L. 1995. Um hibrido entre Picumnus cirratus temminckii e P. albosquamatus guttifer (Piciformes: Picidae). Ararajuba 3: 68–69.

141. Oblehorser H. C. 1930. Notes on a collection of birds from Arizona and New Mexico. Sci. Publ. Cleveland Museum of Natural History 1: 83–124.

142. Olioso G. & Pons J. M. 2011. Variation of the plumage of Green Woodpecker in Languedoc-Roussillon (southern France). Ornithos 18: 73–83.

143. Ottenburghs J. & Nicolaï M. P. 2024. Hybridization constrains the evolution of mimicry complexes in woodpeckers. Journal of Avian Biology 2024.

144. Pons J-M., Olioso C., Curaud C. & Fuchs J, 2011. Phylogeography of the Eurasian green woodpecker (Picus viridis). Journal of Biogeography 38: 311–325.

145. Pons J-M., Masson C., Olioso G. & Fuchs J. 2019. Gene flow and genetic admixture across a secondary contact zone between two divergent lineages of the Eurasian Green Woodpecker Picus viridis. Journal of Ornithology 160: 935–945.

146. Prigogine A. 1987. Hybridization between the megasubspecies cailliautii and permista of the green-backed woodpecker, Campethera cailliautii. Gerfaut 77: 187–204.

147. Prigogine A., & Louette M. 1983. Contacts secondaires entre les taxons appartenant à la super-espèce Dendropicos goertae. Gerfaut 73: 9–83.

148. Ranasinghe R.W., Seneviratne S.S., Irwin D. 2024. Cryptic hybridization dynamics in a three-way hybrid zone of *Dinopium* flamebacks on a tropical island. Ecology and Evolution 14: e70716.

149. Ridgway R. 1914. The birds of North and Middle America. Bull. U.S. Natl. Mus. 50, Part 6

150. Round P.D., Hobday J.M., Kanjanavanit R. & Steward J. S. 2012. A nesting pair of Gecinulus woodpeckers in a likely zone of intergradation between Pale-headed Woodpecker G. grantia and Bamboo Woodpecker G. viridis. Forktail 28: 113–120.

151. Saminda F. P., Darren I. E. & Sampath S. S. 2016. Phenotypic and genetic analysis support distinct species status of the Red-backed Woodpecker (Lesser Sri Lanka Flameback: Dinopium psarodes) of Sri Lanka. The Auk 133: 497–511.

152. Sarkanen S. & Koivusaari J. 2001. Valkoselkätikan esiintymisestä Merenkurkun alueella. Linnut 36: 34–36.

153. Selander R. K. & Giller D. R. 1963. Species Limits in the Woodpecker Genus Centurus (Aves). Bulletin of the American Museum of Natural History 124: 216–273.

154. Seneviratne S. S., Davidson P., Martin K. & Irwin D. E. 2016. Low levels of hybridization across two contact zones among three species of woodpeckers (Sphyrapicus sapsuckers). Journal of Avian Biology 47: 743–910.

155. Seneviratne S. S., Toews D. P. L., Brelsford A. & Irwin D. E. 2012. Concordance of genetic and phenotypic characters across a sapsucker hybrid zone. Journal of Avian Biology 43: 119–130.

156. Sexton C. W. 1986. A possible hybrid Ladder-backed × Downy woodpecker in central Texas. Bulletin of the Texas Ornithological Society, 19: 2–5.

157. Shakya S. B., Fuchs J., Pons, J. M. & Sheldon F. H. 2017. Tapping the woodpecker tree for evolutionary insight. Molecular Phylogenetics and Evolution 116: 182–191.

158. Short L. L. 1971. Systematics and behavior of some North American woodpeckers, genus Picoides (Aves). Bull. Am. Mus. Nat. Hist. 145: 1–118.

159. Short L. L. 1972. Relationships among the four species of the superspecies Celeus elegans (Aves, Picidae). American Museum Novitates No. 2487

160. Short L. L. 1982. Woodpeckers of the World. Delaware Museum of Natural History, USA

161. Short L. L. & Morony J. J. 1970. A second hybrid Williamsons x Red-Naped Sapsucker and an evolutionary history of sapsuckers. The Condor 72: 310–315.

162. Smith J. I. 1987. Evidence of hybridization between Red-Bellied and Golden-Fronted Woodpeckers. The Condor 89: 377–386.

163. Stiles G. & Skutch A. 1989. A Guide to the Birds of Costa Rica. Cornell University Press, Ithaca, NY.

164. Trombino C. L. 1998. Species interactions in the hybrid zone between Red-Breasted (Sphyrapicus ruber) and Red-Naped (Sphyrapicus nuchalis) Sapsuckers: Fitness consequences, reproductive character displacement, and nest site selection. Northern Illinois University ProQuest Dissertations & Theses.

165. Unitt P. 1986. Another hybrid downy x Nuttall’s woodpecker from San Diego County. Western Birds 17: 43–44.

166. Vaurie C. 1959. Systematic notes on Palearctic birds, no. 35. Picidae: The genus Dendrocopos (Part 1). American Museum Novitates, no. 1946.

167. Winkler H., Christie D.A. & Nurney D. 1995. Woodpeckers: A Guide to the Woodpeckers of the World. Houghton Mifflin, Boston, New York.

## Literature

169. Aguillon S. M., Campagna L., Harrison R. G. & Lovette I. J. 2018. A flicker of hope: Genomic data distinguish Northern Flicker taxa despite low levels of divergence. AUK 135: 748–766.

170. Aguillon S. M. & Rohwer V. G. 2022. Revisiting a classic hybrid zone: Movement of the northern flicker hybrid zone in contemporary times. Evolution 76: 1082–1090.

171. Anderson B. W. 1971. Man’s Influence on Hybridization in Two Avian Species in South Dakota. Condor 73: 342–347.

172. Barrowclough G. F., Groth J. G., Bramlett E. K., Lai J. E. & Mauck W. M. 2018. Phylogeography and geographic variation in the Red-bellied Woodpecker (Melanerpes carolinus): characterization of mtDNA and plumage hybrid zones. The Wilson Journal of Ornithology 130: 671–683.

173. Bock C. E. 1971. Pairing in Hybrid Flicker Populations in Eastern Colorado. AUK 88: 921–924.

174. Flockhart D. T. T. & Wiebe K. L. 2007. The role of weather and migration in assortative pairing within the northern flicker (Colaptes auratus) hybrid zone. Evolutionary Ecology Research 9: 887–903.

175. Flockhart D. T. T. & Wiebe K. L. 2008. Variable Weather Patterns Affect Annual Survival of Northern Flickers More than Phenotype in the Hybrid Zone. Condor 110: 701–708.

176. Flockhart D. T. T. & Wiebe K. L. 2009. Absence of Reproductive Consequences of Hybridization in the Northern Flicker (Colaptes Auratus) Hybrid Zone. AUK 126: 351–358.

177. Hudon J., Wiebe K. L. & Stradi, R. 2021. Disruptions of feather carotenoid pigmentation in a subset of hybrid northern flickers (Colaptes auratus) may be linked to genetic incompatibilities. Comparative Biochemistry and Physiology B-Biochemistry & Molecular Biology 251.

178. McGillivary B. W. & Biermann G. C. 1987. Expansion of the Zone of Hybridization of Northern Flickers in Alberta. The Wilson Bulletin 99: 690–692.

179. Moore W. S. 1987. Random Mating in the Northern Flicker Hybrid Zone - Implications for the Evolution of Bright and Contrasting Plumage Patterns in Birds. Evolution 41: 539–546.

180. Moore W. S. & Buchanan D. B. 1985. Stability of the Northern Flicker hybrid zone in historical times: implications for adaptive speciation theory. Evolution 39: 135–151.

181. Moore W. S., Graham J. H. & Price J. T. 1991. Mitochondrial DNA Variation in the Northern Flicker (Colaptes-auratus, Aves). Molecular Biology and Evolution 8: 327.

182. Moore, W. S., and W. D. Koenig. 1986. Comparative reproductive success of yellow-shafted, red-shafted, and hybrid flickers across a hybrid zone. AUK 103: 42–51.

183. Prigogine A. 1987. Hybridization between the megasubspecies cailliautii and permista of the green-backed woodpecker, Campethera cailliautii. Gerfaut 77: 187–204.

184. Rieseberg L. H., Church S. A. & Morjan C. L. 2004. Integration of populations and differentiation of species. New Phytologist 161: 59–69.

185. Short L. L. 1965. Hybridization in the flickers (Colaptes) of North America. Bulletin of the American Museum of Natural History 129 :307–428.

186. Short L. L. 1972. Hybridization, taxonomy and avian evolution. Annals of the Missouri Botanical Garden 3: 447–453.

187. Short L.L. 1982. Woodpeckers of the World. Delaware Museum of Natural History, USA

188. Wiebe K. L. 2000. Assortative mating by color in a population of hybrid Northern Flickers. AUK 117: 525–529.

189. Wiebe K. L. & Bortolotti G. R. 2001. Variation in colour within a population of northern flickers: a new perspective on an old hybrid zone. Canadian Journal of Zoology 79: 1046–1052.

190. Wiebe K. L. & Moore W. S. 2008. Northern Flicker (Colaptes auratus). In A. Poole [ED.], The birds of North America online, No. 166a. Cornell Lab Ornithology, Ithaca, NY.

## Literature

192. Desfayes M. 1969. A possible hybrid Jynx ruficollis x torquilla. Bulletin of the British Ornithologists’ Club 89: 110–112.

193. Goodwin D. 1968. Notes on woodpeckers (Picidae). Bulletin of the British Museum (Natural History) Zoology 17: 1–44.

194. Gorman G. 2022. The Wryneck. Pelagic Publishing, Exeter, Uk.

195. McCarthy E. M. 2006. Handbook of Avian Hybrids of the World. Oxford University Press.

196. Ottenburghs J. & Nicolaï M. P. 2024. Hybridization constrains the evolution of mimicry complexes in woodpeckers. Journal of Avian Biology 2024.

197. Philips W. W. A. 1953. Nests and eggs of Ceylon birds (Picidae and Capitonidae). Ceylon Journal of Science 25: 35–45.

198. Short L.L. 1982. Woodpeckers of the World. Delaware Museum of Natural History, USA

199. Short L.L. & Bock W.J. 1972. Possible hybrid Jynx is an aberrant Jynx ruficollis. Bull. Brit. Orn. Club. 92: 28–31.

200. Winkler H., Christie D.A. & Nurney D. 1995. Woodpeckers: A Guide to the Woodpeckers of the World. Houghton Mifflin, Boston, New York.

